# Deep Learning for RNA Secondary Structure Determination: Gauging Generalizability and Broadening the Scope of Traditional Methods

**DOI:** 10.1101/2025.11.04.686644

**Authors:** Marcell Szikszai, Ting-Yuan Wang, Ryan Krueger, David H. Mathews, Max Ward, Sharon Aviran

## Abstract

The diverse regulatory functions, protein production capacity, and stability of natural and synthetic RNAs are closely tied to their ability to fold into intricate structures. Determining RNA structure is thus fundamental to RNA biology and bioengineering. Among existing approaches to structure determination, computational secondary structure prediction offers a rapid and low-cost strategy and is thus widely used, especially when seeking to identify functional RNA elements in large transcriptomes or screen massive libraries of novel designs. While traditional approaches rely on detailed measurements of folding energetics and/or probabilistic modeling of structural data, recent years have witnessed a surge in deep learning methods, inspired by their tremendous success in protein structure prediction. However, the limited diversity and volume of known RNA structures can impede their ability to accurately predict structures markedly different from the ones they have seen. This is known as the generalization gap and currently poses a major barrier to progress in the field. In this Perspective article, we gauge method generalizability using a new benchmark dataset of structured RNAs we curated from the Protein Data Bank. We also discuss the emergence of deep learning methods for predicting structure probing data and use a new dataset to underscore generalization challenges unique to this domain along with directions for future improvement. Expanding beyond improving predictive accuracy, we review how advances in deep learning have recently enabled scalable and accessible optimization of traditional structure prediction methods and their seamless integration with modern neural networks.

## INTRODUCTION

Computationally determining the secondary structure of RNA has been of interest to researchers since the 1970s, when thermodynamic rules were devised for defining optimal structures (Tinoco et al., 1971) and when (Nussinov et al., 1978) showed a dynamic programming algorithm for computing an RNA secondary structure with maximal number of base pairs. Since then, dynamic programming has been used extensively by approaches incorporating both thermodynamics (Anfinsen, 1973; Waterman & Smith, 1978; Zuker & Stiegler, 1981; Lu et al., 2009) and statistical modeling (Dowell & Eddy, 2004; Do et al., 2006; Rivas et al., 2012; Sakakibara et al., 1994) to improve the accuracy of predicted structures. Today, the most accurate of these are able to correctly predict about 70% of known base pairs (Mathews et al., 2004a; Lorenz et al., 2011), and are thought to generalize well to novel structures.

Since the revived interest in neural networks in the early 2010s (Krizhevsky et al., 2012), several methods have attempted to apply deep learning to RNA structure prediction (see reviews by (Chaturvedi et al., 2025; Zablocki et al., 2025)). This interest was further accelerated with the advent of tools like AlphaFold (Senior et al., 2020; Jumper et al., 2021), which are able to predict the 3D structure of proteins with high accuracy, and have even attempted to predict the 3D structure of RNAs (Abramson et al., 2024), although with much less success (Das et al., 2023; Schneider et al., 2023). RNA is generally accepted to first form secondary structures (2D) before it forms tertiary structures (3D) (Tinoco & Bustamante, 1999), so secondary structure prediction is considered very important. Despite many attempts to solve secondary structure prediction via deep learning, several papers have questioned the generalizability of these models, i.e. how well they perform on sequences not related by homology to those in the training set (Szikszai et al., 2022; Flamm et al., 2022; Justyna et al., 2023). It was shown that achieving high intra-family performance (accuracy when the training on testing sequences that are homologous) is near-trivial with deep learning, but inter-family generalization to RNAs outside the set of sequences homologous to those in the training set is difficult. Here we assemble a new dataset of secondary structures that is outside the training sets for deep learning, which we call Archive-NoFam. Using this dataset we find that deep learning-based methods do not generalize better than thermodynamic methods.

With growing evidence of generalizability challenges, interest has emerged in learning the principles of RNA folding from another data source, namely, structure probing assays. While probing data do not provide explicit, high-resolution information on secondary structures, they have broadly informed both computational structure prediction and a plethora of structure-function studies were performed at transcriptome-scale and under diverse cellular conditions (Aviran & Incarnato, 2022; Wang et al., 2021). This motivated recent efforts to model these data by deep learning as well as design massive synthetic libraries to be probed and used as model train and test data (Boyd et al., 2023; He et al., 2024). Here, we review this recent progress and report an exploratory performance analysis using a dataset we curated to gauge generalizability. We also discuss the broader utility of such models and train and test issues unique to them.

A third theme of this article is the renewed interest in traditional structure prediction methods, motivated by new capabilities for their rapid and flexible optimization, which directly build off of software and hardware infrastructure developed for deep learning. In that context, we review recent applications of a novel differential folding framework to gradient-based model optimization and explain how these advances enable a vision of platforms that fuse together traditional and modern models and jointly learn them from multimodal data.

## RESULTS

### Testing Models for Secondary Structure Prediction

Methods for predicting the secondary structure of RNAs can be placed into two broad categories: single-sequence and alignment/homology based. While deep learning 3D structure prediction– both for proteins and RNAs–makes widespread use of multiple sequence alignments (MSAs) (Abramson et al., 2024), this is not the case for RNA secondary structure. Most methods are single-sequence, and do not make direct use of homology or alignments. We attribute this to two main factors: 1) many of the known (i.e. ground truth) secondary structures are themselves derived from alignment/homology based methods, and 2) a major strength of alignment and homology based methods is that they are not “black-boxes” like deep learning models, and can be interpreted or even incrementally improved by experts. This means naive performance improvements by deep learning tools have minimal impact on their relevance. As a result, we focus on single-sequence methods.

Most tasks where deep learning excels are in domains where a large quantity of *diverse* data is available. The majority of known RNA secondary structures are catalogued by Rfam, the largest database of ncRNA families, i.e. sets of homologous sequences. As of today, Rfam 15.0 contains 4,178 families, each with a consensus secondary structure (Ontiveros-Palacios et al., 2025). While certainly not exhaustive, it is easy to argue that this is fairly representative of the number of known unique secondary structures. It should also be noted that the quality of these families can vary, with some comprised of deep alignments and consensus structures supported by strong experimental evidence (e.g. such as tRNA [<u>RF00005</u>]), but some families comprising of only shallow alignments, sometimes as low as two sequences, and low-confidence structures that have not been confirmed experimentally, but rather predicted with tools like RNAalifold (Bernhart et al., 2008), e.g. small nucleolar RNA snoR03 (<u>RF01584</u>).

The secondary structure of ncRNAs is strongly conserved, much more so than sequence (Pace et al., 1999). This means that two ncRNAs from the same family, even if they have relatively low sequence similarity, will have matching secondary structures in that there is a core set of recognizably homologous stems and loops. Beyond these Rfam families, there exist a small number of experimentally determined structures for synthetic RNAs (for example, molecules designed to fold into a specific secondary structure), but to our knowledge, no publicly available database exists that comprehensively curates these. As a result, with very few exceptions, nearly all deep learning models for secondary structure prediction are trained on Rfam, or more accurately, subsets of Rfam. The two most commonly used datasets, ArchiveII (Sloma & Mathews, 2016) and bpRNA-1m (Danaee et al., 2018), contain subsets of the families from Rfam, with the only exception being a few synthetic constructs in bpRNA-1m from the RCSB Protein Data Bank (PDB) (Berman et al., 2000). See Tables 1 and 2.

**Table 1.**
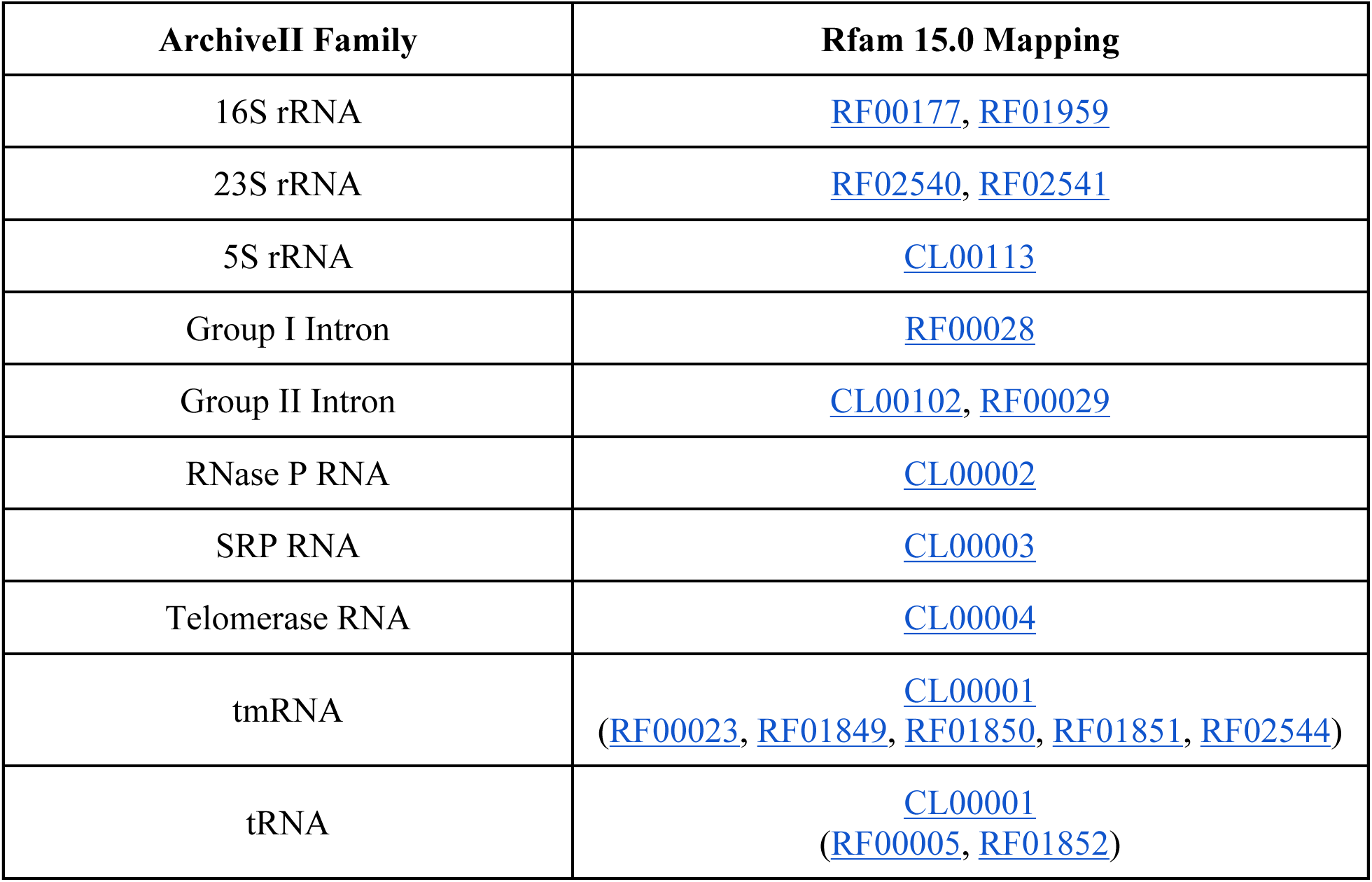
Mapping from ArchiveII families to Rfam 15.0 clans and families. Note that tRNA and tmRNA share a clan (<u>CL00001</u>) in Rfam 15.0.

**Table 2.** Mapping from bpRNA-1m sources to Rfam families. Note that while no naive mapping exists between the bpRNA-1m PDB sequences and Rfam 15.0, our analysis finds significant overlap between the two.

| **bpRNA-1m source** | **Citation** | **Rfam 15.0 mapping** |
| --- | --- | --- |
| The Comparative RNA Web (CRW) Site | (Cannone et al., 2002) | CL00111 |
| tmRNA Database | (Zwieb et al., 2003) | CL00001<br>(RF00023, RF01849,<br>RF01850, RF01851,<br>RF02544) |
| Signal Recognition Particle Database | (Rosenblad et al., 2003) | CL00003 |
| Sprinzl tRNA Database (tRNAdb) | (Jühling et al., 2009) | CL00001<br>(RF00005, RF01852) |
| The RNase P Database | (Brown, 1998) | CL00002 |
| The RNA Family Database | (Nawrocki et al., 2015) | Rfam 12.2 |
| RCSB Protein Data Bank | (Berman et al., 2000) | N/A |

An important strength of Rfam is that families are well-defined by Infernal covariance models (Nawrocki et al., 2009). A covariance model is a profile stochastic context-free grammar, based on an MSA (called a *seed alignment*) and a consensus secondary structure per family. This means that ncRNAs with structural homology to Rfam can easily be identified with Infernal and assigned an expectation value that a random sequence will score at least as well as the query sequence in terms of homology to a particular family. While there is no obvious direct mapping of the PDB structures used by bpRNA-1m to Rfam available, we have performed a homology search to create this mapping using Infernal (Nawrocki & Eddy, 2013). We find 346 sequence-structure pairs in this set (just 0.34% of the total bpRNA-1m dataset) that do not show homology to Rfam at an E-value threshold of 0.05. Of these, 273 are shorter than 32 nucleotides, which makes finding homology to Rfam families difficult, leaving only 73 structures that are unlikely to be included in Rfam.

Perhaps unsurprisingly, given the limited diversity in the available data (after all, a dataset based on Rfam can only contain 4,178 “relatively unique” secondary structures), several papers have pointed out significant issues with generalization to structures outside of the training sets for many models. The first of these papers by (Rivas et al., 2012) investigated discriminative statistical methods and probabilistic models, and showed that a lack of diversity in the training set causes issues with generalization. Later, papers by (Szikszai et al., 2022) and (Flamm et al., 2022) extended this work to show similar issues for deep learning models, bringing into question the practical utility of the existing models at the time.

In fact, the practical utility of single-sequence models, especially based on deep learning, must be carefully examined within the context of generalization. In cases where there is homology to an existing Rfam family, there is little reason to use single-sequence models over tools like Infernal and CaCoFold (Rivas, 2020). Not only are these tools extremely accurate, they also provide further insights about structural conservation. The missing piece of the puzzle is then novel families and synthetic RNAs. Naturally, accurate structure prediction of novel families is extremely useful. Additionally, inverse folding relies on accurate folding to determine the RNA structures and sequences, that are out of distribution of natural ncRNAs (Ward et al., 2023), and mRNA open reading frame design often takes secondary structure into consideration to design more structured RNAs which are found to be more stable against degradation (Mauger et al., 2019; Wayment-Steele et al., 2021; Zhang et al., 2023). Crucially, this means that the utility of single-sequence deep learning models for novel ncRNAs or synthetic RNAs is extremely high, if they are able to provide more accurate structure than traditional methods.

To address the practical utility of deep learning models, we curated a dataset of 89 experimentally determined secondary structures across 47 categories with no structural homology to Rfam at an E-value threshold of 1.0^1^. See Supplemental File 1 for details of all 3D structures. Using this dataset, called Archive-NoFam, we benchmarked five highly cited or relevant deep learning methods for secondary structure prediction. See Materials and Methods for details about dataset curation and method selection. See Fig. 1 for a breakdown of our results and Supplemental Tables S1-S7 for detailed performance analysis for each target.

**Figure 1.**
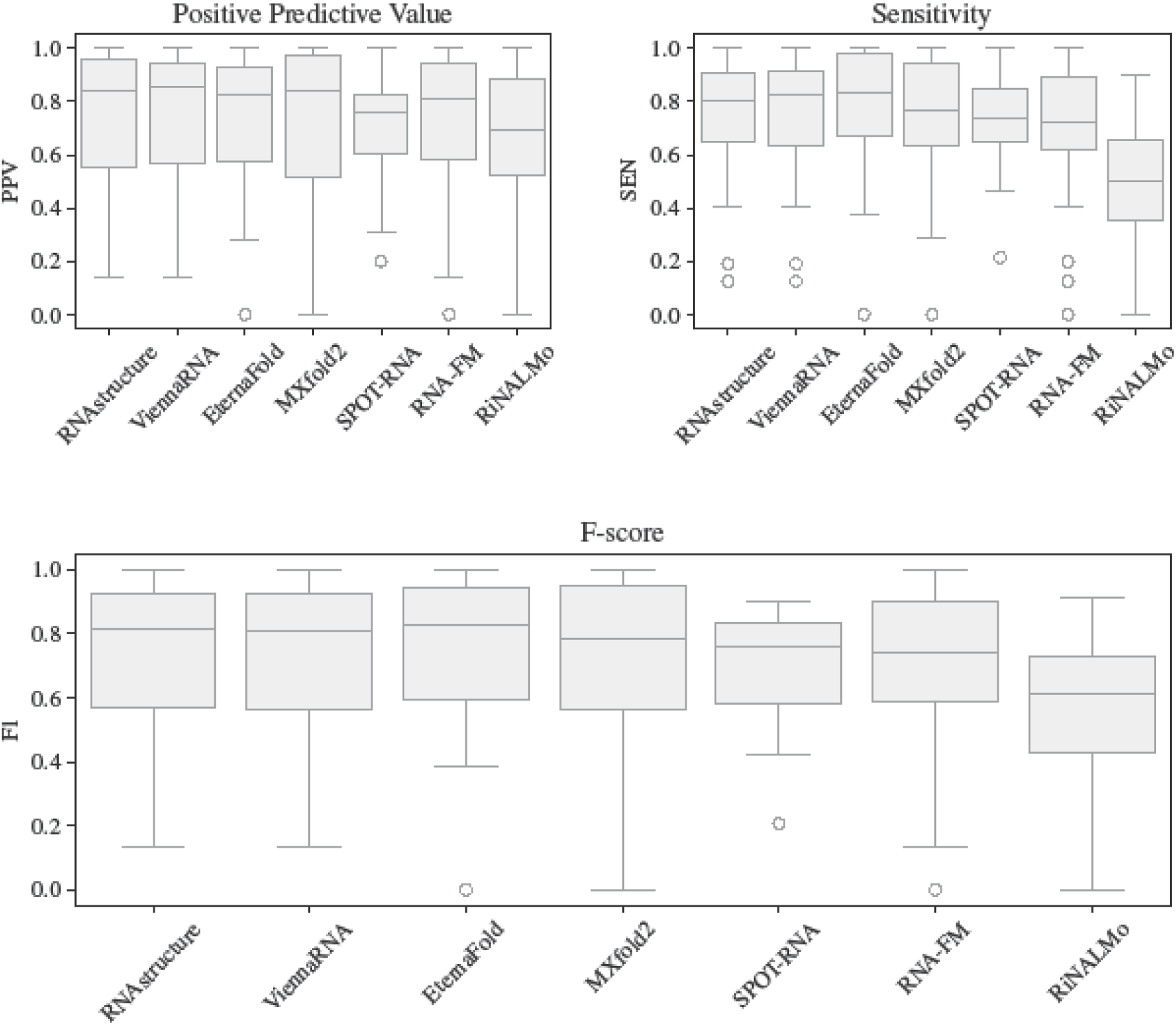
Boxplot of secondary structure prediction benchmarking results. Positive Predictive Value (PPV), Sensitivity (SEN), and F-score (𝐹*_1_*) of benchmarked data-driven methods compared to RNAstructure and ViennaRNA. Each group (e.g. Theophyline aptamer, minE/minF aptamer, Cas9 guide, etc.) is used as a single data point by averaging the targets within that group. All overlaps found in Supplemental Table S8 (*, **) and Supplemental Table S9 (†, ‡) are excluded from this calculation.

It should be noted that bpRNA-1m also contains a number of secondary structures derived from the PDB, some of which–as previously mentioned above–do not strongly hit any Rfam family. To address this, we have investigated the overlap of these PDB structures with our dataset. We find that Malachite green aptamer, r(CCUG), r(CUG), VS ribozyme, and ASH1 mRNA E3-localization element are also used in the bpRNA-1m dataset, meaning these targets are also a part of the training set of all evaluated models. Further, we have found two Theophyline aptamers, one Malachite green aptamer, an IAPV IRES, 13 VS ribozymes, and one r(CUG) repeat that is not directly in our dataset but belongs under one of our target groups. See Supplemental Table S8 for the overlap between bpRNA-1m’s PDB structures and our benchmarking dataset. Note that we exclude overlapping structures from our analysis here, see Supplemental Table S1-S3 for the full results.

It is visually clear from Fig. 1 that none of the deep learning methods tested demonstrate substantial improvement over thermodynamics. Pairwise paired t-tests between the methods showed no significant difference in mean performance between any models, except RiNALMo, which performed significantly worse than each other method. See Supplemental Table S7 for details.

The highest mean F-score method is EternaFold (𝐹*_1_* = *0*.*758*), a conditional log-linear model based on CONTRAfold, although the improvements over RNAstructure and ViennaRNA are not statistically significant (𝑝 = *0*.*592* and 𝑝 = *0*.*475*, respectively). It should also be noted that the Eterna dataset used to train the model contains many synthetic design structures, aptamers, and other possibly similar structures to our benchmarking set. We were unable to find strong direct similarity between our sequences (with the exception of a Theophylline aptamer [`8d28_A`]) and EternaFold’s training sequences. However, this is a poor replacement for true structural similarity, which unfortunately cannot be done as some of the data used to train EternaFold is chemical reactivity scores. As a consequence, we recommend interpreting EternaFold’s improvements over RNAstructure and ViennaRNA with caution.

Of the deep learning methods, excluding overlapping structures, MXfold2 is the best performing (𝐹*_1_* = *0*.*742*) and generally in agreement with the method’s self-reported inter-family performance, slightly outperforming RNAstructure and ViennaRNA although not significantly (𝑝 = *0*.*959* and 𝑝 = *0*.*800*, respectively). Previous findings by (Szikszai et al., 2022) also showed that MXfold2 was the method closest to inter-family generalization when evaluated using a particularly difficult family-fold cross-validation strategy on the relatively small ArchiveII dataset. MXfold2 includes “thermodynamic integration,” meaning the model is constrained to avoid predictions that do not agree with thermodynamics, a strategy demonstrably successful in avoiding poor generalization to RNAs unseen during training.

SPOT-RNA performs second best in the deep learning category when including all benchmarks (F1=0.743). However, for our Fig. 1, we exclude the above mentioned bpRNA-1m PDB targets, and a number of additional targets due to the pretraining strategy used by SPOT-RNA. This strategy uses a subset of the structures in the PDB, some of which overlap with our benchmarking dataset. We identified 29 structures (17 common PDB IDs) that are in both SPOT-RNA’s pretraining set and our benchmarking set with 23 groups that are present in both. See Supplemental Tables S9 for a detailed breakdown of overlapping structures, and Supplemental Fig. S1 for overall performance (including overlapping structures). After excluding all overlapping structures, the mean F-score (𝐹*_1_* = *0*.*702*) drops below RNA-FM (𝐹*_1_*= *0*.*710*), providing further evidence that data leakage is a significant concern during benchmarking, and showing reduced generalization for SPOT-RNA.

In the case of SPOT-RNA 2, we find it performs better than SPOT-RNA (see Supplemental Fig. S2, and Supplemental Tables S10-S16 for detailed results), although not significantly (𝑝 = *0*.*619*). Also note that as explained in Materials and Methods, a further six targets are excluded from testing SPOT-RNA 2. Finally, in the case of the language models, RNA-FM (𝐹*_1_* = *0*.*710*) performs better than RiNALMo (𝐹*_1_*= *0*.*544*), but still does not beat the performance of RNAstructure or ViennaRNA.

### Models for Structure Probing Data Prediction

A major bottleneck in deep-learning-based prediction of RNA structure is the limited volume and diversity of high-resolution structures and multiple sequence alignments (Schneider et al., 2023). This limitation has driven recent interest in utilizing structure probing data as a complementary, low-resolution source of structural information. Structure probing experiments currently provide the most affordable and practical strategy for extracting large-scale structural information from complex RNA samples (see reviews (Kwok et al., 2015; Spitale & Incarnato, 2023; Weeks, 2021). They use reagents (e.g., SHAPE and DMS), which differentially react with nucleotides depending on their structural context (e.g., paired/unpaired), and, most commonly, detect the reactions via next-generation sequencing. Thus, they measure each nucleotide’s *reactivity* to the reagent, which correlates with its propensity to be unpaired (Eddy, 2014; Sükösd et al., 2013; Weeks & Mauger, 2011).

Several hundred datasets are publicly available, with many encompassing transcriptome-scale information for a multitude of organisms, cell types, and experimental conditions (Mu et al., 2025). As such, they hold the potential to mitigate the data scarcity issue. Moreover, these data span a breadth of RNAs, notably mRNAs, which are markedly different in sequence and structure from the non-coding RNAs used to inform structure prediction models. This could enable a much broader exploration of the sequence-structure space. Nonetheless, their scale and breadth also pose a major hurdle to their integration into deep learning workflows, as they are dominated by long mRNA sequences, whose structures are unknown. Processing inputs of that size can be prohibitively computationally costly for most models currently used to predict structure (e.g., attention-based architectures). When structures are known, it is standard practice to fragment long sequences into smaller, independently-folding domains (Mathews et al., 1999), but in the absence of such knowledge, one needs to arbitrarily delineate a local, folding-relevant, sequence context. The resulting mismatch between the folding-relevant and the chosen contexts might introduce significant errors and impair our ability to learn the true underlying structures. Another challenge arises from a limitation inherent to probing assays, namely, that they glean incomplete and noisy information, which can be challenging to interpret or translate into explicit structures. As such, they are often used as auxiliary data in computational structure prediction to boost accuracy (see reviews (Aviran & Incarnato, 2022; Sloma & Mathews, 2015)).

Several models were recently developed to predict probing data. Bliss et al. used a convolutional neural network (CNN) to map local sequence and thermodynamics–derived structure features to SHAPE data obtained from ∼200 mRNAs (Bliss et al., 2020). More recently, two massively parallel studies were undertaken, where large-scale, custom-designed RNA libraries were probed and transformer-based models developed and fitted to the data (Boyd et al., 2023; He et al., 2024). Both studies attempted to enhance data diversity in terms of reagents used and sequences explored. Each group reported a sequence-to-signal model (Toneyan et al., 2022), where signal takes the form of reactivities or reads. He et al.’s data and model were acquired and developed in the context of a Kaggle challenge, a public competition dubbed Ribonanza. The sequences, model, and probing data are thus publicly available. However, this is not the case in (Boyd et al., 2023).

As noted above, the motivation underlying such endeavors is to enhance prediction of structure or structure-dependent properties, such as in-solution stability, and both studies demonstrated progress toward these goals. However, reactivity prediction is also useful for data imputation—a critical need in transcriptome-wide datasets. These data are plagued by missing values and noisy readouts, driven by low transcript abundance, solvent inaccessibility, and library preparation biases. Despite their limitations, they provide information on structural ensembles in a range of cellular conditions and thus were instrumental in advancing a swath of *in vivo* structure-function and comparative studies (see reviews (Bose et al., 2024; Choudhary et al., 2017; Wang et al., 2021; Weeks, 2021)). Enhancing their quality and scope may advance similar studies. Notwithstanding, the challenge here is that the underlying RNAs are long.

To address this need, (Gong et al., 2021) used a CNN and long short-term memory networks to predict reactivity from sequence and nearby reactivities and recently trained their model on a mixture of icSHAPE datasets and then fine-tuned to individual datasets (Mu et al., 2025). Imputation took place iteratively in rolling windows for which over 50% of the values were available. Lastly, a sequence-to-signal model was recently reported, which highlights another application of imputation (Mizrahi et al., 2025). Here, a model to predict protein binding transcriptome-wide *in vivo* was trained with RNA sequence and probing data as input. To augment the latter, an auxiliary CNN was trained on icSHAPE data and its predictions were used in training and in inference. A third utility of probing data models is to accelerate characterization of a reagent’s complex sequence-structure-signal relationship, particularly in single-stranded regions. To date, this has been done by probing small libraries of RNAs with known structures (Busan et al., 2019; Marinus et al., 2021; Xiao et al., 2022), hence probing such RNAs *in silico* may remove a data bottleneck.

Generalization challenges encountered in reactivity prediction extend beyond those common to structure predictors. In both domains, it is desirable to generalize to instances longer than and/or structurally dissimilar to seen data. However, probing data display variability even in the absence of sequence or structural differences (Kutchko & Laederach, 2017; Weeks, 2021), and this should be considered when evaluating performance and when curating training data. Contributing factors include technical variables, such as library preparation steps, sequencing coverage, and choices of reagent and modification detection strategy (reviewed in (Choudhary et al., 2017)). On top of these, experimental conditions, such as temperature, interactions with proteins, and ligand and salt concentrations can differ between studies and samples and result in detectable reactivity differences. With any such departure from the conditions in which training data were collected, one can expect performance degradation. One way to bridge this gap is to train on a mixture of datasets representing diverse conditions and possibly also further fine-tune to specific datasets (Boyd et al., 2023; Mu et al., 2025). A possible alternative could be to adjust the different datasets using techniques such as quantile normalization, to bring their statistical properties closer (Bolstad et al., 2003).

### Gauging Generalizability

We sought to gauge the generalizability of sequence-to-signal models as they are analogous to sequence-to-structure models. We focused on RibonanzaNet, hereafter called RNet, since the model and data are available, and a comprehensive assessment was reported (He et al., 2024). He et al. used reactivity mean absolute error (MAE) as an evaluation metric and trained RNet to minimize it over more than 8×10^5^ sequences of lengths 115-206 probed by SHAPE and DMS *in vitro*. They reported MAE≅0.14 for approx. 10^6^ test sequences of lengths 177-457. Sequences were compiled from diverse sources and derived from genomes and synthetic designs.

To evaluate performance, we curated a dataset of 2,678 sequences of lengths 27-1,024 probed *in vitro* by three SHAPE reagents or by DMS. These correspond to transcript regions from SARS-CoV-2, HIV-1, *E. coli*, and *H. sapiens*, for which probing data were available (Lajarte et al., 2024; Manfredonia et al., 2020; Mustoe et al., 2018; Siegfried et al., 2014). All studies but one probed full transcripts, whose lengths far exceed that of Ribonanza sequences. Since folding is highly context-dependent, arbitrary fragmentation might introduce sequence-data mismatches, thus we resorted to literature that delineated regions deemed to form structured, independently folding domains. For these three data sources, we labeled a region as validated if high fidelity methods (e.g., covariance modeling, NMR) established its folding and as putative if SHAPE-guided analyses revealed a high degree of structure. We applied Ribonanza’s normalization strategy to the data and computed MAEs between RNet’s predictions and our data. See Materials and Methods and Supplemental Methods for details.

To establish a baseline MAE, we computed the median reactivity per base (A, C, G, U) in each data source and predicted the base-specific and source-specific median for each nucleotide. Since the median is the value that minimizes the MAE, this represents a best guess for a null model that is based on statistical behavior. A comparison of MAE between RNet and our baseline revealed variable, data-specific trends (Fig. 2). For some data sources, RNet clearly improves upon the baseline whereas for others, it performs comparably and sometimes even worse. Even when performances differ, the MAEs are of the same order of magnitude and within 10%-20% of each other. At the single sequence level, RNet often outperforms the baseline, but not always.

**Figure 2.**
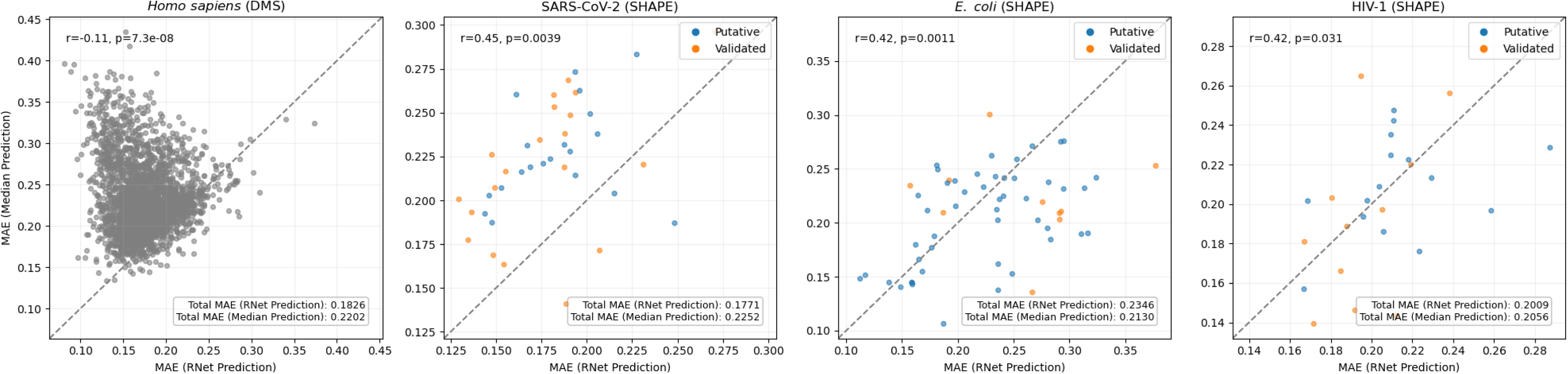
Comparison between RNet prediction error and a base-specific median predictor error. Each point represents one sequence, with blue and orange denoting putative and validated structural elements, respectively. The x-axis shows the MAE from RNet predictions while the y-axis shows the MAE from the nucleotide-wise median predictor, where medians were calculated separately for A, C, G, and U in each dataset. Points above and below the dashed diagonal line correspond to sequences for which RNet performs better and worse than the median predictor, respectively. Shown r and p are Pearson correlation coefficient and its associated p-value, respectively. In each panel, the inset reports the dataset-level MAE.

By considering *in vitro* SHAPE/DMS data, we minimize discrepancies rooted in experimental variables. In that context, it is worth noting that SHAPE is a family of reagents, with our SHAPE data obtained using NMIA, 1M7, and NAI and Ribonanza data obtained using 2A3. Despite shared chemical principles, these reagents can display differential reactivity patterns, thereby precluding full control of experimental variables (Marinus et al., 2021; Xiao et al., 2023). Additionally, we expect the aforementioned technical factors to give rise to inter-lab variability and higher MAEs than reported by He et al. (Supplemental Table S17). Here, we are most interested in gauging performance upon deviations from the training data with respect to sequence length or sequence-structure content. Since deep learning models tend to over-fit the length distribution of their training data, and since the training data for RNet is dominated by a specific length (177nt), we expected to see evidence of over-fitting on our test data. However, overall we did not observe MAE dependence on input length for instances within the range of RNet’s train or test data (Supplemental Fig. S3). When inputs exceed the longest test sequence, MAE increases very slowly, indicating the model is consistent across lengths and suggesting some evidence for generalization. Notably, we observed generally higher and more variable MAEs among short RNAs (< 100nt). We also did not observe MAE dependence on an element’s designation as putative or validated.

Importantly, He et al. padded short sequences in two ways, experimentally and computationally, to bring most sequences to the same length for efficiency purposes, with the former type also designed to reduce interactions with the original RNA. To test if inference is robust to experimental padding, we compared MAE for short RNAs with/without padding while retaining additional flanking sequences common to all Ribonanza data. The minor differences observed suggest invariance to this type of context extension (Supplemental Fig. S4). Since flanking sequences were added to all data, we also examined differences with/without them while keeping short RNAs padded. We found that predictions for short RNAs were sometimes more sensitive to such extensions (Supplemental Fig. S5). To ensure all inputs in a train/test batch have the same length, special padding tokens are typically added (computationally) to equalize lengths to the maximum input size. Computational padding is also applied in RNet. However, due to shuffling during training, one cannot determine which inputs were subject to padding and to which lengths (Materials and Methods). We therefore examined the impact of padding up to several lengths that dominate the Ribonanza data (Supplemental Fig. S6). We found that it introduced considerable performance variability, irrespective of RNA length, with highly variable gains or losses. It is worth noting that since certain sequence lengths dominate the data, we expect that padding was rare during training and testing, and it is likely best to avoid it at inference. However, we point this out to alert users re-training the model on variable-length data or using a pre-trained RNet to predict in batches.

Next, we set out to examine if high sequence similarity to one of the training sequences results in better predictions. We did not find a clear effect, however, we noticed substantial variation in MAE among 14 sequences with near-perfect or perfect similarity to training sequences, corresponding to both putative and validated elements (Supplemental Fig. S7-S8). A closer look revealed these are short sequences (27-88nt), hereafter called queries, each mapping to multiple training instances, hereafter called targets. After restricting analysis to targets with high-quality data, we inspected the multiple targets that matched each of the remaining ten queries and observed significant discrepancies between their reactivity profiles (see, e.g., Supplemental Fig. S9), likely arising due to a small number of mutations inside or outside each query. This highlights the major role sequence context plays in folding and the need to quantify similarity at the signal or structure level. It also indicates that these target profiles display considerably different similarities (in reactivity) to our data (Supplemental Fig. S10). Yet, we found that when target profiles are, on average, more similar to our data, the prediction error tends to be smaller. For each query, we then averaged its target profiles nucleotide-wise and compared to RNet’s prediction. Interestingly, the average profiles are generally concordant with RNet’s predictions and appear to be much more similar to them than individual training profiles (Fig. 3 and Supplemental Fig. S11-S12). Furthermore, using the average target profile as a predictor tends to slightly outperform RNet (Supplemental Fig. S13). See Materials and Methods for details.

**Figure 3.**
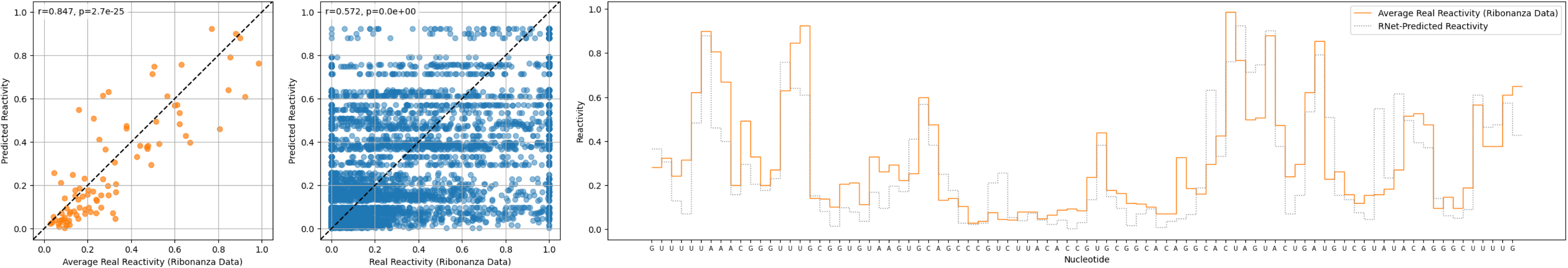
Comparison of RNet predicted reactivity profile against averaged and individual reactivity profiles for Ribonanza training sequences highly similar to the query region (SARS-CoV-2, 13459-13546). Left: Predicted reactivity plotted against the averaged target reactivity. Middle: Predicted reactivity plotted against multiple individual target reactivities, illustrating high variability among targets. Right: Reactivities predicted by RNet (gray) and reactivities derived from Ribonanza data by averaging over all targets (orange). Also shown are the Pearson correlation coefficient (r) and its associated p-value (p) and the query sequence.

### Measuring Dissimilarity in Signal Space

Our use of MAE as a performance measure enabled a direct comparison to He et al.’s results. While RNet was trained to minimize MAE loss, other works minimized mean squared error (MSE) loss (Bliss et al., 2020; Gong et al., 2021; Mizrahi et al., 2025) and evaluated performance via Pearson correlation between predictions and real data and/or select structure predictions constrained by predicted data (Bliss et al., 2020; Mu et al., 2025).

Despite the popularity of MAE/MSE in learning and benchmarking regression models, it is worth considering alternative loss (or dissimilarity) formulations tailored to the unique properties of probing data. One such property is asymmetry. First, it has been observed that higher reactivities tend to display higher variances or lower concordance (i.e., heteroskedasticity), and so higher errors are to be expected in this regime (Choudhary et al., 2016; Low & Weeks, 2010). Second, information content is not uniformly distributed over the dynamic range (Deng et al., 2016). For example, using SHAPE data for RNAs with known structures, it was shown that while zero/near-zero reactivities provide the strongest evidence a base is paired, we quickly lose certainty as reactivities increase, to the point where values in [0.2, 0.3] are no longer informative (i.e., paired/unpaired is equally likely). The situation is different among high reactivities (e.g., >=0.7), which all provide similarly strong evidence of the unpaired state (Eddy, 2014). What this means is that mispredicting 0 as 0.25 is likely to have much greater impact on data interpretation and predicted data-constrained structures than mispredicting 0.7 as 0.95. Moreover, when interpreting MAE, one should keep in mind that the data are highly skewed towards zero, and thus mispredictions of low reactivities dominate it (Sükösd et al., 2013). Taken together, these suggest that metrics or data transformations more tolerant of errors among high values may better capture the similarity between the underlying secondary structures.

Another consideration in assessing similarity is that significant structure differences tend to manifest as reactivity differences spanning a few to tens of adjacent nucleotides (Woods & Laederach, 2017). Such differences are often visually detectable as mismatches in motif-level data patterns, such as the “valleys” and “peaks” typically seen in double-and single-stranded regions, respectively. Detecting differences at the region and/or pattern level can also reduce the noise introduced by each reactivity (Ledda & Aviran, 2018; Selega et al., 2017). Unfortunately, MAE/MSE is oblivious to patterns and does not pool information across adjacent nucleotides. Nonetheless, data-driven comparative analyses have grappled with the question of how to detect changes at the level of structure directly from probing data (see review (Choudhary et al., 2017)). Numerous differential analysis methods were developed to quantitatively address this need while accounting for common confounding factors (Choudhary et al., 2019; Marangio et al., 2021; Mizrahi et al., 2018; Smola et al., 2015; Tapsin et al., 2018; Wan et al., 2014; Woods & Laederach, 2017; Yu et al., 2022). All methods quantify differences at a regional level, and some are also pattern-aware, yet they each approach similarity quantification differently. This body of work can provide insights and sophisticated tools for improved assessment of reactivity predictors. In cases where similarity is quantified through a differentiable function, one could also envision leveraging it as a training objective.

Finally, there are additional, indirect ways to assess reactivity prediction models. First, one could assess statistical, population-level properties by making predictions for RNAs with known structures and comparing structural context-dependent distributions between predicted and real data (Sükösd et al., 2013). Second, one could compare the structures predicted by algorithms guided by the predicted data against predictions guided by the real data.

### Deep Learning Broadens the Scope of Traditional Methods

Beyond improving predictive accuracy, recent advances have shown that traditional methods can be optimized through a differentiable framework. This approach blurs the boundary between physical modeling and machine learning, enabling both thermodynamic and neural models to be refined using gradient-based principles.

Deep learning models are trained through gradient descent, whose success has shown the power of gradient-based optimization in complex, high-dimensional spaces (Ruder, 2017). This is enabled by automatic differentiation using hardware accelerators, which efficiently backpropagates gradients through a model’s computational graph (Baydin et al., 2018). Gradient computation has long been central to numerous approaches to RNA secondary structure prediction. In these frameworks, training involves maximizing the log-likelihood of structural data, such as observed sequence-structure pairs and/or structure probing measurements (Clote, 2025). The inside-outside algorithm, a dynamic programming procedure analogous to backpropagation (Eisner, 2016), recursively computes a vector of expected feature/rule counts, which is a core component of the log-likelihood’s gradient. The inside-outside recursions were derived manually, thereby enabling complex end-to-end gradient-based learning decades before modern automatic differentiation frameworks. They have been used to learn stochastic context-free grammar (SCFG) and conditional log-linear models (Andronescu et al., 2010; Do et al., 2006; Rivas et al., 2012; Wayment-Steele et al., 2022).

A complementary paradigm for secondary structure prediction is based on Turner’s nearest-neighbor model of folding thermodynamics (Mathews et al., 2004b; Mittal et al., 2024). This physically grounded framework assigns free energy changes to local structural motifs such as stacks, loops, and junctions, using a large set of experimentally derived parameters. Historically, these parameters were obtained from optical-melting experiments through expert-driven fitting procedures that rely on carefully designed constructs and specialized statistical techniques (Andronescu et al., 2014; Zuber et al., 2018). Andronescu *et al*. later refined the Turner model using maximum-likelihood optimization that combined sequence-structure pairs and thermodynamic measurements (Andronescu et al., 2007). They manually derived the required gradients through the inside-outside recursions, but evaluating them proved computationally expensive and not practically scalable (Andronescu et al., 2010).

These limitations and the advent of deep learning motivated a new generation of *differentiable folding* methods that compute gradients automatically through the dynamic programming recursions underlying RNA folding algorithms (e.g., McCaskill’s algorithm for computing the RNA partition function) (Krueger & Ward, 2025; Matthies et al., 2024). Rather than relying on manually derived analytical gradients for specific objectives, these methods leverage novel software frameworks that enable automatic differentiation, such as JAX (Bradbury et al., 2018), to backpropagate through the entire folding computation. This enables efficient, scalable, and exact gradient computation for arbitrary differentiable objectives, with respect to thousands of parameters simultaneously, transforming previously intractable model-fitting problems into routine optimization tasks. For example, Krueger *et al*. define a structural objective based on the geometric mean of log-probabilities across RNA families, ensuring that all families contribute equally to the optimization, and combine it with a thermodynamic loss term that enforces agreement with optical-melting experiments (Krueger et al., 2025). The resulting composite objective is optimized by gradient descent, yielding optimized nearest-neighbor parameters that improve both structural accuracy and thermodynamic consistency across the ArchiveII dataset and a new RNAometer benchmark dataset that contains a database of experimentally-determined free energy changes. The improvements are substantial: optimized models increase the predicted probabilities of native sequence-structure pairs by many orders of magnitude compared with the Turner 2004 parameters. This work demonstrates how the new platform enables the use of flexible objective functions that explicitly balance biases across heterogeneous data sources.

While differentiable folding was developed in the context of thermodynamic models, its flexibility in defining custom objectives also allows probabilistic frameworks to be revisited. For example, Pratap *et al*. applied this paradigm to train SCFG parameters directly on natural RNA sequences, without any structural supervision, by maximizing only the likelihood of the observed sequences (Pratap et al., 2025). Remarkably, canonical Watson-Crick-Franklin and wobble base-pairing rules emerged spontaneously after just a few epochs of training, and the resulting 21-parameter model achieved comparable accuracy to thermodynamic predictors trained on structural data. This work underscores how differentiable frameworks expand the model design space, enabling unconventional training objectives such as unsupervised discovery of structural rules directly from sequence data.

Beyond improving existing thermodynamic and probabilistic frameworks, differentiable folding establishes a new bridge between physics-based modeling and deep learning. Optimized parameters can generate higher-fidelity base-pairing probabilities or other ensemble observables as training data, while the differentiable algorithms themselves can be embedded as modules within larger neural architectures. For example, in a neural network trained to predict RNA tertiary structure, one could supply the outputs of a dynamic-programming-based base-pair probability (BPP) prediction module whose folding parameters are jointly trained with the weights of the neural network, rather than relying on fixed BPP inputs. This is analogous to how differentiable sequence alignment algorithms have enabled end-to-end training in protein modeling (Petti et al., 2023). This convergence suggests a unified direction for RNA structure prediction: rather than viewing thermodynamic and deep-learning models as competing paradigms, differentiable optimization turns them into interoperable components of a single, trainable ecosystem that links physical interpretability with data-driven generalization.

## CONCLUSION

The field of RNA structure prediction is undergoing tremendous growth driven by rapid developments in artificial intelligence. Considering that the diversity and volume of known secondary structures is limited and challenging to grow with existing technologies, developing improved technologies to collect more, better, and different data along with methods that better utilize existing data to avoid over-fitting may be more critical to closing the generalization gap than scaling the size and assortment of deep learning models. Alongside such advances, careful performance evaluation must also take place.

Our benchmarks on deep learning methods that predict secondary structure directly have shown that thermodynamic folding is still as relevant as ever for structure prediction. While some deep learning methods, like MXfold2 (which is constrained by thermodynamics) do not suffer from issues with generalization, they still do not outperform RNAstructure or ViennaRNA on a test set not represented in the training data. We further showed that EternaFold performs well on our targets, which are largely made up of aptamers and synthetic structures. Although once again, EternaFold still does not show statistically significant improvements over RNAstructure or ViennaRNA, despite the possibility of data leakage between our test set and the training set. EternaFold uses the recursions of the nearest neighbor model and simply fits new parameters, meaning it is a mixed machine learning and thermodynamics model rather than de novo deep learning model (such as SPOT-RNA and RiNALMo). This further supports our observation that data-driven methods perform better when constrained by thermodynamics, such as MXfold2 and EternaFold.

One potential direction for computational advancement entails improving how deep learning models use known structural constraints during training. While most existing neural architectures output pairwise weight matrices that are later converted into discrete base-pairing structures, the training and extraction stages are often treated as independent. This disconnect can limit performance as a model may minimize numerical loss between predicted and target contact maps yet fail to produce physically valid or consistent secondary structures. A similar challenge arose in the development of AlphaFold where predicted 3D coordinates must satisfy the triangle inequality, prompting the development of the triangle attention mechanism to enforce geometric consistency. Analogous architectural innovations that explicitly couple training objectives with the structural constraints embedded in the downstream extraction process could likewise prove critical for improving base-pair prediction.

Leveraging additional, complementary sources of information, such as structure probing, is another promising direction, where here too, care must be taken in choosing training objectives and in assessing performance. Recent major initiatives entailing the design and production of massive RNA libraries followed by structure probing and the fitting of neural networks to these data underscore the potential of these assays to circumvent current limitations and enhance structure learning. Our analysis highlighted the strengths and weaknesses of pioneering efforts to model these data and we hope these will stimulate additional algorithmic innovation. Such undertakings also provide new opportunities to judiciously design both train and test data to maximize structural diversity within and between train and test subsets. Such a degree of control over the data is unusual and not feasible when learning from a limited repertoire of known sequence-structure pairs. Yet, achieving structural diversity via sequence design is not a trivial task and warrants additional method development.

## MATERIALS AND METHODS

### Software and Datasets

Ground-truth secondary structures for our benchmarking dataset, as well as predictions are available in Supplemental File 2. Detailed descriptions and information relating to the ground-truth structures is available in Supplemental File 1. Datasets used in RNet’s performance evaluation are summarized in Supplemental Methods. Sequences and probing data can be found in Supplemental File 3.

### Structure Data Curation

We present a benchmarking dataset of RNA structures, Archive-NoFam, that shares no homology with any RNAs in existing and commonly used datasets for training secondary structure prediction models. These sequence-structure pairs are curated from the PDB with the purpose of finding molecules that are not present in existing training sets to simulate the structure prediction of novel RNAs.

To construct our dataset, we start with the RNA3DB (version 2024-12-04-full-release^2^) dataset (Szikszai et al., 2024). This dataset divides all chains in the PDB into structurally dissimilar clusters, termed “components”. We use the component that shares no homology to any Rfam family (termed “component 0” by RNA3DB’s nomenclature), containing 1,167 3D structures in total. These chains are filtered for our benchmarking dataset, first by systematically removing structurally uninteresting RNAs (e.g. no secondary structure), and then further filtered manually to identify suitable target structures.

Suitable target structures are manually identified by considering a wide range of criteria, including resolution, structure determination method, clash score, 2D structure, and origin. We required that the RNA chain component of the structure be resolved continuously, allowing missing portions at the 5’ or 3’ ends. We allowed exceptions to this rule for missing nucleotides in hairpin loops. In the end, this results in 89 total chains, which can be divided into a set of 47 structurally dissimilar groups. These groups are made up of 18 aptamers, 3 CRISPR guide RNAs, 4 internal ribosome entry sites (IRES), 5 ribozymes, 3 disease-associated nucleotide repeats, 5 miscellaneous synthetic

To obtain sequences suitable for all benchmarked methods, we use RNA3DB’s parser to remove modifications from our dataset. This parser “demodifies” all residues and converts them into an alphabet of {𝐴, 𝐶, 𝐺, 𝑈} sequences by using the closest canonical nucleotide equivalent to each modification. Base pairs are extracted via RNAView (Yang et al., 2003), which classifies them according to Leontis-Westhof notation (Leontis & Westhof, 2001). These base pairs are processed into secondary structures by considering only 𝐺 − 𝐶, 𝐴 − 𝑈, and 𝐺 − 𝑈 (wobble) Watson-Crick-Franklin/Watson-Crick-Franklin pairs with cis glycosidic bond orientation (𝑊𝑊𝑐). Incompatible overlapping base pairs (e.g. two 𝑊𝑊𝑐 pairs to the same residue) are removed by computing the maximum-cardinality secondary structure. As the final pre-processing step, we also remove residues from the 5’ and 3’ ends of the sequences with unresolved 3D structures (e.g. `2xdb_G`, `8w35_C`) or if they are in a duplex (e.g. `8gkh_W`, `7el1_B`).

### Structure Prediction Benchmarking Strategy

Recent reviews of deep learning methods for secondary structure prediction by (Zablocki et al., 2025) and (Chaturvedi et al., 2025) have discussed the generalization gap for many existing methods. As a result, our methodology focuses on the most popular methods which have not shown poor generalization to date. To identify methods we choose to benchmark, we have taken the top 20 papers for the search term “deep learning RNA secondary structure” after 2020, based on the *Citation Count* sorting criteria from *Semantic Scholar* and *relevance* sorting criteria from *Google Scholar* (retrieved 2025-08-27). See Supplemental Tables S18-S19 for details.

After filtering for only relevant papers, this yields six methods: MXfold2 (Sato et al., 2021), UFold (Fu et al., 2021), E2Efold (X. Chen et al., 2020), SPOT-RNA 2 (Singh et al., 2021), RiNALMo (Penić et al., 2025), and REDfold (C.-C. Chen & Chan, 2023). We also identified RNA-FM (J. Chen et al., 2022) as another language model to compare against RiNALMo, and also opted to include SPOT-RNA (Singh et al., 2019), which is an earlier version of SPOT-RNA 2 that does not use alignments. For completeness, we also included EternaFold (Wayment-Steele et al., 2022) in our analysis, a conditional log-linear model trained on “over 20,000 synthetic RNA constructs designed on the RNA design platform Eterna” in addition to the traditional secondary structure dataset used by other methods.

For the final comparison, we benchmarked RNAstructure (Reuter & Mathews, 2010) and ViennaRNA (Lorenz et al., 2011) (both traditional thermodynamics models that use the Turner 2004 thermodynamic model (Mathews et al., 2004b) although with differences in the energy calculation for multibranch and exterior loops) against MXfold2, SPOT-RNA, RiNALMo, and EternaFold on our proposed dataset. In the case of RNA-FM and RiNALMo, which both make multiple sets of structure prediction head weights available, we used the heads trained on bpRNA-1m. UFold and E2Efold were omitted from our testing as previous literature already established poor generalization (Szikszai et al., 2022), and REDfold was also excluded as they were not able to show evidence of inter-family generalization in their benchmarks. All calculations of Positive Predictive Value, Sensitivity, and F-score were done using exact comparison, without allowing for flexible pairings as sometimes done when using structures derived from comparative sequence analysis per (Mathews, 2019).

Despite using alignments, we also benchmarked SPOT-RNA 2 on our dataset, although we were unable to run the program on six of our targets (VS ribozyme [4r4v_A], IAPV IRES [6p5i_1, 6p5n_1], CrPV 5’UTR IRES [6w2t_A], TSV IRES [8evp_EC], Cas12 guide RNA [8rdu_1]) due to a segmentation fault in the inference pipeline.

### Reactivity Prediction Benchmarking Strategy

We re-normalized the probing data extracted from each study to account for the fact that these studies normalized their data differently. We followed He et al.’s strategy and scaled the data from each source (separately), such that the 90th percentile was set to 1. For DMS data, we only considered data for A and C. Prior to computing MAE, values were clipped to [0,1] to match He et al.’s workflow. We computed MAE per sequence and per data source as 𝑀𝐴𝐸 = 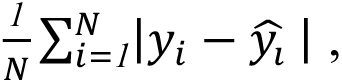 where *N* is the number of values, 𝑦*_i_* is the *ith* value, and 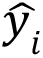 is the *ith* predicted value.

For prediction, we used He et al.’s source code^3^ with the weights available on the Kaggle site^4^. We also used their code to mimic experimental sequence padding and to pad sequences computationally. See Supplemental Methods for details. Note that computational padding is applied during training as well as when making predictions for multiple sequences. However, during training, sequences are randomly shuffled before being grouped into batches, where padding length is determined by the longest sequence in a batch. Because shuffling takes place each epoch and batch composition is not recorded, it is impossible to trace the batches to which a sequence was assigned and their padding length.

We used MMseqs2 (version 14-7e284, easy-search with parameters --min-seq-id 0.0, --search-type 3,-a) to assess sequence similarity (Steinegger & Söding, 2017). Our sequences were used as queries and Ribonanza sequences as targets. From the alignment output, similarity was calculated as the number of identical bases (nident) divided by the query length (qlen). For each query, the maximum similarity across all alignments was recorded. For analyses focusing on high-similarity sequences, we further removed Ribonanza training sequences of low quality, specifically those with SN_filter=0 (corresponding to signal-to-noise < 1.0.00 or reads < 100).

## ACKNOWLEDGEMENTS

This work was partially supported by NIH grants R21GM148835 to S.A. and R35GM145283 to D.H.M. Some of the material is based upon work supported by NSF grant UWSC13223.

## Footnotes

1 ^1^Note that this is an extremely generous threshold to minimize false negatives.

2 ^2^Retrieved from: https://github.com/marcellszi/rna3db/releases/tag/2024-12-04-full-releasestructures, and 9 miscellaneous non-synthetics. For more details about the dataset, see Supplemental Fig. S14-S15.

3 ^3^Downloaded from: https://github.com/Shujun-He/RibonanzaNeton4/10/25

4 ^4^Downloaded from: https://www.kaggle.com/datasets/shujun717/ribonanzanet-weightson4/29/25

