## Supplemental Tables for "Deep Learning for RNA Secondary Structure Determination: Gauging Generalizability and Broadening the Scope of Traditional Methods"

**2 bpRNA-1m PDB set homology** **10**

**3 SPOT-RNA transfer learning PDB set homology** **12**

**4 SPOT-RNA & SPOT-RNA 2 results** **14**

### 1 Single-sequence benchmarking results

#### 1.1 Positive Predictive Value

| | RNAstructure | ViennaRNA | EternaFold | MXfold2 | SPOT-RNA | RNA-FM | RiNALMo |
|---|---|---|---|---|---|---|---|
| **Aptamers** | | | | | | | |
| Theophyline aptamer* | 0.80 | 0.80 | 0.76 | 0.80 | 0.64 | 0.80 | **0.88** |
| minE/minF aptamer | 0.75 | 0.75 | **0.82** | 0.77 | 0.68‡ | 0.74 | 0.72 |
| RhoBAST aptamer | **1.00** | **1.00** | 0.94 | **1.00** | 0.81 | 0.97 | 0.88 |
| Spinach aptamer | 0.61 | 0.59 | 0.58 | 0.37 | **0.75**† | 0.59 | 0.53 |
| Mango aptamer | 0.73 | 0.73 | **0.92** | 0.91 | 0.73‡ | 0.87 | 0.89 |
| Vitamin B12 aptamer | 0.14 | 0.14 | 0.60 | 0.75 | 0.64† | 0.14 | **1.00** |
| Squash aptamer | 0.84 | **0.91** | 0.87 | **0.91** | 0.89 | 0.70 | 0.81 |
| Pepper aptamer | **1.00** | **1.00** | 0.00 | **1.00** | 0.83 | **1.00** | 0.00 |
| Chili aptamer | 0.53 | 0.53 | 0.65 | 0.50 | **0.80** | **0.80** | 0.67 |
| Corn aptamer | 0.45 | 0.45 | 0.42 | 0.50 | **0.64**‡ | 0.45 | 0.50 |
| DIR2s aptamer | 0.88 | **1.00** | 0.75 | 0.78 | 0.94‡ | **1.00** | 0.93 |
| Clivia aptamer | 0.55 | 0.57 | 0.71 | **1.00** | 0.71 | 0.80 | 0.67 |
| A9g aptamer | **0.82** | 0.45 | 0.50 | 0.45 | 0.42 | 0.55 | 0.67 |
| Beetroot aptamer | **0.92** | **0.92** | **0.92** | **0.92** | 0.61 | 0.83 | 0.59 |
| Malachite green aptamer** | **1.00** | **1.00** | **1.00** | **1.00** | 0.92‡ | **1.00** | **1.00** |
| 11F7t aptamer | 0.90 | 0.90 | **1.00** | 0.90 | 0.85‡ | **1.00** | 0.62 |
| K1 aptamer | **1.00** | **1.00** | **1.00** | **1.00** | 0.70‡ | **1.00** | 0.75 |
| Tetracycline aptamer | **1.00** | 0.94 | 0.94 | **1.00** | **1.00**‡ | **1.00** | **1.00** |
| **CRISPR RNA guides** | | | | | | | |
| Cas9 guide | 0.87 | 0.87 | 0.92 | **0.93** | 0.78 | 0.92 | 0.65 |
| Cas12 guide | 0.56 | 0.58 | 0.56 | 0.60 | 0.61 | **0.74** | 0.58 |
| Cas13 guide | 0.48 | 0.49 | 0.48 | **0.51** | 0.39‡ | 0.50 | 0.25 |
| **IRES** | | | | | | | |
| IAPV IRES* | 0.46 | 0.45 | 0.43 | 0.48 | **0.52**† | 0.51 | 0.35 |
| CrPV 5'UTR IRES | 0.22 | 0.21 | 0.28 | 0.22 | 0.22† | **0.34** | 0.15 |
| TSV IRES | 0.29 | 0.29 | 0.48 | 0.51 | 0.56 | 0.51 | **0.58** |
| PSIV IGR IRES | 0.64 | 0.64 | 0.82 | 0.82 | 0.82‡ | 0.82 | **1.00** |
| **Ribozymes** | | | | | | | |
| Synthetic ligase ribozyme | 0.84 | 0.85 | 0.85 | 0.85 | 0.76‡ | **0.88** | 0.75 |
| VS ribozyme** | 0.62 | 0.90 | 0.83 | 0.92 | **0.93**† | 0.63 | 0.42 |
| Self-alkylating ribozyme | **1.00** | **1.00** | 0.83 | **1.00** | 0.80 | 0.86 | 0.18 |
| Diels-Alder ribozyme | **1.00** | 0.89 | 0.89 | **1.00** | 0.73 | 0.89 | 0.88 |
| Methyltransferase ribozyme | **0.97** | 0.87 | 0.95 | **0.97** | 0.93 | 0.90 | 0.88 |

Continued on next page

| | RNAstructure | ViennaRNA | EternaFold | MXfold2 | SPOT-RNA | RNA-FM | RiNALMo |
|---|---|---|---|---|---|---|---|
| **Disease-associated nucleotide repeats** | | | | | | | |
| r(CUG)** | **1.00** | **1.00** | **1.00** | **1.00** | 0.75 | **1.00** | **1.00** |
| r(CCUG)** | **1.00** | **1.00** | **1.00** | **1.00** | 0.80‡ | **1.00** | 0.85 |
| r(AUUCU) | **1.00** | **1.00** | **1.00** | **1.00** | 0.77 | **1.00** | **1.00** |
| **Miscellaneous synthetics** | | | | | | | |
| Nanoarchitecture 1 | 0.88 | 0.88 | **0.93** | 0.33 | 0.20 | 0.00 | 0.00 |
| Nanoarchitecture 2 | **0.95** | **0.95** | **0.95** | **0.95** | 0.74 | 0.67 | 0.45 |
| Synthetic hairpin | **1.00** | **1.00** | **1.00** | **1.00** | 0.75 | **1.00** | **1.00** |
| Synthetic tetraloop-tetraloop receptor | 0.86 | 0.86 | 0.91 | **0.94** | 0.88‡ | 0.94 | 0.93 |
| Synthetic G-quadruplex | 0.62 | 0.62 | 0.62 | **0.83** | 0.64† | 0.62 | 0.67 |
| **Miscellaneous** | | | | | | | |
| ToxI | 0.27 | 0.27 | 0.53 | 0.00 | **0.58**‡ | 0.42 | 0.26 |
| Structured part of acrIF8-aca2 5' UTR | 0.92 | 0.92 | 0.92 | 0.75 | **1.00** | **1.00** | **1.00** |
| Saguaro cactus viral mRNA structure | 0.83 | 0.84 | **0.97** | 0.90 | 0.86 | 0.77 | 0.82 |
| ITS-2 | 0.56 | 0.58 | 0.56 | 0.65 | 0.46 | 0.62 | **0.82** |
| ENE | 0.95 | 0.95 | 0.91 | 0.98 | 0.90† | 0.98 | **0.99** |
| ASH1 mRNA E3-localization element** | **0.80** | **0.80** | **0.80** | **0.80** | 0.62‡ | **0.80** | 0.69 |
| SCNMV xrRNA | **1.00** | **1.00** | 0.73 | 0.78 | 0.75‡ | 0.90 | 0.83 |
| Influenza B vRNA promoter | **0.36** | **0.36** | 0.29 | **0.36** | 0.31 | 0.31 | 0.33 |
| NAD-II riboswitch | 0.44 | 0.44 | 0.68 | 0.50 | **0.87** | 0.25 | 0.39 |
| **Descriptive statistics** | | | | | | | |
| Mean | 0.75 | 0.75 | 0.76 | **0.77** | 0.71 | 0.74 | 0.68 |
| Median | 0.84 | **0.86** | 0.83 | 0.85 | 0.75 | 0.80 | 0.72 |
| Min | 0.14 | 0.14 | 0.00 | 0.00 | **0.20** | 0.00 | 0.00 |
| Max | **1.00** | **1.00** | **1.00** | **1.00** | **1.00** | **1.00** | **1.00** |
| Std. Dev. | 0.25 | 0.25 | 0.23 | 0.25 | 0.19 | 0.26 | **0.28** |

**Supplemental Table S1**. **Positive Predictive Value (PPV) on the proposed benchmarking dataset.** Values with † indicate homology to SPOT-RNA's PDB transfer learning dataset, while values with ‡ indicate exact match between PDB structures.

#### 1.2 Sensitivity

| | RNAstructure | ViennaRNA | EternaFold | MXfold2 | SPOT-RNA | RNA-FM | RiNALMo |
|---|---|---|---|---|---|---|---|
| **Aptamers** | | | | | | | |
| Theophyline aptamer* | 0.97 | 0.97 | **1.00** | 0.70 | **1.00** | 0.97 | 0.60 |
| minE/minF aptamer | 0.65 | 0.65 | 0.84 | 0.73 | **0.92**$^{\ddagger}$ | 0.73 | 0.63 |
| RhoBAST aptamer | 0.78 | **0.83** | **0.83** | 0.81 | 0.72 | 0.74 | 0.62 |
| Spinach aptamer | 0.62 | 0.64 | 0.62 | 0.36 | **0.86**$^{\dagger}$ | 0.44 | 0.36 |
| Mango aptamer | 0.83 | 0.83 | **0.97** | **0.97** | **0.97**$^{\ddagger}$ | **0.97** | 0.66 |
| Vitamin B12 aptamer | 0.12 | 0.12 | 0.38 | 0.38 | **0.88**$^{\dagger}$ | 0.12 | 0.12 |
| Squash aptamer | **0.88** | **0.88** | 0.83 | **0.88** | 0.67 | 0.67 | 0.54 |
| Pepper aptamer | **0.94** | **0.94** | 0.00 | **0.94** | 0.62 | **0.94** | 0.00 |
| Chili aptamer | 0.60 | 0.60 | 0.73 | 0.60 | **0.80** | **0.80** | 0.40 |
| Corn aptamer | 0.71 | 0.71 | 0.71 | 0.71 | **1.00**$^{\ddagger}$ | 0.71 | 0.57 |
| DIR2s aptamer | 0.79 | **0.89** | 0.79 | 0.74 | 0.84$^{\ddagger}$ | 0.68 | 0.68 |
| Clivia aptamer | 0.74 | 0.82 | **1.00** | **1.00** | 0.82 | **1.00** | 0.65 |
| A9g aptamer | **1.00** | 0.56 | 0.67 | 0.56 | 0.56 | 0.67 | 0.67 |
| Beetroot aptamer | **0.85** | **0.85** | **0.85** | **0.85** | 0.46 | 0.62 | 0.38 |
| Malachite green aptamer** | **1.00** | **1.00** | **1.00** | **1.00** | 0.85$^{\ddagger}$ | 0.77 | 0.77 |
| 11F7t aptamer | 0.82 | 0.82 | **1.00** | 0.82 | **1.00**$^{\ddagger}$ | 0.91 | 0.45 |
| K1 aptamer | **1.00** | **1.00** | **1.00** | **1.00** | **1.00**$^{\ddagger}$ | **1.00** | 0.86 |
| Tetracycline aptamer | 0.89 | 0.89 | **0.94** | **0.94** | 0.89$^{\ddagger}$ | 0.72 | 0.61 |
| **CRISPR RNA guides** | | | | | | | |
| Cas9 guide | 0.90 | 0.91 | **0.96** | **0.96** | 0.88 | 0.87 | 0.34 |
| Cas12 guide | 0.58 | 0.60 | **0.62** | 0.60 | 0.57 | 0.60 | 0.41 |
| Cas13 guide | **0.60** | **0.60** | **0.60** | **0.60** | 0.54$^{\ddagger}$ | 0.52 | 0.25 |
| **IRES** | | | | | | | |
| IAPV IRES* | 0.74 | 0.74 | 0.74 | **0.75** | 0.68$^{\dagger}$ | 0.64 | 0.30 |
| CrPV 5'UTR IRES | 0.52 | 0.52 | **0.61** | 0.52 | 0.25$^{\dagger}$ | 0.50 | 0.20 |
| TSV IRES | 0.44 | 0.44 | **0.69** | 0.64 | 0.64 | 0.62 | 0.36 |
| PSIV IGR IRES | 0.50 | 0.50 | 0.64 | 0.64 | **1.00**$^{\ddagger}$ | 0.64 | 0.50 |
| **Ribozymes** | | | | | | | |
| Synthetic ligase ribozyme | 0.71 | 0.75 | **0.77** | 0.75 | 0.69$^{\ddagger}$ | 0.76 | 0.45 |
| VS ribozyme** | 0.59 | 0.93 | 0.89 | **0.95** | 0.82$^{\dagger}$ | 0.43 | 0.21 |
| Self-alkylating ribozyme | **0.94** | **0.94** | 0.83 | **0.94** | 0.67 | 0.67 | 0.11 |
| Diels-Alder ribozyme | **1.00** | **1.00** | **1.00** | **1.00** | **1.00** | **1.00** | 0.88 |
| Methyltransferase ribozyme | 0.88 | 0.82 | **0.97** | 0.85 | 0.88 | 0.82 | 0.49 |
| **Disease-associated nucleotide repeats** | | | | | | | |
| r(CUG)** | 0.83 | 0.83 | 0.92 | 0.92 | **1.00** | 0.92 | 0.67 |

Continued on next page

| | **RNAstructure** | **ViennaRNA** | **EternaFold** | **MXfold2** | **SPOT-RNA** | **RNA-FM** | **RiNALMo** |
|---|---|---|---|---|---|---|---|
| r(CCUG)** | **1.00** | **1.00** | **1.00** | 0.88 | 0.94‡ | **1.00** | 0.65 |
| r(AUUCU) | **0.83** | **0.83** | **0.83** | 0.75 | **0.83** | **0.83** | 0.75 |
| **Miscellaneous synthetics** | | | | | | | |
| Nanoarchitecture 1 | **1.00** | **1.00** | **1.00** | 0.29 | 0.21 | 0.00 | 0.00 |
| Nanoarchitecture 2 | **1.00** | **1.00** | **1.00** | **1.00** | 0.85 | 0.60 | 0.23 |
| Synthetic hairpin | 0.67 | 0.67 | **1.00** | **1.00** | **1.00** | **1.00** | 0.50 |
| Synthetic tetraloop-tetraloop receptor | **1.00** | **1.00** | **1.00** | **1.00** | **1.00**‡ | 0.97 | 0.90 |
| Synthetic G-quadruplex | 0.71 | 0.71 | 0.71 | 0.71 | **1.00**† | 0.71 | 0.57 |
| **Miscellaneous** | | | | | | | |
| ToxI | 0.19 | 0.19 | 0.38 | 0.00 | **0.71**‡ | 0.40 | 0.23 |
| Structured part of acrIF8-aca2 5' UTR | **1.00** | **1.00** | **1.00** | 0.75 | 0.75 | 0.75 | 0.58 |
| Saguaro cactus viral mRNA structure | 0.76 | 0.79 | **0.91** | 0.82 | 0.87 | 0.65 | 0.48 |
| ITS-2 | **0.89** | **0.89** | 0.83 | **0.89** | 0.68 | **0.89** | 0.83 |
| ENE | **1.00** | **1.00** | **1.00** | 0.99 | **1.00**† | 0.92 | 0.82 |
| ASH1 mRNA E3-localization element** | **1.00** | **1.00** | **1.00** | **1.00** | 0.83‡ | **1.00** | 0.75 |
| SCNMV xrRNA | 0.89 | **1.00** | 0.89 | 0.78 | **1.00**‡ | **1.00** | 0.56 |
| Influenza B vRNA promoter | **0.67** | **0.67** | **0.67** | **0.67** | **0.67** | **0.67** | **0.67** |
| NAD-II riboswitch | 0.41 | 0.41 | 0.60 | 0.40 | **0.80** | 0.20 | 0.20 |
| **Descriptive statistics** | | | | | | | |
| Mean | 0.78 | 0.78 | **0.81** | 0.77 | 0.80 | 0.72 | 0.50 |
| Median | 0.83 | 0.83 | 0.84 | 0.81 | **0.84** | 0.73 | 0.54 |
| Min | 0.12 | 0.12 | 0.00 | 0.00 | **0.21** | 0.00 | 0.00 |
| Max | **1.00** | **1.00** | **1.00** | **1.00** | **1.00** | **1.00** | 0.90 |
| Std. Dev. | 0.21 | 0.21 | 0.21 | 0.22 | 0.19 | **0.24** | 0.23 |

**Supplemental Table S2**. **Sensitivity on the proposed benchmarking dataset.** Values with † indicate homology to SPOT-RNA's PDB transfer learning dataset, while values with ‡ indicate exact match between PDB structures.

##### 1.3 F-score

| | RNAstructure | ViennaRNA | EternaFold | MXfold2 | SPOT-RNA | RNA-FM | RiNALMo |
|---|---|---|---|---|---|---|---|
| **Aptamers** | | | | | | | |
| Theophyline aptamer* | **0.87** | **0.87** | 0.85 | 0.73 | 0.77 | **0.87** | 0.69 |
| minE/minF aptamer | 0.69 | 0.69 | **0.82** | 0.75 | 0.78‡ | 0.73 | 0.67 |
| RhoBAST aptamer | 0.88 | **0.91** | 0.88 | 0.90 | 0.76 | 0.84 | 0.73 |
| Spinach aptamer | 0.61 | 0.62 | 0.60 | 0.36 | **0.80**† | 0.50 | 0.43 |
| Mango aptamer | 0.78 | 0.78 | **0.95** | 0.94 | 0.83‡ | 0.92 | 0.74 |
| Vitamin B12 aptamer | 0.13 | 0.13 | 0.46 | 0.50 | **0.74**† | 0.13 | 0.22 |
| Squash aptamer | 0.86 | **0.89** | 0.85 | **0.89** | 0.76 | 0.68 | 0.65 |
| Pepper aptamer | **0.97** | **0.97** | 0.00 | **0.97** | 0.71 | **0.97** | 0.00 |
| Chili aptamer | 0.56 | 0.56 | 0.69 | 0.55 | **0.80** | **0.80** | 0.50 |
| Corn aptamer | 0.56 | 0.56 | 0.53 | 0.59 | **0.78**‡ | 0.56 | 0.53 |
| DIR2s aptamer | 0.83 | **0.94** | 0.77 | 0.76 | 0.89‡ | 0.81 | 0.79 |
| Clivia aptamer | 0.63 | 0.68 | 0.83 | **1.00** | 0.76 | 0.89 | 0.64 |
| A9g aptamer | **0.90** | 0.50 | 0.57 | 0.50 | 0.48 | 0.60 | 0.67 |
| Beetroot aptamer | **0.88** | **0.88** | **0.88** | **0.88** | 0.52 | 0.70 | 0.47 |
| Malachite green aptamer** | **1.00** | **1.00** | **1.00** | **1.00** | 0.88‡ | 0.87 | 0.87 |
| 11F7t aptamer | 0.86 | 0.86 | **1.00** | 0.86 | 0.92‡ | 0.95 | 0.53 |
| K1 aptamer | **1.00** | **1.00** | **1.00** | **1.00** | 0.82‡ | **1.00** | 0.80 |
| Tetracycline aptamer | 0.94 | 0.91 | 0.94 | **0.97** | 0.94‡ | 0.84 | 0.76 |
| **CRISPR RNA guides** | | | | | | | |
| Cas9 guide | 0.89 | 0.89 | 0.94 | **0.95** | 0.83 | 0.90 | 0.42 |
| Cas12 guide | 0.57 | 0.59 | 0.59 | 0.60 | 0.58 | **0.64** | 0.47 |
| Cas13 guide | 0.53 | 0.53 | 0.52 | **0.55** | 0.45‡ | 0.50 | 0.25 |
| **IRES** | | | | | | | |
| IAPV IRES* | 0.56 | 0.56 | 0.54 | 0.58 | **0.58**† | 0.57 | 0.32 |
| CrPV 5'UTR IRES | 0.31 | 0.29 | 0.38 | 0.31 | 0.23† | **0.41** | 0.17 |
| TSV IRES | 0.35 | 0.35 | 0.57 | 0.57 | **0.60** | 0.56 | 0.44 |
| PSIV IGR IRES | 0.56 | 0.56 | 0.72 | 0.72 | **0.90**‡ | 0.72 | 0.67 |
| **Ribozymes** | | | | | | | |
| Synthetic ligase ribozyme | 0.77 | 0.80 | 0.81 | 0.80 | 0.73‡ | **0.81** | 0.55 |
| VS ribozyme** | 0.61 | 0.92 | 0.86 | **0.94** | 0.87† | 0.51 | 0.28 |
| Self-alkylating ribozyme | **0.97** | **0.97** | 0.83 | **0.97** | 0.73 | 0.75 | 0.14 |
| Diels-Alder ribozyme | **1.00** | 0.94 | 0.94 | **1.00** | 0.84 | 0.94 | 0.88 |
| Methyltransferase ribozyme | 0.93 | 0.84 | **0.96** | 0.91 | 0.90 | 0.85 | 0.63 |
| **Disease-associated nucleotide repeats** | | | | | | | |
| r(CUG)** | 0.91 | 0.91 | **0.96** | **0.96** | 0.86 | **0.96** | 0.80 |

Continued on next page

| | **RNAstructure** | **ViennaRNA** | **EternaFold** | **MXfold2** | **SPOT-RNA** | **RNA-FM** | **RiNALMo** |
|---|---|---|---|---|---|---|---|
| r(CCUG)** | **1.00** | **1.00** | **1.00** | 0.94 | 0.86‡ | **1.00** | 0.73 |
| r(AUUCU) | **0.91** | **0.91** | **0.91** | 0.86 | 0.80 | **0.91** | 0.86 |
| **Miscellaneous synthetics** | | | | | | | |
| Nanoarchitecture 1 | 0.93 | 0.93 | **0.97** | 0.31 | 0.21 | 0.00 | 0.00 |
| Nanoarchitecture 2 | **0.98** | **0.98** | **0.98** | **0.98** | 0.79 | 0.63 | 0.30 |
| Synthetic hairpin | 0.80 | 0.80 | **1.00** | **1.00** | 0.86 | **1.00** | 0.67 |
| Synthetic tetraloop-tetraloop receptor | 0.92 | 0.92 | 0.95 | **0.97** | 0.94‡ | 0.95 | 0.92 |
| Synthetic G-quadruplex | 0.67 | 0.67 | 0.67 | 0.77 | **0.78**† | 0.67 | 0.62 |
| **Miscellaneous** | | | | | | | |
| ToxI | 0.22 | 0.22 | 0.44 | 0.00 | **0.64**‡ | 0.41 | 0.25 |
| Structured part of acrIF8-aca2 5' UTR | **0.96** | **0.96** | **0.96** | 0.75 | 0.86 | 0.86 | 0.74 |
| Saguaro cactus viral mRNA structure | 0.79 | 0.81 | **0.93** | 0.86 | 0.87 | 0.70 | 0.61 |
| ITS-2 | 0.68 | 0.70 | 0.67 | 0.75 | 0.55 | 0.73 | **0.82** |
| ENE | 0.97 | 0.97 | 0.95 | **0.98** | 0.95† | 0.95 | 0.89 |
| ASH1 mRNA E3-localization element** | **0.89** | **0.89** | **0.89** | **0.89** | 0.71‡ | **0.89** | 0.72 |
| SCNMV xrRNA | 0.94 | **1.00** | 0.80 | 0.78 | 0.86‡ | 0.95 | 0.67 |
| Influenza B vRNA promoter | **0.47** | **0.47** | 0.40 | **0.47** | 0.42 | 0.42 | 0.44 |
| NAD-II riboswitch | 0.42 | 0.42 | 0.64 | 0.44 | **0.83** | 0.22 | 0.26 |
| **Descriptive statistics** | | | | | | | |
| Mean | 0.75 | 0.76 | **0.77** | 0.76 | 0.74 | 0.72 | 0.56 |
| Median | **0.86** | **0.86** | 0.85 | 0.86 | 0.79 | 0.80 | 0.63 |
| Min | 0.13 | 0.13 | 0.00 | 0.00 | **0.21** | 0.00 | 0.00 |
| Max | **1.00** | **1.00** | **1.00** | **1.00** | 0.95 | **1.00** | 0.92 |
| Std. Dev. | 0.23 | 0.23 | 0.22 | 0.23 | 0.17 | 0.24 | **0.24** |

**Supplemental Table S3**. **F-score on the proposed benchmarking dataset.** Values with † indicate homology to SPOT-RNA's PDB transfer learning dataset, while values with ‡ indicate exact match between PDB structures.

#### 1.4 Homology filtered results

| | **RNAstructure** | **ViennaRNA** | **EternaFold** | **MXfold2** | **SPOT-RNA** | **RNA-FM** | **RiNALMo** |
|---|---|---|---|---|---|---|---|
| Mean | 0.741 | 0.731 | 0.742 | **0.754** | 0.701 | 0.732 | 0.665 |
| Median | 0.841 | **0.853** | 0.826 | 0.841 | 0.760 | 0.809 | 0.691 |
| Min | 0.143 | 0.143 | 0.000 | 0.000 | **0.200** | 0.000 | 0.000 |
| Max | **1.000** | **1.000** | **1.000** | **1.000** | **1.000** | **1.000** | **1.000** |
| Std. Dev. | 0.257 | 0.258 | 0.239 | 0.261 | 0.205 | 0.265 | **0.288** |

**Supplemental Table S4**. **Descriptive statistics of PPV on the benchmarking dataset with all homology removed.** All overlaps found in Table S8 (*, **) and Table S9 (†, ‡) are excluded from this calculation.

| | **RNAstructure** | **ViennaRNA** | **EternaFold** | **MXfold2** | **SPOT-RNA** | **RNA-FM** | **RiNALMo** |
|---|---|---|---|---|---|---|---|
| Mean | 0.758 | 0.757 | **0.792** | 0.746 | 0.726 | 0.708 | 0.488 |
| Median | 0.804 | 0.821 | **0.833** | 0.765 | 0.734 | 0.718 | 0.500 |
| Min | 0.125 | 0.125 | 0.000 | 0.000 | **0.214** | 0.000 | 0.000 |
| Max | **1.000** | **1.000** | **1.000** | **1.000** | **1.000** | **1.000** | 0.900 |
| Std. Dev. | 0.219 | 0.220 | 0.215 | 0.231 | 0.181 | **0.238** | 0.236 |

**Supplemental Table S5**. **Descriptive statistics of sensitivity on the benchmarking dataset with all homology removed.** All overlaps found in Table S8 (*, **) and Table S9 (†, ‡) are excluded from this calculation.

| | RNAstructure | ViennaRNA | EternaFold | MXfold2 | SPOT-RNA | RNA-FM | RiNALMo |
|---|---|---|---|---|---|---|---|
| Mean | 0.740 | 0.735 | **0.758** | 0.742 | 0.702 | 0.710 | 0.544 |
| Median | 0.817 | 0.807 | **0.827** | 0.787 | 0.761 | 0.741 | 0.612 |
| Min | 0.133 | 0.133 | 0.000 | 0.000 | **0.207** | 0.000 | 0.000 |
| Max | **1.000** | **1.000** | **1.000** | **1.000** | 0.903 | **1.000** | 0.915 |
| Std. Dev. | 0.232 | 0.235 | 0.225 | 0.243 | 0.178 | 0.241 | **0.243** |

**Supplemental Table S6. Descriptive statistics of F-score on the benchmarking dataset with all homology removed.** All overlaps found in Table **S8** (*, **) and Table **S9** (†, ‡) are excluded from this calculation.

#### 1.5 Statistical testing

| | ViennaRNA | EternaFold | MXfold2 | SPOT-RNA | RNA-FM | RiNALMo |
|---|---|---|---|---|---|---|
| **RNAstructure** | 6.43e-01 | 5.92e-01 | 9.59e-01 | 1.15e-01 | 3.45e-01 | 2.82e-05 |
| **ViennaRNA** | | 4.75e-01 | 8.00e-01 | 1.80e-01 | 4.24e-01 | 5.42e-05 |
| **EternaFold** | | | 6.51e-01 | 2.08e-01 | 2.35e-01 | 4.02e-07 |
| **MXfold2** | | | | 5.28e-02 | 1.94e-01 | 6.55e-06 |
| **SPOT-RNA** | | | | | 8.68e-01 | 2.19e-03 |
| **RNA-FM** | | | | | | 6.83e-06 |

**Supplemental Table S7. The $p$-values from paired $t$-tests between F-score of the different methods.** All overlaps found in Table **S8** (*, **) and Table **S9** (†, ‡) are excluded from this calculation. Note that there are no corrections, so the values must be carefully interpreted to avoid the multiple testing problem. Red values are statistically significant ($p < 0.05$).

#### 2 bpRNA-1m PDB set homology

| | bpRNA-1m Entry |
|---|---|
| **Aptamers** | |
| Theophyline aptamer | 1eht_A, 1o15_A |
| minE/minF aptamer | |
| RhoBAST aptamer | |
| Spinach aptamer | |
| Mango aptamer | |
| Vitamin B12 aptamer | |
| Squash aptamer | |
| Pepper aptamer | |
| Chili aptamer | |
| Corn aptamer | |
| DIR2s aptamer | |
| Clivia aptamer | |
| A9g aptamer | |
| Beetroot aptamer | |
| Malachite green aptamer | 1f1t_A**, 1q8n_A |
| 11F7t aptamer | |
| K1 aptamer | |
| Tetracycline aptamer | |
| **CRISPR RNA guides** | |
| Cas9 guide | |
| Cas12 guide | |
| Cas13 guide | |
| **IRES** | |
| IAPV IRES | 2n8v_X |
| CrPV 5'UTR IRES | |
| TSV IRES | |
| PSIV IGR IRES | |
| **Ribozymes** | |
| Synthetic ligase ribozyme | |
| VS ribozyme | 4r4v_A**, 2n3r_A, 2mtj_A, 1tbk_A, 1ow9_A, 2mtk_A, 2mis_A, 1tjz_A, 2n3q_A, 4r4p_A, 2l5z_A, 1yn1_A, 1hwq_A, 1yn2_A |
| Self-alkylating ribozyme | |
| Diels-Alder ribozyme | |
| Methyltransferase ribozyme | |
| **Disease-associated nucleotide repeats** | |
| r(CUG) | 4pcj_A**, 4fnj_A |

Continued on next page

| | bpRNA-1m Entry |
|---|---|
| r(CCUG) | 4k27_U** |
| r(AUUCU) | |
| **Miscellaneous synthetics** | |
| Nanoarchitecture 1 | |
| Nanoarchitecture 2 | |
| Synthetic hairpin | |
| Synthetic tetraloop-tetraloop receptor | |
| Synthetic G-quadruplex | |
| **Miscellaneous** | |
| ToxI | |
| Structured part of acrIF8-aca2 5' UTR | |
| Saguaro cactus viral mRNA structure | |
| ITS-2 | |
| ENE | |
| ASH1 mRNA E3-localization element | 5m0h_A** |
| SCNMV xrRNA | |
| Influenza B vRNA promoter | |
| NAD-II riboswitch | |

**Supplemental Table S8**. **Overlapping PDB IDs between our benchmarking dataset and bpRNA-1m's PDB dataset.** Entries with ** indicate an exact match (i.e. exact PDB chain are present in both datasets), while regular entries indicate related structures found during manual curation. Note that author chain IDs are added (after underscore) to the PDB ID.

### 3 SPOT-RNA transfer learning PDB set homology

| | SPOT-RNA Entry |
|---|---|
| **Aptamers** | |
| Theophyline aptamer | |
| minE/minF aptamer | 4m4o_B‡ |
| RhoBAST aptamer | |
| Spinach aptamer | 4kzd_R |
| Mango aptamer | 6e8s_B‡, 6c63_A |
| Vitamin B12 aptamer | 1ddy_A |
| Squash aptamer | |
| Pepper aptamer | |
| Chili aptamer | |
| Corn aptamer | 5bjo_Y‡ |
| DIR2s aptamer | 6db8_R‡ |
| Clivia aptamer | |
| A9g aptamer | |
| Beetroot aptamer | |
| Malachite green aptamer | 1f1t_A‡, 6gzk_A |
| 11F7t aptamer | 5voe_A‡ |
| K1 aptamer | 6sy4_C‡ |
| Tetracycline aptamer | 3egz_B‡ |
| **CRISPR RNA guides** | |
| Cas9 guide | |
| Cas12 guide | |
| Cas13 guide | 6dtd_C‡, 6aay_B‡ |
| **IRES** | |
| IAPV IRES | 2n8v_X |
| CrPV 5'UTR IRES | 4v84_AV |
| TSV IRES | |
| PSIV IGR IRES | 4v83_CV‡ |
| **Ribozymes** | |
| Synthetic ligase ribozyme | 3hhn_E‡ |
| VS ribozyme | 4r4p_A, 2mtj_A |
| Self-alkylating ribozyme | |
| Diels-Alder ribozyme | |
| Methyltransferase ribozyme | |
| **Disease-associated nucleotide repeats** | |
| r(CUG) | |
| r(CCUG) | 4k27_U‡ |
| r(AUUCU) | |
| **Miscellaneous synthetics** | |
| Nanoarchitecture 1 | |

Continued on next page

| | SPOT-RNA Entry |
|---|---|
| Nanoarchitecture 2 | |
| Synthetic hairpin | |
| Synthetic tetraloop-tetraloop receptor | 6dvk_H‡ |
| Synthetic G-quadruplex | 5de5_A |
| **Miscellaneous** | |
| ToxI | 2xd0_G, 4ato_G, 4rmo_H‡ |
| Structured part of acrIF8-aca2 5' UTR | |
| Saguaro cactus viral mRNA structure | |
| ITS-2 | |
| ENE | 4plx_A |
| ASH1 mRNA E3-localization element | 5m0h_A‡ |
| SCNMV xrRNA | 6d3p_A‡ |
| Influenza B vRNA promoter | |
| NAD-II riboswitch | |

**Supplemental Table S9**. **Overlapping PDB IDs between our benchmarking dataset and SPOT-RNA's transfer learning dataset.** Entries with ‡ indicate an exact match (i.e. exact PDB chain are present in both datasets), while regular entries indicate related structures found during manual curation. Note that author chain IDs are added (after underscore) to the PDB ID.

### 4 SPOT-RNA & SPOT-RNA 2 results

#### 4.1 Positive Predictive Value

| | RNAstructure | ViennaRNA | SPOT-RNA | SPOT-RNA 2 |
|---|---|---|---|---|
| **Aptamers** | | | | |
| Theophyline aptamer* | **0.80** | **0.80** | 0.64 | 0.62 |
| minE/minF aptamer | **0.75** | **0.75** | 0.68‡ | 0.68 |
| RhoBAST aptamer | **1.00** | **1.00** | 0.81 | 0.81 |
| Spinach aptamer | 0.61 | 0.59 | **0.75**† | 0.65 |
| Mango aptamer | **0.73** | **0.73** | 0.73‡ | 0.71 |
| Vitamin B12 aptamer | 0.14 | 0.14 | **0.64**† | **0.64** |
| Squash aptamer | 0.84 | **0.91** | 0.89 | 0.57 |
| Pepper aptamer | **1.00** | **1.00** | 0.83 | 0.83 |
| Chili aptamer | 0.53 | 0.53 | **0.80** | **0.80** |
| Corn aptamer | 0.45 | 0.45 | **0.64**‡ | **0.64** |
| DIR2s aptamer | 0.88 | **1.00** | 0.94‡ | 0.89 |
| Clivia aptamer | 0.55 | 0.57 | **0.71** | 0.69 |
| A9g aptamer | **0.82** | 0.45 | 0.42 | 0.42 |
| Beetroot aptamer | **0.92** | **0.92** | 0.61 | 0.79 |
| Malachite green aptamer** | **1.00** | **1.00** | 0.92‡ | 0.85 |
| 11F7t aptamer | **0.90** | **0.90** | 0.85‡ | 0.00 |
| K1 aptamer | **1.00** | **1.00** | 0.70‡ | 0.70 |
| Tetracycline aptamer | **1.00** | 0.94 | **1.00**‡ | 0.89 |
| **CRISPR RNA guides** | | | | |
| Cas9 guide | **0.87** | 0.87 | 0.78 | 0.82 |
| Cas12 guide | 0.56 | 0.58 | **0.61** | 0.54 |
| Cas13 guide | 0.48 | **0.49** | 0.39‡ | 0.42 |
| **IRES** | | | | |
| IAPV IRES* | 0.46 | 0.45 | **0.52**† | nan |
| CrPV 5'UTR IRES | **0.22** | 0.21 | 0.22† | nan |
| TSV IRES | 0.29 | 0.29 | **0.56** | nan |
| PSIV IGR IRES | 0.64 | 0.64 | **0.82**‡ | 0.81 |
| **Ribozymes** | | | | |
| Synthetic ligase ribozyme | 0.84 | **0.85** | 0.76‡ | 0.73 |
| VS ribozyme** | 0.62 | 0.90 | **0.93**† | nan |
| Self-alkylating ribozyme | **1.00** | **1.00** | 0.80 | 0.80 |
| Diels-Alder ribozyme | **1.00** | 0.89 | 0.73 | 0.70 |
| Methyltransferase ribozyme | **0.97** | 0.87 | 0.93 | 0.85 |

Continued on next page

| | **RNAstructure** | **ViennaRNA** | **SPOT-RNA** | **SPOT-RNA 2** |
|---|---|---|---|---|
| **Disease-associated nucleotide repeats** | | | | |
| r(CUG)** | **1.00** | **1.00** | 0.75 | 0.75 |
| r(CCUG)** | **1.00** | **1.00** | 0.80‡ | 0.77 |
| r(AUUCU) | **1.00** | **1.00** | 0.77 | 0.77 |
| **Miscellaneous synthetics** | | | | |
| Nanoarchitecture 1 | **0.88** | **0.88** | 0.20 | 0.20 |
| Nanoarchitecture 2 | **0.95** | **0.95** | 0.74 | 0.74 |
| Synthetic hairpin | **1.00** | **1.00** | 0.75 | 0.86 |
| Synthetic tetraloop-tetraloop receptor | 0.86 | 0.86 | **0.88**‡ | **0.88** |
| Synthetic G-quadruplex | 0.62 | 0.62 | **0.64**† | 0.42 |
| **Miscellaneous** | | | | |
| ToxI | 0.27 | 0.27 | **0.58**‡ | 0.57 |
| Structured part of acrIF8-aca2 5' UTR | 0.92 | 0.92 | **1.00** | **1.00** |
| Saguaro cactus viral mRNA structure | 0.83 | 0.84 | **0.86** | **0.86** |
| ITS-2 | 0.56 | **0.58** | 0.46 | 0.46 |
| ENE | **0.95** | **0.95** | 0.90† | 0.89 |
| ASH1 mRNA E3-localization element** | **0.80** | **0.80** | 0.62‡ | 0.00 |
| SCNMV xrRNA | **1.00** | **1.00** | 0.75‡ | 0.75 |
| Influenza B vRNA promoter | **0.36** | **0.36** | 0.31 | 0.31 |
| NAD-II riboswitch | 0.44 | 0.44 | **0.87** | 0.79 |
| **Descriptive statistics** | | | | |
| Mean | **0.75** | 0.75 | 0.71 | 0.67 |
| Median | 0.84 | **0.86** | 0.75 | 0.74 |
| Min | 0.14 | 0.14 | **0.20** | 0.00 |
| Max | **1.00** | **1.00** | **1.00** | **1.00** |
| Std. Dev. | 0.25 | **0.25** | 0.19 | 0.23 |

**Supplemental Table S10**. **Positive Predictive Value (PPV) on the proposed benchmarking dataset for SPOT-RNA and SPOT-RNA 2.** Values with † indicate homology to SPOT-RNA's PDB transfer learning dataset, while values with ‡ indicate exact match between PDB structures. § indicates structures that we were unable to predict with SPOT-RNA 2 (see *Materials and Methods* for mode information).

#### 4.2 Sensitivity

| | RNAstructure | ViennaRNA | SPOT-RNA | SPOT-RNA 2 |
|---|---|---|---|---|
| **Aptamers** | | | | |
| Theophyline aptamer* | 0.97 | 0.97 | **1.00** | **1.00** |
| minE/minF aptamer | 0.65 | 0.65 | **0.92**$^{\ddagger}$ | **0.92** |
| RhoBAST aptamer | 0.78 | **0.83** | 0.72 | 0.72 |
| Spinach aptamer | 0.62 | 0.64 | **0.86**$^{\dagger}$ | 0.74 |
| Mango aptamer | 0.83 | 0.83 | **0.97**$^{\ddagger}$ | **0.97** |
| Vitamin B12 aptamer | 0.12 | 0.12 | **0.88**$^{\dagger}$ | **0.88** |
| Squash aptamer | **0.88** | **0.88** | 0.67 | 0.67 |
| Pepper aptamer | **0.94** | **0.94** | 0.62 | **0.94** |
| Chili aptamer | 0.60 | 0.60 | **0.80** | **0.80** |
| Corn aptamer | 0.71 | 0.71 | **1.00**$^{\ddagger}$ | **1.00** |
| DIR2s aptamer | 0.79 | **0.89** | 0.84$^{\ddagger}$ | 0.84 |
| Clivia aptamer | 0.74 | 0.82 | 0.82 | **0.88** |
| A9g aptamer | **1.00** | 0.56 | 0.56 | 0.56 |
| Beetroot aptamer | **0.85** | **0.85** | 0.46 | 0.65 |
| Malachite green aptamer** | **1.00** | **1.00** | 0.85$^{\ddagger}$ | 0.85 |
| 11F7t aptamer | 0.82 | 0.82 | **1.00**$^{\ddagger}$ | 0.00 |
| K1 aptamer | **1.00** | **1.00** | **1.00**$^{\ddagger}$ | **1.00** |
| Tetracycline aptamer | **0.89** | **0.89** | **0.89**$^{\ddagger}$ | **0.89** |
| **CRISPR RNA guides** | | | | |
| Cas9 guide | 0.90 | 0.91 | 0.88 | **0.96** |
| Cas12 guide | 0.58 | **0.60** | 0.57 | 0.57 |
| Cas13 guide | 0.60 | 0.60 | 0.54$^{\ddagger}$ | **0.62** |
| **IRES** | | | | |
| IAPV IRES* | **0.74** | **0.74** | 0.68$^{\dagger}$ | nan |
| CrPV 5'UTR IRES | **0.52** | **0.52** | 0.25$^{\dagger}$ | nan |
| TSV IRES | 0.44 | 0.44 | **0.64** | nan |
| PSIV IGR IRES | 0.50 | 0.50 | **1.00**$^{\ddagger}$ | 0.93 |
| **Ribozymes** | | | | |
| Synthetic ligase ribozyme | 0.71 | **0.75** | 0.69$^{\ddagger}$ | 0.74 |
| VS ribozyme** | 0.59 | **0.93** | 0.82$^{\dagger}$ | nan |
| Self-alkylating ribozyme | **0.94** | **0.94** | 0.67 | 0.67 |
| Diels-Alder ribozyme | **1.00** | **1.00** | **1.00** | **1.00** |
| Methyltransferase ribozyme | **0.88** | 0.82 | **0.88** | 0.82 |
| **Disease-associated nucleotide repeats** | | | | |
| r(CUG)** | 0.83 | 0.83 | **1.00** | **1.00** |

Continued on next page

| | RNAstructure | ViennaRNA | SPOT-RNA | SPOT-RNA 2 |
|---|---|---|---|---|
| r(CCUG)** | **1.00** | **1.00** | 0.94‡ | **1.00** |
| r(AUUCU) | **0.83** | **0.83** | **0.83** | **0.83** |
| **Miscellaneous synthetics** | | | | |
| Nanoarchitecture 1 | **1.00** | **1.00** | 0.21 | 0.21 |
| Nanoarchitecture 2 | **1.00** | **1.00** | 0.85 | 0.85 |
| Synthetic hairpin | 0.67 | 0.67 | **1.00** | **1.00** |
| Synthetic tetraloop-tetraloop receptor | **1.00** | **1.00** | **1.00**‡ | **1.00** |
| Synthetic G-quadruplex | 0.71 | 0.71 | **1.00**† | 0.71 |
| **Miscellaneous** | | | | |
| ToxI | 0.19 | 0.19 | **0.71**‡ | 0.63 |
| Structured part of acrIF8-aca2 5' UTR | **1.00** | **1.00** | 0.75 | 0.75 |
| Saguaro cactus viral mRNA structure | 0.76 | 0.79 | **0.87** | **0.87** |
| ITS-2 | **0.89** | **0.89** | 0.68 | 0.68 |
| ENE | **1.00** | **1.00** | **1.00**† | **1.00** |
| ASH1 mRNA E3-localization element** | **1.00** | **1.00** | 0.83‡ | 0.00 |
| SCNMV xrRNA | 0.89 | **1.00** | **1.00**‡ | **1.00** |
| Influenza B vRNA promoter | **0.67** | **0.67** | **0.67** | **0.67** |
| NAD-II riboswitch | 0.41 | 0.41 | **0.80** | **0.80** |
| **Descriptive statistics** | | | | |
| Mean | 0.78 | 0.78 | **0.80** | 0.78 |
| Median | 0.83 | 0.83 | **0.84** | **0.84** |
| Min | 0.12 | 0.12 | **0.21** | 0.00 |
| Max | **1.00** | **1.00** | **1.00** | **1.00** |
| Std. Dev. | 0.21 | 0.21 | 0.19 | **0.24** |

**Supplemental Table S11**. **Sensitivity on the proposed benchmarking dataset for SPOT-RNA and SPOT-RNA 2.** Values with † indicate homology to SPOT-RNA's PDB transfer learning dataset, while values with ‡ indicate exact match between PDB structures. § indicates structures that we were unable to predict with SPOT-RNA 2 (see *Materials and Methods* for mode information).

##### 4.3 F-score

| | RNAstructure | ViennaRNA | SPOT-RNA | SPOT-RNA 2 |
|---|---|---|---|---|
| **Aptamers** | | | | |
| Theophyline aptamer* | **0.87** | **0.87** | 0.77 | 0.76 |
| minE/minF aptamer | 0.69 | 0.69 | **0.78**‡ | **0.78** |
| RhoBAST aptamer | 0.88 | **0.91** | 0.76 | 0.76 |
| Spinach aptamer | 0.61 | 0.62 | **0.80**† | 0.69 |
| Mango aptamer | 0.78 | 0.78 | **0.83**‡ | 0.81 |
| Vitamin B12 aptamer | 0.13 | 0.13 | **0.74**† | **0.74** |
| Squash aptamer | 0.86 | **0.89** | 0.76 | 0.62 |
| Pepper aptamer | **0.97** | **0.97** | 0.71 | 0.88 |
| Chili aptamer | 0.56 | 0.56 | **0.80** | **0.80** |
| Corn aptamer | 0.56 | 0.56 | **0.78**‡ | **0.78** |
| DIR2s aptamer | 0.83 | **0.94** | 0.89‡ | 0.86 |
| Clivia aptamer | 0.63 | 0.68 | 0.76 | **0.77** |
| A9g aptamer | **0.90** | 0.50 | 0.48 | 0.48 |
| Beetroot aptamer | **0.88** | **0.88** | 0.52 | 0.71 |
| Malachite green aptamer** | **1.00** | **1.00** | 0.88‡ | 0.85 |
| 11F7t aptamer | 0.86 | 0.86 | **0.92**‡ | 0.00 |
| K1 aptamer | **1.00** | **1.00** | 0.82‡ | 0.82 |
| Tetracycline aptamer | **0.94** | 0.91 | **0.94**‡ | 0.89 |
| **CRISPR RNA guides** | | | | |
| Cas9 guide | 0.89 | **0.89** | 0.83 | 0.88 |
| Cas12 guide | 0.57 | **0.59** | 0.58 | 0.54 |
| Cas13 guide | 0.53 | **0.53** | 0.45‡ | 0.50 |
| **IRES** | | | | |
| IAPV IRES* | 0.56 | 0.56 | **0.58**† | nan |
| CrPV 5'UTR IRES | **0.31** | 0.29 | 0.23† | nan |
| TSV IRES | 0.35 | 0.35 | **0.60** | nan |
| PSIV IGR IRES | 0.56 | 0.56 | **0.90**‡ | 0.87 |
| **Ribozymes** | | | | |
| Synthetic ligase ribozyme | 0.77 | **0.80** | 0.73‡ | 0.73 |
| VS ribozyme** | 0.61 | **0.92** | 0.87† | nan |
| Self-alkylating ribozyme | **0.97** | **0.97** | 0.73 | 0.73 |
| Diels-Alder ribozyme | **1.00** | 0.94 | 0.84 | 0.82 |
| Methyltransferase ribozyme | **0.93** | 0.84 | 0.90 | 0.83 |
| **Disease-associated nucleotide repeats** | | | | |
| r(CUG)** | **0.91** | **0.91** | 0.86 | 0.86 |

Continued on next page

| | RNAstructure | ViennaRNA | SPOT-RNA | SPOT-RNA 2 |
|---|---|---|---|---|
| r(CCUG)** | **1.00** | **1.00** | 0.86‡ | 0.87 |
| r(AUUCU) | **0.91** | **0.91** | 0.80 | 0.80 |
| **Miscellaneous synthetics** | | | | |
| Nanoarchitecture 1 | **0.93** | **0.93** | 0.21 | 0.21 |
| Nanoarchitecture 2 | **0.98** | **0.98** | 0.79 | 0.79 |
| Synthetic hairpin | 0.80 | 0.80 | 0.86 | **0.92** |
| Synthetic tetraloop-tetraloop receptor | 0.92 | 0.92 | **0.94**‡ | **0.94** |
| Synthetic G-quadruplex | 0.67 | 0.67 | **0.78**† | 0.53 |
| **Miscellaneous** | | | | |
| ToxI | 0.22 | 0.22 | **0.64**‡ | 0.60 |
| Structured part of acrIF8-aca2 5' UTR | **0.96** | **0.96** | 0.86 | 0.86 |
| Saguaro cactus viral mRNA structure | 0.79 | 0.81 | **0.87** | **0.87** |
| ITS-2 | 0.68 | **0.70** | 0.55 | 0.55 |
| ENE | **0.97** | **0.97** | 0.95† | 0.94 |
| ASH1 mRNA E3-localization element** | **0.89** | **0.89** | 0.71‡ | 0.00 |
| SCNMV xrRNA | 0.94 | **1.00** | 0.86‡ | 0.86 |
| Influenza B vRNA promoter | **0.47** | **0.47** | 0.42 | 0.42 |
| NAD-II riboswitch | 0.42 | 0.42 | **0.83** | 0.79 |
| **Descriptive statistics** | | | | |
| Mean | 0.75 | **0.76** | 0.74 | 0.71 |
| Median | **0.86** | **0.86** | 0.79 | 0.79 |
| Min | 0.13 | 0.13 | **0.21** | 0.00 |
| Max | **1.00** | **1.00** | 0.95 | 0.94 |
| Std. Dev. | 0.23 | **0.23** | 0.17 | 0.22 |

**Supplemental Table S12**. **F-score on the proposed benchmarking dataset for SPOT-RNA and SPOT-RNA 2.** Values with † indicate homology to SPOT-RNA's PDB transfer learning dataset, while values with ‡ indicate exact match between PDB structures. § indicates structures that we were unable to predict with SPOT-RNA 2 (see *Materials and Methods* for mode information).

#### 4.4 Homology filtered results

| | **RNAstructure** | **ViennaRNA** | **SPOT-RNA** | **SPOT-RNA 2** |
|---|---|---|---|---|
| Mean | **0.741** | 0.731 | 0.701 | 0.681 |
| Median | 0.841 | **0.853** | 0.760 | 0.736 |
| Min | 0.143 | 0.143 | **0.200** | 0.000 |
| Max | **1.000** | **1.000** | **1.000** | **1.000** |
| Std. Dev. | 0.257 | **0.258** | 0.205 | 0.212 |

**Supplemental Table S13**. **Descriptive statistics of PPV on the benchmarking dataset with all homology removed for SPOT-RNA and SPOT-RNA 2.** All overlaps found in Table **S8** (*, **) and Table **S9** (†, ‡) are excluded from this calculation.

| | **RNAstructure** | **ViennaRNA** | **SPOT-RNA** | **SPOT-RNA 2** |
|---|---|---|---|---|
| Mean | 0.758 | 0.757 | 0.726 | **0.783** |
| Median | 0.804 | 0.821 | 0.734 | **0.825** |
| Min | 0.125 | 0.125 | **0.214** | 0.000 |
| Max | **1.000** | **1.000** | **1.000** | **1.000** |
| Std. Dev. | 0.219 | **0.220** | 0.181 | 0.213 |

**Supplemental Table S14**. **Descriptive statistics of sensitivity on the benchmarking dataset with all homology removed for SPOT-RNA and SPOT-RNA 2.** All overlaps found in Table **S8** (*, **) and Table **S9** (†, ‡) are excluded from this calculation.

| | RNAstructure | ViennaRNA | SPOT-RNA | SPOT-RNA 2 |
|---|---|---|---|---|
| Mean | **0.740** | 0.735 | 0.702 | 0.720 |
| Median | **0.817** | 0.807 | 0.761 | 0.782 |
| Min | 0.133 | 0.133 | **0.207** | 0.000 |
| Max | **1.000** | **1.000** | 0.903 | 0.942 |
| Std. Dev. | 0.232 | **0.235** | 0.178 | 0.202 |

**Supplemental Table S15**. **Descriptive statistics of F-score on the benchmarking dataset with all homology removed for SPOT-RNA and SPOT-RNA 2.** All overlaps found in Table **S8** (*, **) and Table **S9** (†, ‡) are excluded from this calculation.

#### 4.5 Statistical testing

| | ViennaRNA | SPOT-RNA | SPOT-RNA 2 |
|---|---|---|---|
| RNAstructure | 6.43e-01 | 1.15e-01 | 3.32e-01 |
| ViennaRNA | | 1.80e-01 | 3.78e-01 |
| SPOT-RNA | | | 6.19e-01 |

**Supplemental Table S16**. **The $p$-values from paired $t$-tests between F-scores of SPOT-RNA and SPOT-RNA 2.** All overlaps found in Table **S8** (*, **) and Table **S9** (†, ‡) are excluded from this calculation. Note that there are no corrections, so the values must be carefully interpreted to avoid the multiple testing problem. No values are statistically significant ($p < 0.05$).

### 5 Models for probing data prediction

| **Dataset** | **MAE** |
|---|---|
| SARS-CoV-2 | 0.177 |
| HIV-1 | 0.201 |
| *E. coli* | 0.235 |
| *Homo Sapiens* | 0.183 |

**Supplemental Table S17**. **Mean Absolute Error (MAE) for each dataset.** MAE values were computed across all nucleotides in a dataset. Sequences were not computationally or experimentally padded, and no flanking sequences or barcodes were included.

### 6 Method identification

#### 6.1 Semantic Scholar

| Title | Authors | Year | DOI | Citations | Citations/Year | Relevance | Method |
|---|---|---|---|---|---|---|---|
| RNA secondary structure prediction using deep learning with thermodynamic integration | Sato et al. | 2021 | 10.1038/s41467-021-21194-4 | 290 | 72.50 | ✓ | MXfold2 |
| RiNALMo: general-purpose RNA language models can generalize well on structure prediction tasks | Penić et al. | 2024 | 10.1038/s41467-025-60872-5 | 38 | 38.00 | ✓ | RiNALMo |
| UFold: fast and accurate RNA secondary structure prediction with deep learning | Fu et al. | 2020 | 10.1093/nar/gkab1074 | 169 | 33.80 | ✓ | Ufold |

Continued on next page

| Title | Authors | Year | DOI | Citations | Citations/Year | Relevance | Method |
|---|---|---|---|---|---|---|---|
| Recent trends in RNA informatics: a review of machine learning and deep learning for RNA secondary structure prediction and RNA drug discovery | Sato et al. | 2023 | 10.1093/bib/bbad186 | 48 | 24.00 | ✓ | Review |
| RNA Secondary Structure Prediction By Learning Unrolled Algorithms | Chen et al. | 2020 | 10.48550/arXiv.2002.05810 | 119 | 23.80 | ✓ | E2Efold |
| All-atom RNA structure determination from cryo-EM maps. | Li et al. | 2024 | 10.1038/s41587-024-02149-8 | 23 | 23.00 | ✗ | |
| Informative RNA base embedding for RNA structural alignment and clustering by deep representation learning | Akiyama et al. | 2022 | 10.1093/nargab/lqac012 | 68 | 22.67 | ✗ | |
| Predicting dynamic cellular protein–RNA interactions by deep learning using in vivo RNA structures | Sun et al. | 2021 | 10.1038/s41422-021-00476-y | 83 | 20.75 | ✗ | |

Continued on next page

| Title | Authors | Year | DOI | Citations | Citations/Year | Relevance | Method |
|---|---|---|---|---|---|---|---|
| Improved RNA secondary structure and tertiary base-pairing prediction using evolutionary profile, mutational coupling and two-dimensional transfer learning | Singh et al. | 2021 | 10.1093/bioinformatics/btab165 | 81 | 20.25 | ✓ | SPOT-RNA 2 |
| Deep learning models for RNA secondary structure prediction (probably) do not generalize across families | Szikszai et al. | 2022 | 10.1093/bioinformatics/btac415 | 56 | 18.67 | ✓ | Review |
| Machine learning for RNA 2D structure prediction benchmarked on experimental data | Justyna et al. | 2023 | 10.1093/bib/bbad153 | 32 | 16.00 | ✓ | Review |
| Review of machine learning methods for RNA secondary structure prediction | Zhao et al. | 2020 | 10.1371/journal.pcbi.1009291 | 65 | 13.00 | ✓ | Review |

Continued on next page

| Title | Authors | Year | DOI | Citations | Citations/Year | Relevance | Method |
|---|---|---|---|---|---|---|---|
| REDfold: accurate RNA secondary structure prediction using residual encoder-decoder network | Chen et al. | 2023 | 10.1186/s12859-023-05238-8 | 25 | 12.50 | ✓ | REDfold |
| gRNAde: Geometric Deep Learning for 3D RNA inverse design | Joshi et al. | 2023 | 10.1101/2024.03.31.587283 | 19 | 9.50 | ✗ | |
| Detecting protein and DNA/RNA structures in cryo-EM maps of intermediate resolution using deep learning | Wang et al. | 2021 | 10.1038/s41467-021-22577-3 | 32 | 8.00 | ✗ | |
| Deep learning predicts short non-coding RNA functions from only raw sequence data | Noviello et al. | 2020 | 10.1371/journal.pcbi.1008415 | 33 | 6.60 | ✗ | |

Continued on next page

| Title | Authors | Year | DOI | Citations | Citations/Year | Relevance | Method |
|---|---|---|---|---|---|---|---|
| Predicting RNA distance-based contact maps by integrated deep learning on physics-inferred secondary structure and evolutionary-derived mutational coupling | Singh et al. | 2022 | 10.1093/bioinformatics/btac421 | 18 | 6.00 | ✗ | |
| Caveats to Deep Learning Approaches to RNA Secondary Structure Prediction | Flamm et al. | 2021 | 10.3389/fbinf.2022.835422 | 22 | 5.50 | ✓ | Review |
| Improved Predicting of The Sequence Specificities of RNA Binding Proteins by Deep Learning | Tayara et al. | 2020 | 10.1109/tcbb.2020.2981335 | 22 | 4.40 | ✗ | |

Continued on next page

| Title | Authors | Year | DOI | Citations | Citations/Year | Relevance | Method |
|---|---|---|---|---|---|---|---|
| Deep neural networks for inferring binding sites of RNA-binding proteins by using distributed representations of RNA primary sequence and secondary structure | Deng et al. | 2020 | 10.1186/s12864-020-07239-w | 21 | 4.20 | ✗ | |

**Supplemental Table S18**. **Semantic Scholar method identification.** Top 20 papers on Semantic Scholar sorted by citation count for the search term "deep learning RNA secondary structure" released after 2020 (retrieved 2025-08-27).

#### 6.2 Google Scholar

| Title | Authors | Year | DOI | Citations | Citations/Year | Relevance | Method |
|---|---|---|---|---|---|---|---|
| RNA secondary structure prediction using deep learning with thermodynamic integration | Sato et al. | 2021 | 10.1038/s41467-021-21194-4 | 445 | 111.25 | ✓ | MXfold2 |
| Geometric deep learning of RNA structure | Townshend et al. | 2021 | 10.1126/science.abe5650 | 415 | 103.75 | ✗ | |
| UFold: fast and accurate RNA secondary structure prediction with deep learning | Fu et al. | 2022 | 10.1093/nar/gkab1074 | 237 | 79.00 | ✓ | Ufold |
| Advances and opportunities in RNA structure experimental determination and computational modeling | Zhang et al. | 2022 | 10.1038/s41592-022-01623-y | 141 | 47.00 | ✓ | Review |

Continued on next page

| **Title** | **Authors** | **Year** | **DOI** | **Citations** | **Citations/Year** | **Relevance** | **Method** |
|---|---|---|---|---|---|---|---|
| Integrating end-to-end learning with deep geometrical potentials for ab initio RNA structure prediction | Li et al. | 2023 | 10.1038/s41467-023-41303-9 | 83 | 41.50 | ✗ | |
| RNA secondary structure packages evaluated and improved by high-throughput experiments | Wayment-Steele et al. | 2022 | 10.1038/s41592-022-01605-0 | 117 | 39.00 | ✓ | Review |
| Predicting dynamic cellular protein–RNA interactions by deep learning using in vivo RNA structures | Sun et al. | 2021 | 10.1038/s41422-021-00476-y | 131 | 32.75 | ✗ | |
| Recent trends in RNA informatics: a review of machine learning and deep learning for RNA secondary structure prediction and RNA drug discovery | Sato et al. | 2023 | 10.1093/bib/bbad186 | 63 | 31.50 | ✓ | Review |

Continued on next page

| Title | Authors | Year | DOI | Citations | Citations/Year | Relevance | Method |
|---|---|---|---|---|---|---|---|
| RNA secondary structure prediction by learning unrolled algorithms | Chen et al. | 2020 | 10.48550/arXiv.2002.05810 | 155 | 31.00 | ✓ | E2Efold |
| De Novo RNA Tertiary Structure Prediction at Atomic Resolution Using Geometric Potentials from Deep Learning | Pearce et al. | 2022 | 10.1101/2022.05.15.491755 | 89 | 29.67 | ✗ | |
| A deep learning approach to programmable RNA switches | Angenent-Mari et al. | 2020 | 10.1038/s41467-020-18677-1 | 148 | 29.60 | ✗ | |
| Protein–RNA interaction prediction with deep learning: structure matters | Wei et al. | 2022 | 10.1093/bib/bbab540 | 83 | 27.67 | ✗ | |
| Review of machine learning methods for RNA secondary structure prediction | Zhao et al. | 2021 | 10.1371/journal.pcbi.1009291 | 107 | 26.75 | ✓ | Review |

Continued on next page

| Title | Authors | Year | DOI | Citations | Citations/Year | Relevance | Method |
|---|---|---|---|---|---|---|---|
| Deep learning models for RNA secondary structure prediction (probably) do not generalize across families | Szikszai et al. | 2022 | 10.1093/bioinformatics/btac415 | 80 | 26.67 | ✓ | Review |
| E2Efold-3D: end-to-end deep learning method for accurate de novo RNA 3D structure prediction | Shen et al. | 2022 | 10.48550/arXiv.2207.01586 | 80 | 26.67 | ✗ | |
| Improved RNA secondary structure and tertiary base-pairing prediction using evolutionary profile, mutational coupling and two-dimensional transfer learning | Singh et al. | 2021 | 10.1093/bioinformatics/btab165 | 100 | 25.00 | ✓ | SPOT-RNA 2 |
| Caveats to deep learning approaches to RNA secondary structure prediction | Flamm et al. | 2022 | 10.3389/fbinf.2022.835422 | 33 | 11.00 | ✓ | Review |

Continued on next page

| Title | Authors | Year | DOI | Citations | Citations/Year | Relevance | Method |
|---|---|---|---|---|---|---|---|
| Deep Learning Method for RNA Secondary Structure Prediction with Pseudoknots Based on Large-Scale Data | Shen et al. | 2021 | 10.1155/2021/6699996 | 8 | 2.00 | ✓ | Unamed |
| A survey on deep learning in DNA/RNA motif mining | He et al. | 2021 | 10.1093/bib/bbaa229 | 0 | 0.00 | ✗ | |
| trRosettaRNA: automated prediction of RNA 3D structure with transformer network | Wang et al. | 2023 | 10.1038/s41467-023-42528-4 | 0 | 0.00 | ✗ | |

**Supplemental Table S19**. **Google Scholar method identification.** Top 20 papers on Google Scholar sorted by relevance count for the search term “deep learning RNA secondary structure” released after 2020 (retrieved 2025-08-27).
