## Supplemental Figures for "Deep Learning for RNA Secondary Structure Determination: Gauging Generalizability and Broadening the Scope of Traditional Methods"

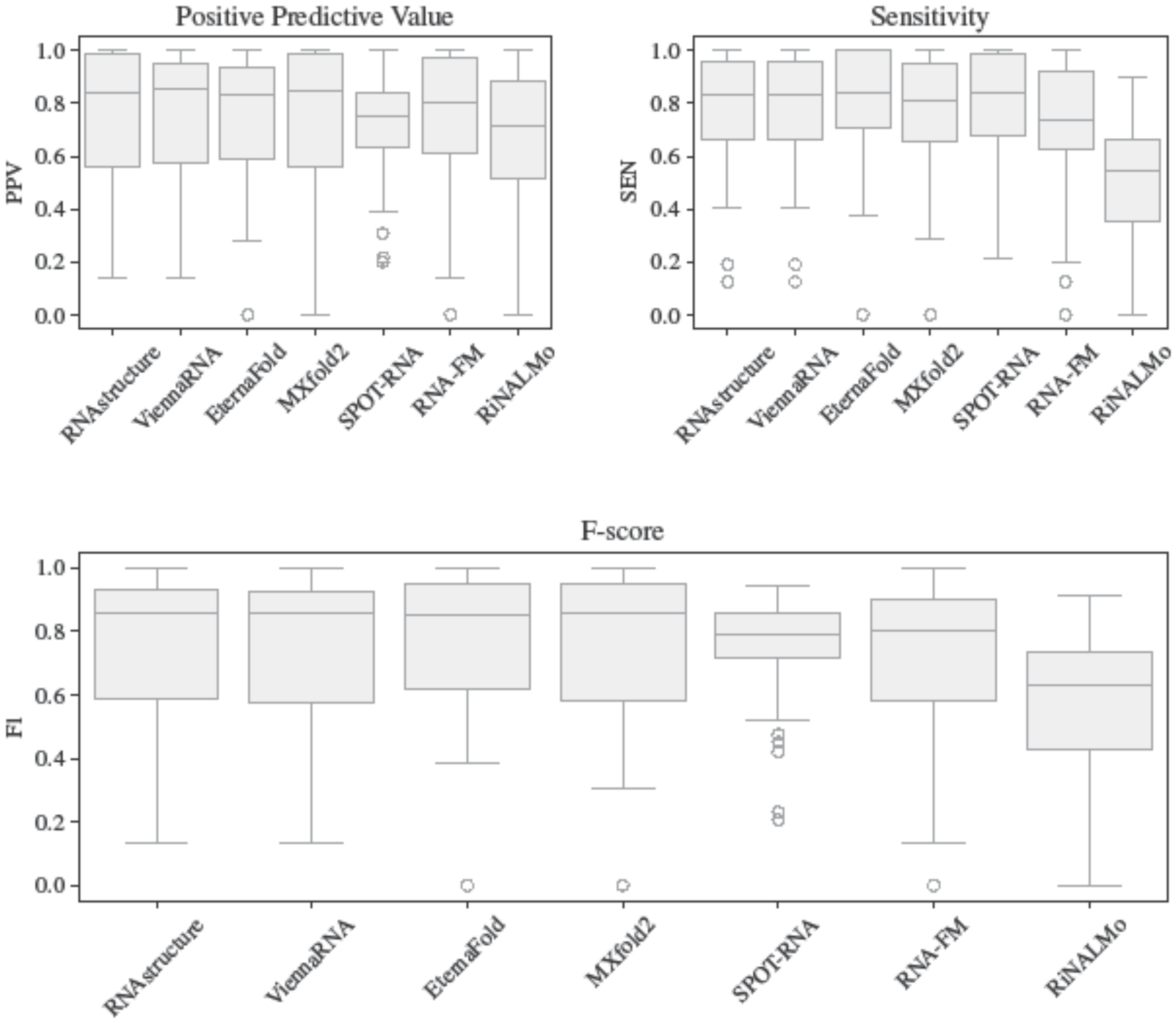


**Supplemental Figure S1. Boxplot of secondary structure prediction benchmarking results.** Positive Predictive Value (PPV), Sensitivity (SEN), and F-score (F1) of benchmarked data-driven methods compared to RNAstructure and ViennaRNA. Each group (e.g. Theophyline aptamer, minE/minF aptamer, Cas9 guide, etc.) is used as a single data point by averaging the targets within that group. All targets are included.

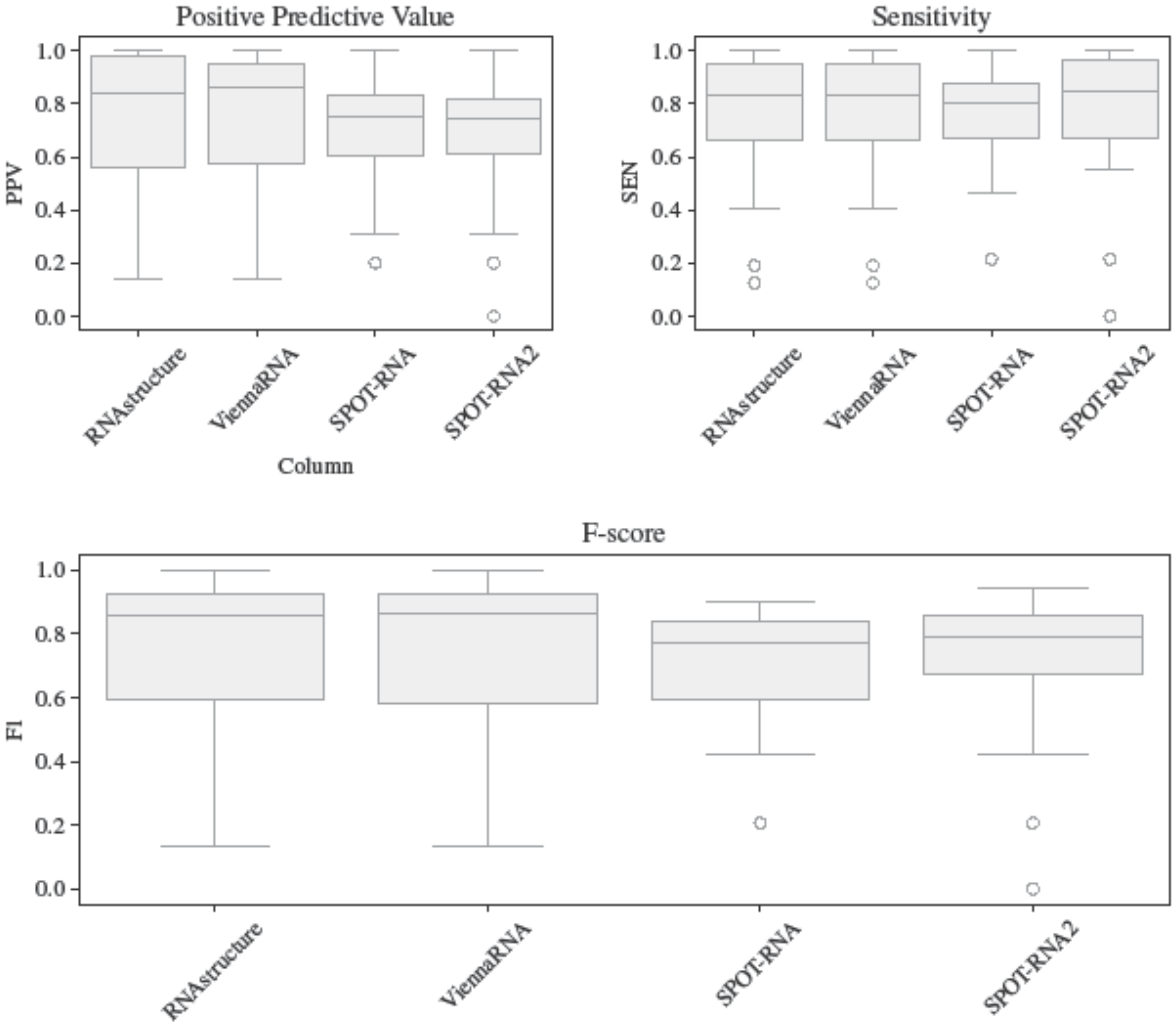


**Supplemental Figure S2. Boxplot of secondary structure prediction results for SPOT-RNA and SPOT-RNA 2.** Positive Predictive Value (PPV), Sensitivity (SEN), and F-score (F1) of benchmarked data-driven methods compared to RNAstructure and ViennaRNA. Each group (e.g. Theophyline aptamer, minE/minF aptamer, Cas9 guide, etc.) is used as a single data point by averaging the targets within that group. All overlaps found in Supplemental Table S8 (*, **) and Supplemental Table S9 (†, ‡) are excluded from this calculation.

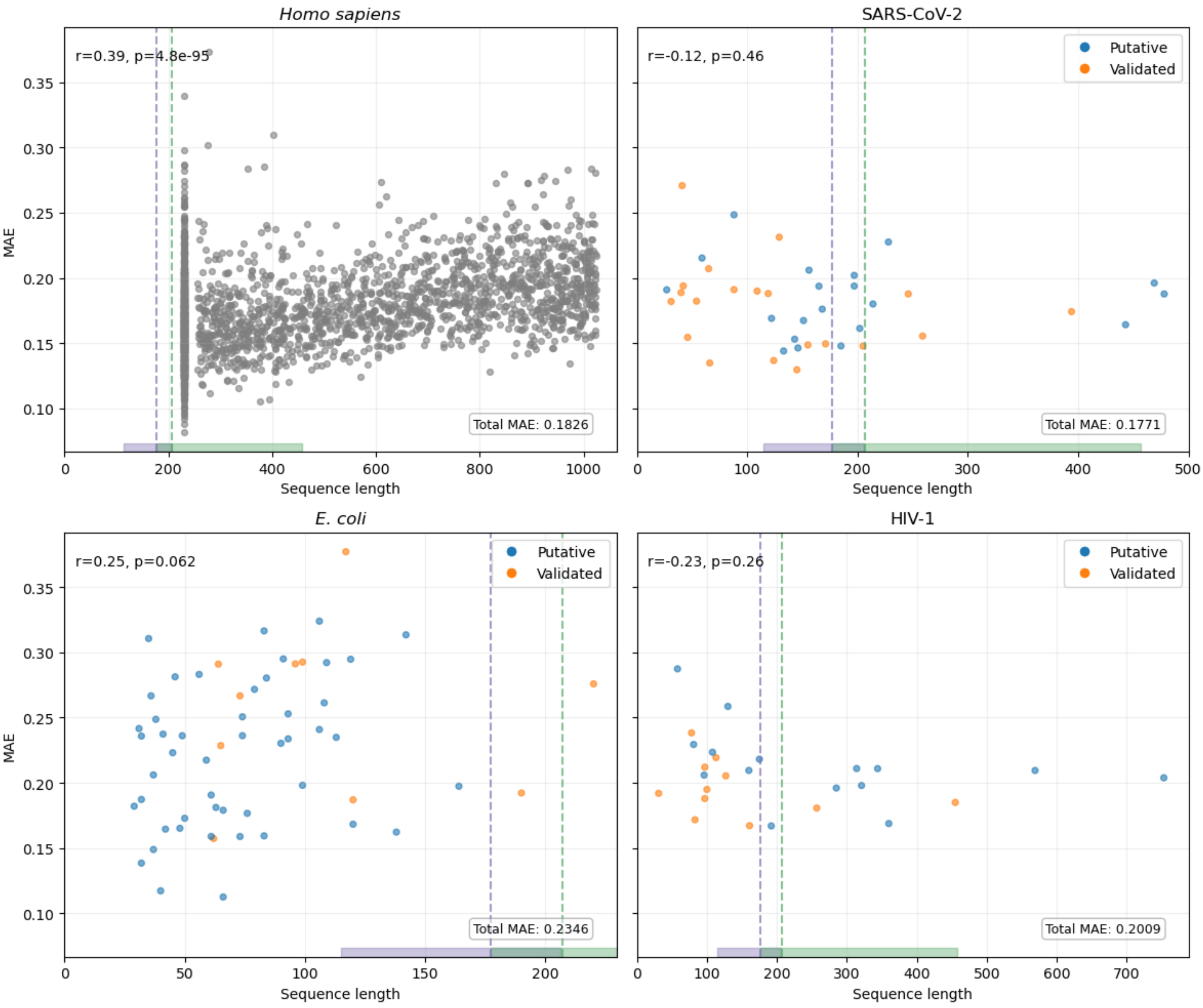


**Supplemental Figure S3. Relationship between sequence length and prediction error across four datasets.** Each point represents one sequence; orange indicates validated and blue indicates putative elements. The purple and green vertical lines mark the most prevalent sequence lengths in Ribonanza's train and test datasets (177nt and 207nt, respectively). The purple and green horizontal bars denote the range of sequence lengths in the train and test datasets (115–206nt and 177–457nt, respectively). Shown r and p values are the Pearson correlation coefficient and its associated *p*-value.

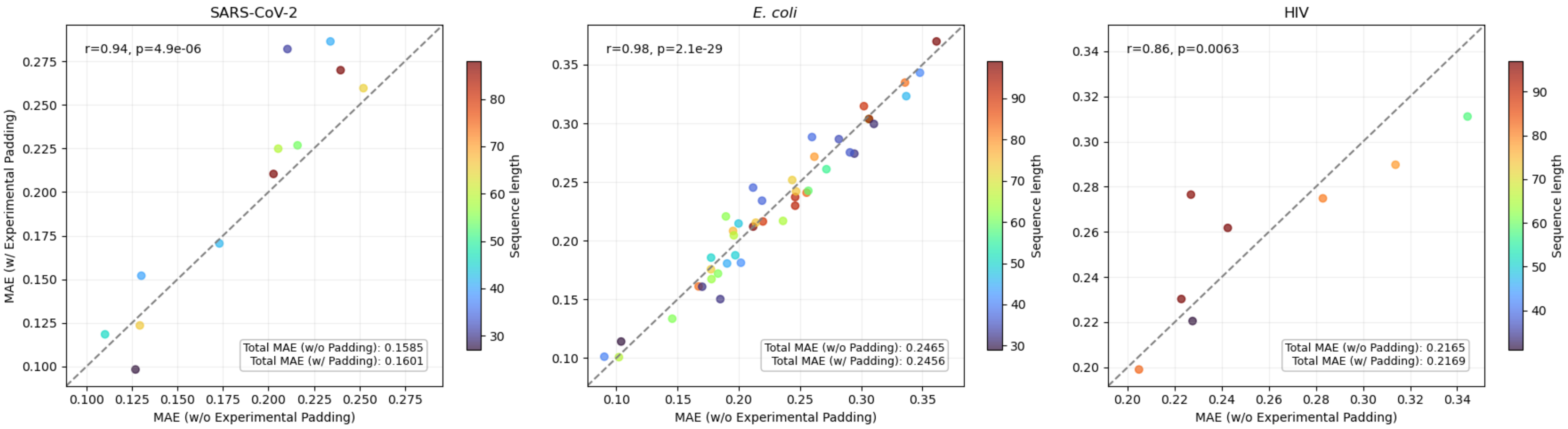


**Supplemental Figure S4. Effect of experimental padding on prediction error (MAE) for sequences with original length <100 nt.** Each point represents one sequence, color-coded by its length. The x and y axes show the MAE calculated without and with experimental padding, respectively. In both conditions, flanking sequences and barcodes were included. The dashed diagonal line indicates equality (i.e., y=x). Shown r and p values are the Pearson correlation coefficient and its associated p-value. For each panel, the inset reports the dataset-level MAE. For each panel, the inset reports dataset-level MAE.

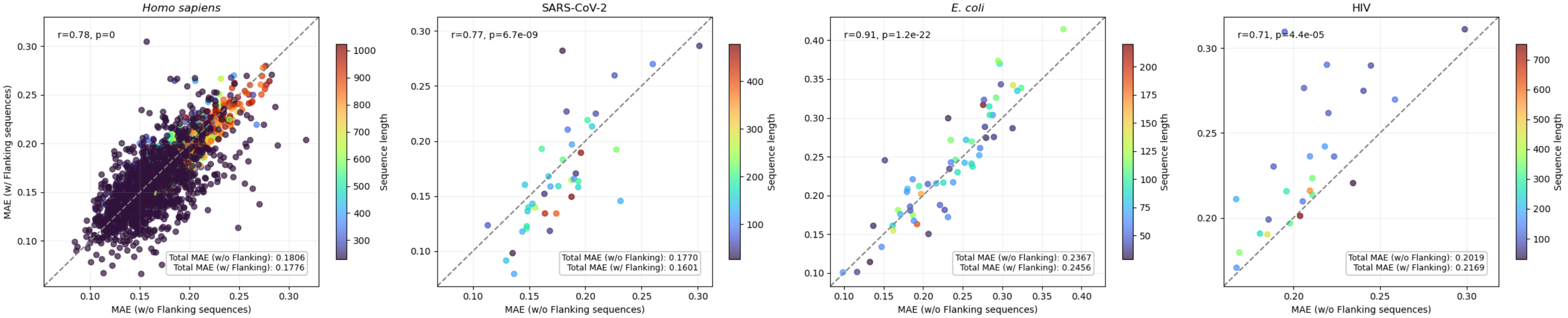


**Supplemental Figure S5. Effect of flanking sequences on prediction error (MAE) across four datasets.** Each point represents one sequence, color-coded by its length. The x and y axes show the MAE calculated without and with flanking sequences, respectively. Sequences shorter than 100nt were experimentally padded to 100nt. The dashed diagonal line indicates equality (i.e., y=x). Shown r and p values are the Pearson correlation coefficient and its associated p-value. For each panel, the inset reports the dataset-level MAE.

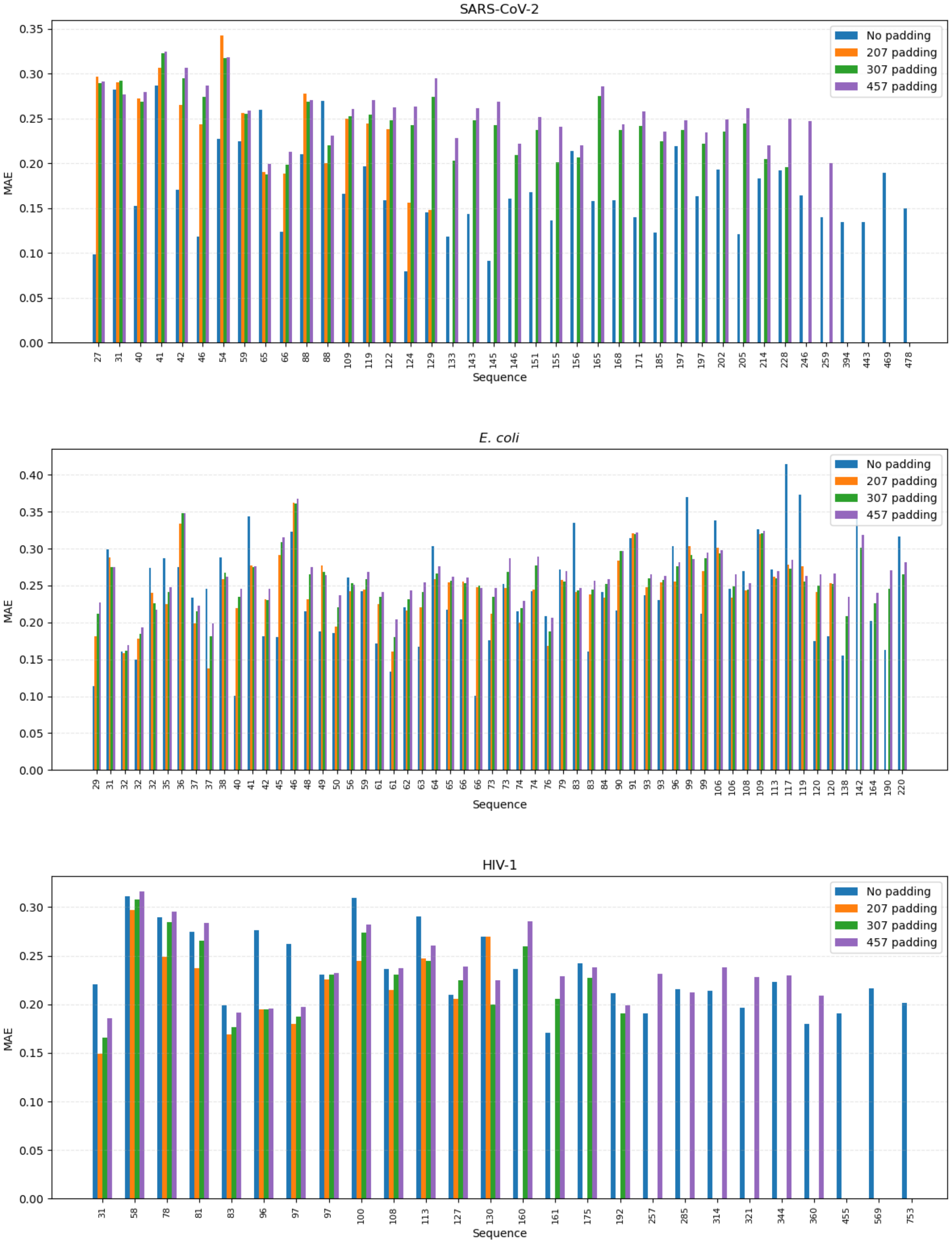


**Supplemental Figure S6. Effect of computational padding on prediction error.** All sequences include an added barcode and constant 3’ and 5’ flanking sequences. Sequences shorter than 100nt were experimentally padded to 100nt. Each group of bars represents one sequence, labeled by its original sequence length (without experimental padding, barcode, or flanking sequences). Bars indicate MAE values without padding (blue) or with computational padding up to 207nt (orange), 307nt (green), and 457znt (purple). Sequences longer than the target padding length were excluded from the corresponding condition.

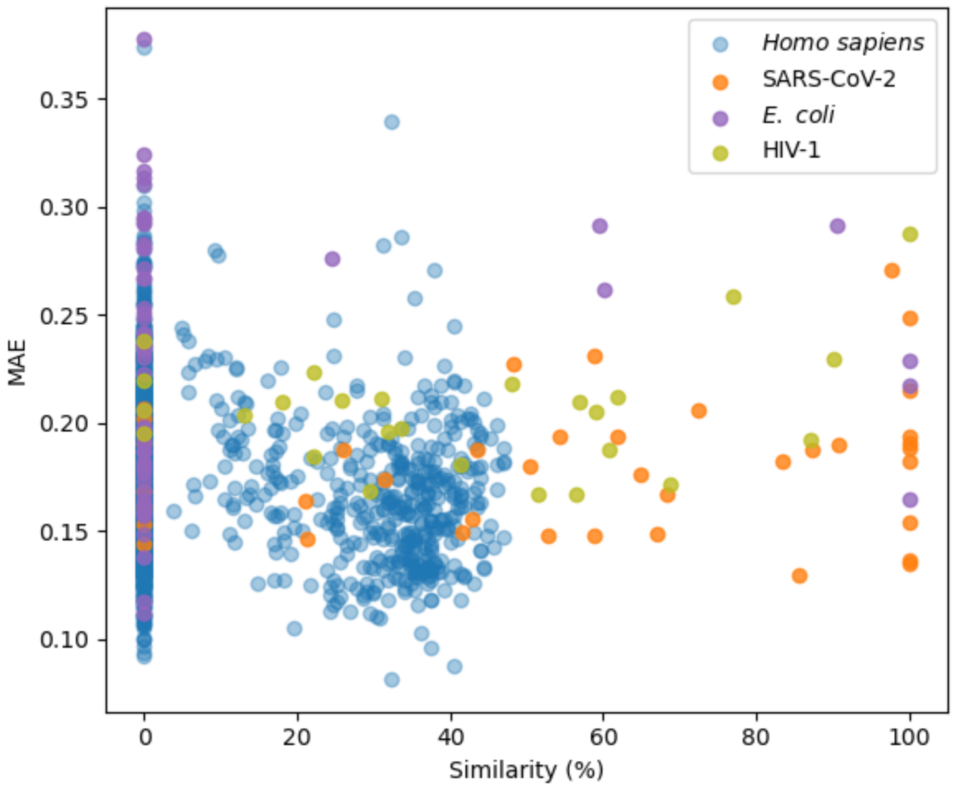


**Supplemental Figure S7**. **Relationship between sequence similarity and prediction error.** Each point represents one sequence, color-coded by dataset.

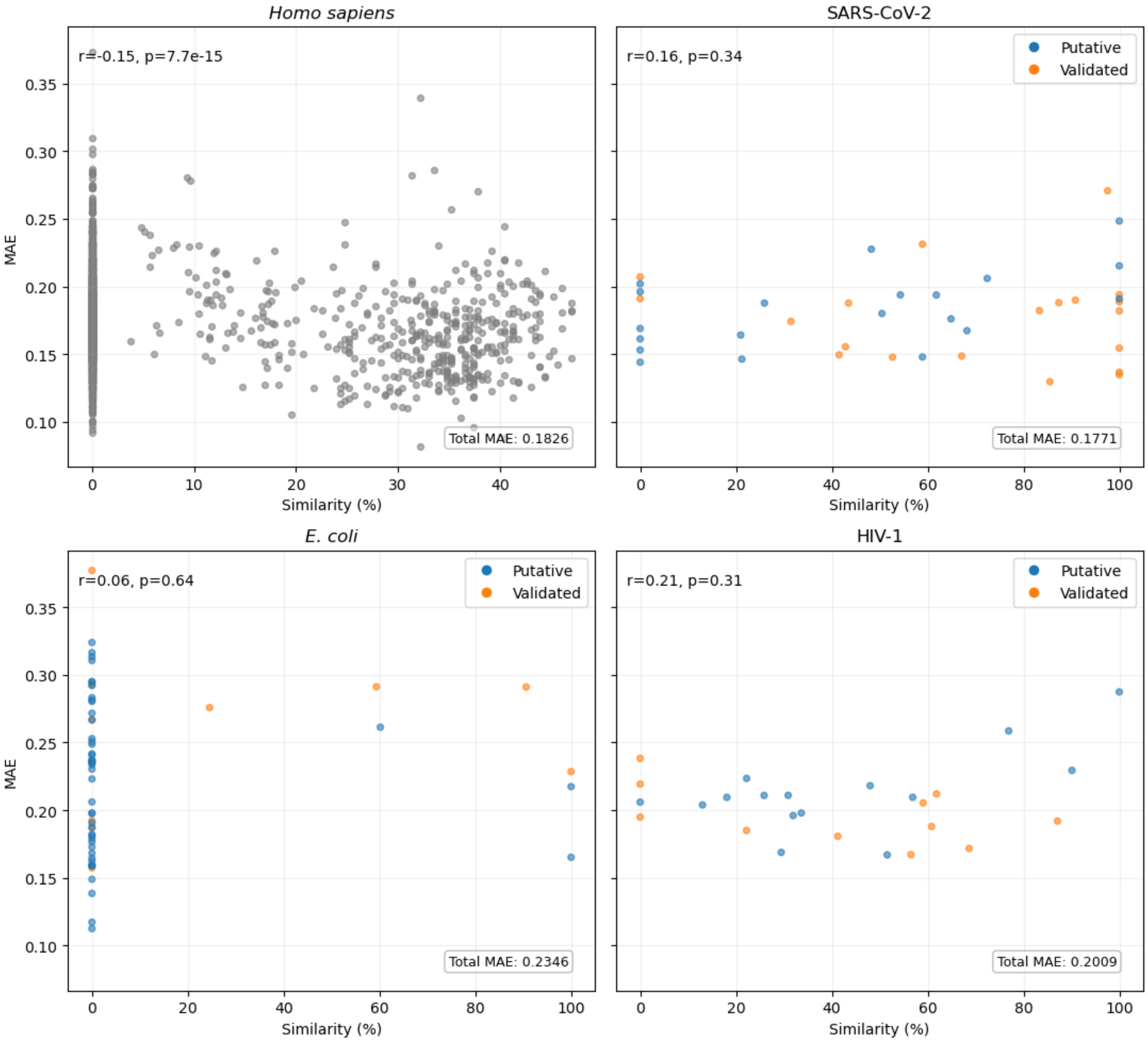


**Supplemental Figure S8. Relationship between sequence similarity and prediction error.** Each point represents one sequence, with blue indicating putative and orange indicating validated elements. Shown r and p values are Pearson correlation coefficients and their associated *p*-values. For each panel, the inset reports dataset-level MAE.

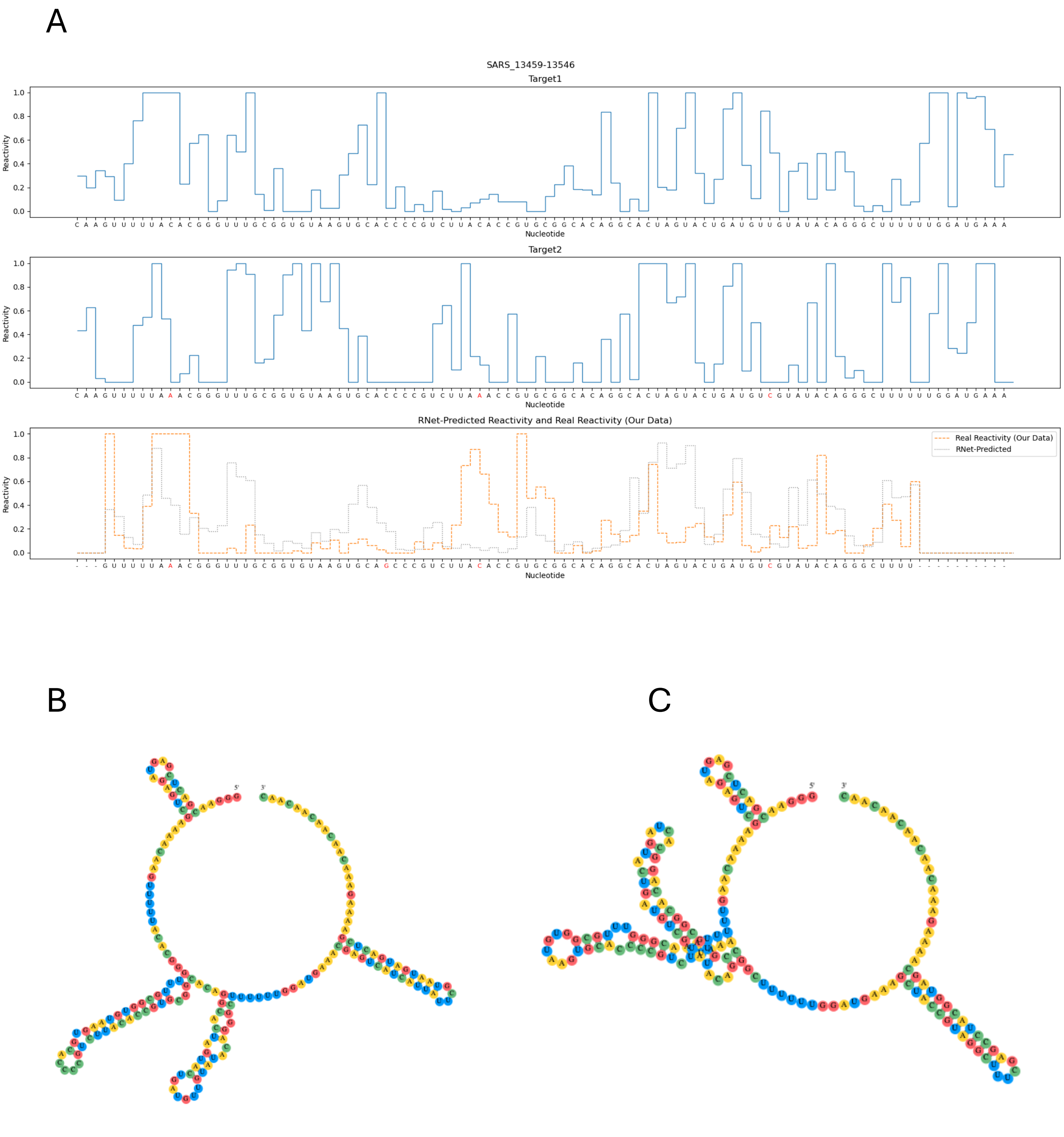


**Supplemental Figure S9. Comparison of reactivity profiles between two training instances corresponding to the same query, and their secondary structures predicted by EternaFold, in the SARS-CoV-2 dataset.**

**(A)** The top and middle panels show experimental reactivity profiles for two targets that are highly similar to the query. The bottom panel shows the experimental reactivity of the query (purple) and the RNet-predicted reactivity (green). Nucleotides aligned between the query and each target are shown on the x-axis. Nucleotides differing between target 1 (1e7d3d32d513) and target 2 (c90cbe2eff36) are highlighted in red on the target 2 x-axis, while nucleotides differing between the query and either target are highlighted in red on the query x-axis.

**(B, C)** Secondary structures of target 1 and target 2, respectively, predicted by EternaFold, and color-coded by nucleotide identity.

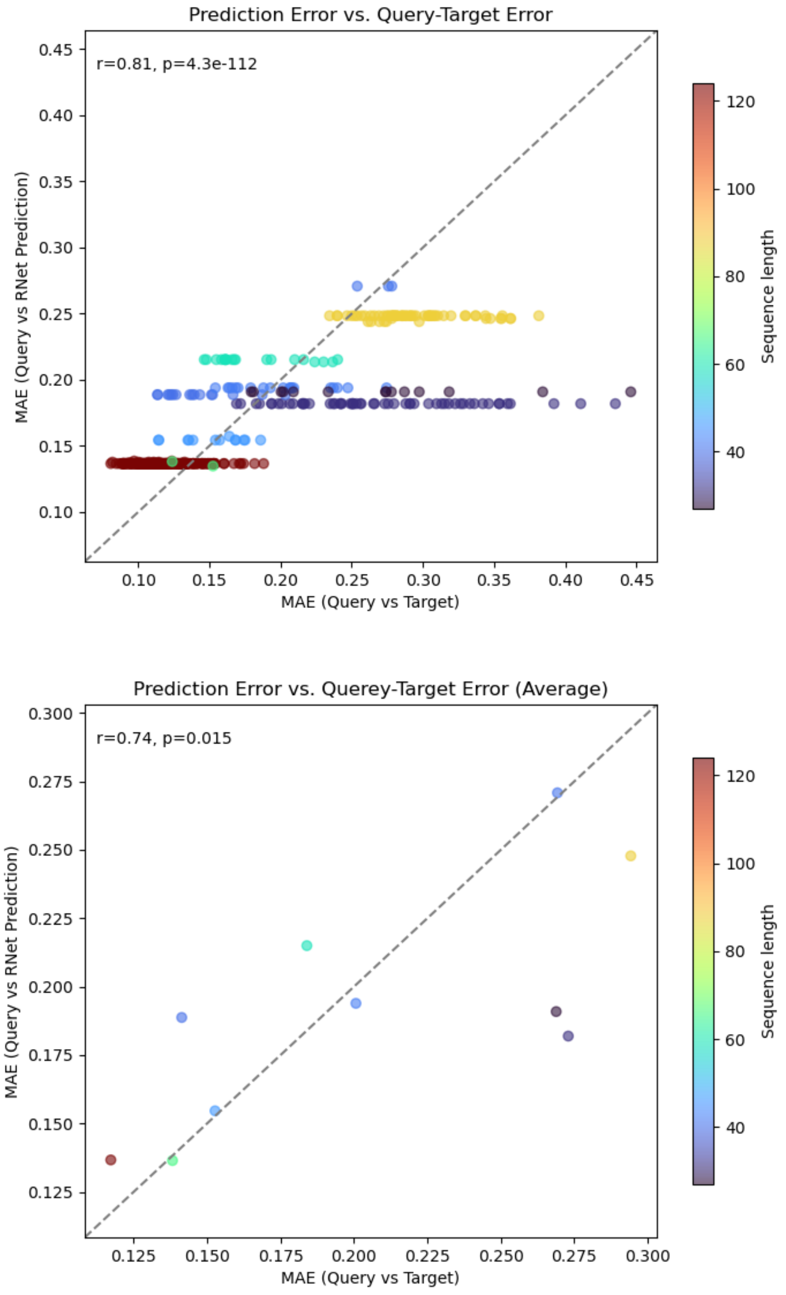


**Supplemental Figure S10. Relationship between query-target error and Rnet prediction error for our data (queries) against multiple training instances (targets) with high sequence similarity to queries in the SARS-CoV-2 dataset.** The x-axis shows the MAE between a query profile (our data) and a target profile (Ribonanza data for a highly similar training sequence). The y-axis shows the MAE between a query profile (our data) and RNet's prediction. Prediction errors were adjusted according to the nucleotides aligned between each query and its targets. The dashed diagonal line indicates equality (i.e., y=x). Shown r and p values are the Pearson correlation coefficient and its associated p-value. (A) Individual query-target pairs, illustrating substantial variation in similarity between our data (query profile) and the profiles of Ribonanza targets with high similarity to the query sequence. Each point represents a query-target pair, color-coded by query sequence length. (B). MAE values averaged over all targets matching to each query compared with the MAE for RNet's predictions.

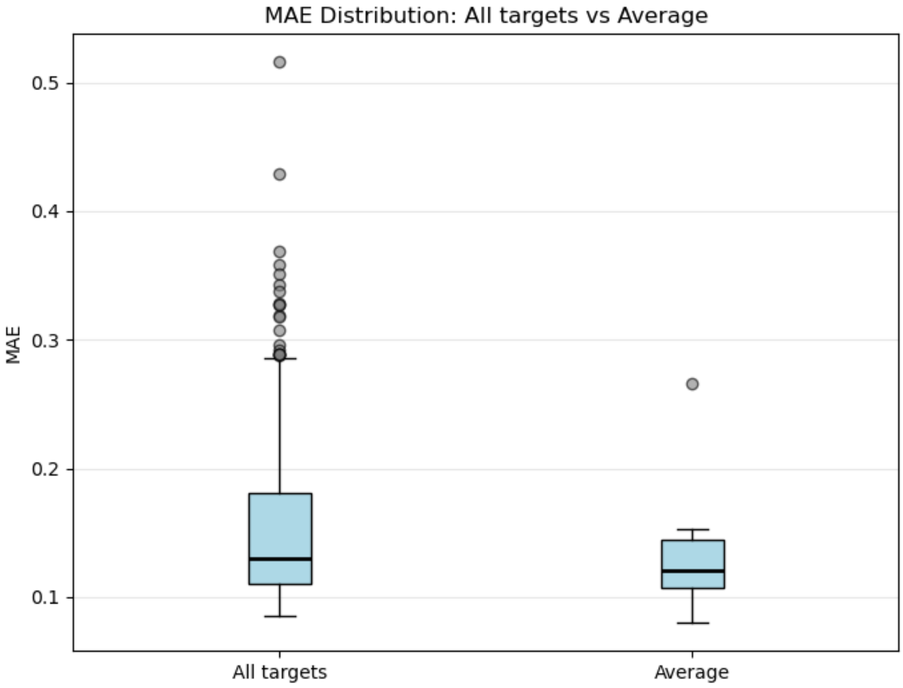


**Supplemental Figure S11**. **Distribution of MAE between Ribonanza data reactivity and RNet-predicted reactivity for individual targets (All targets) versus averaged profiles (Average) in the SARS-CoV-2 dataset**.

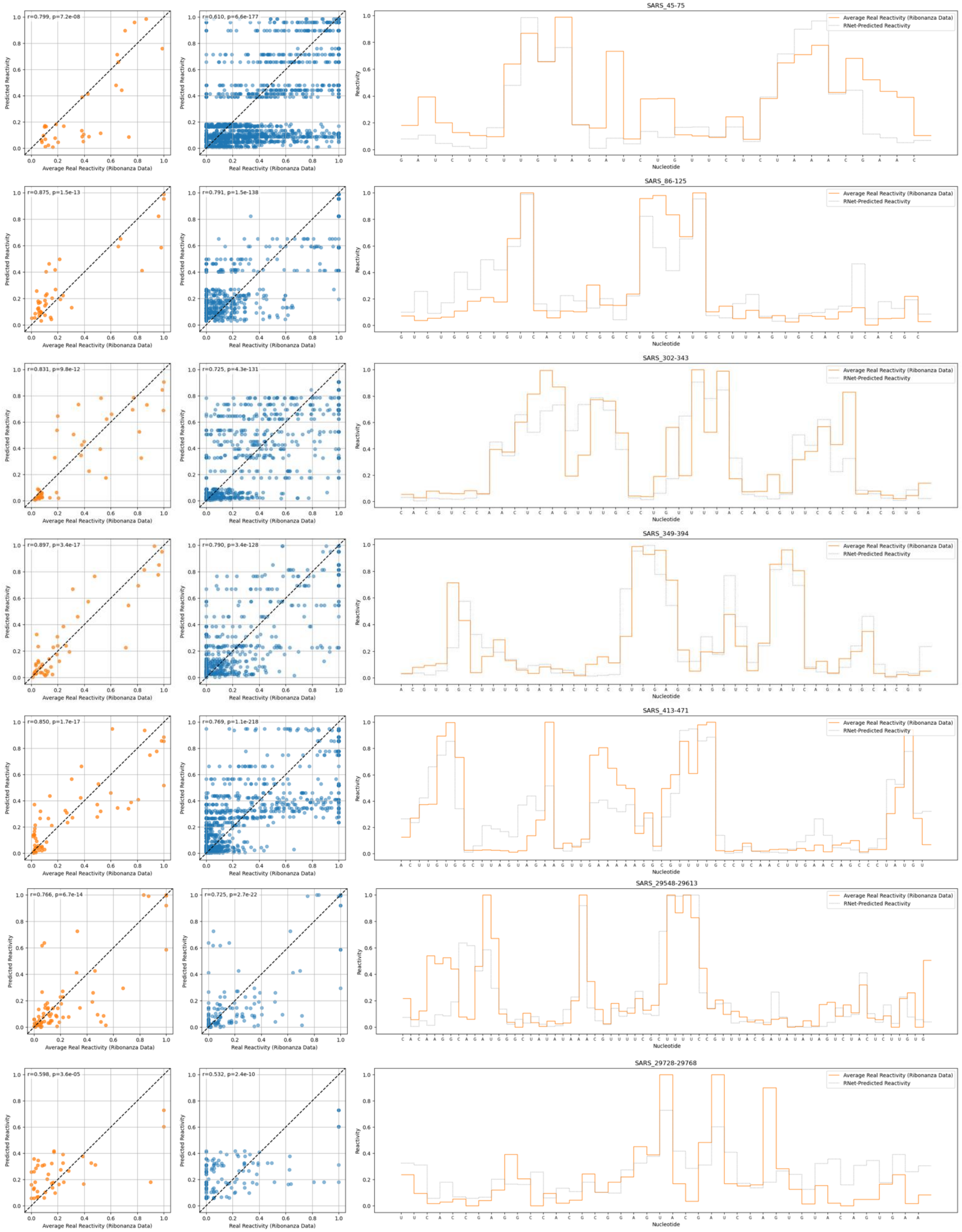

**Supplemental Figure S12. Comparison of RNet predictions with averaged and individual training profiles for selected queries in the SARS-CoV-2 dataset.** Each row corresponds to a query sequence that had multiple targets of high sequence similarity in the Ribonanza data. Left: Predicted reactivity plotted against the averaged target profile. Middle: Predicted reactivity plotted against multiple individual target profiles, illustrating high variability among targets. Right: Reactivities predicted by RNet (gray) and reactivities derived from Ribonanza data by averaging over all targets (orange). Also shown are the Pearson correlation coefficient (r) and its associated *p*-value (p), the query sequence, and each element's transcript coordinates.

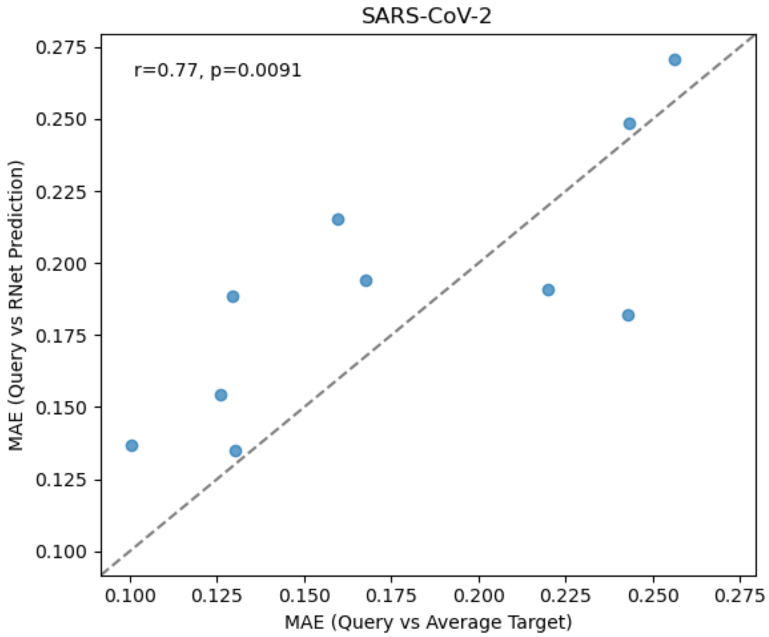


**Supplemental Figure S13. Comparison between RNet prediction error and the error of averaged profiles of highly similar targets in the SARS-CoV-2 dataset.** The x-axis shows the MAE between each query's reactivity profile and the averaged reactivity profile of training sequences with high sequence similarity to that query (Query vs. Average Target). The y-axis shows the MAE between each query's reactivity profile and Rnet's prediction (Query vs. RNet Prediction). Points below and above the dashed diagonal line correspond to sequences for which RNet performs better and worse than the average target profile, respectively. Shown r and p are the Pearson correlation coefficient and its *p*-value.

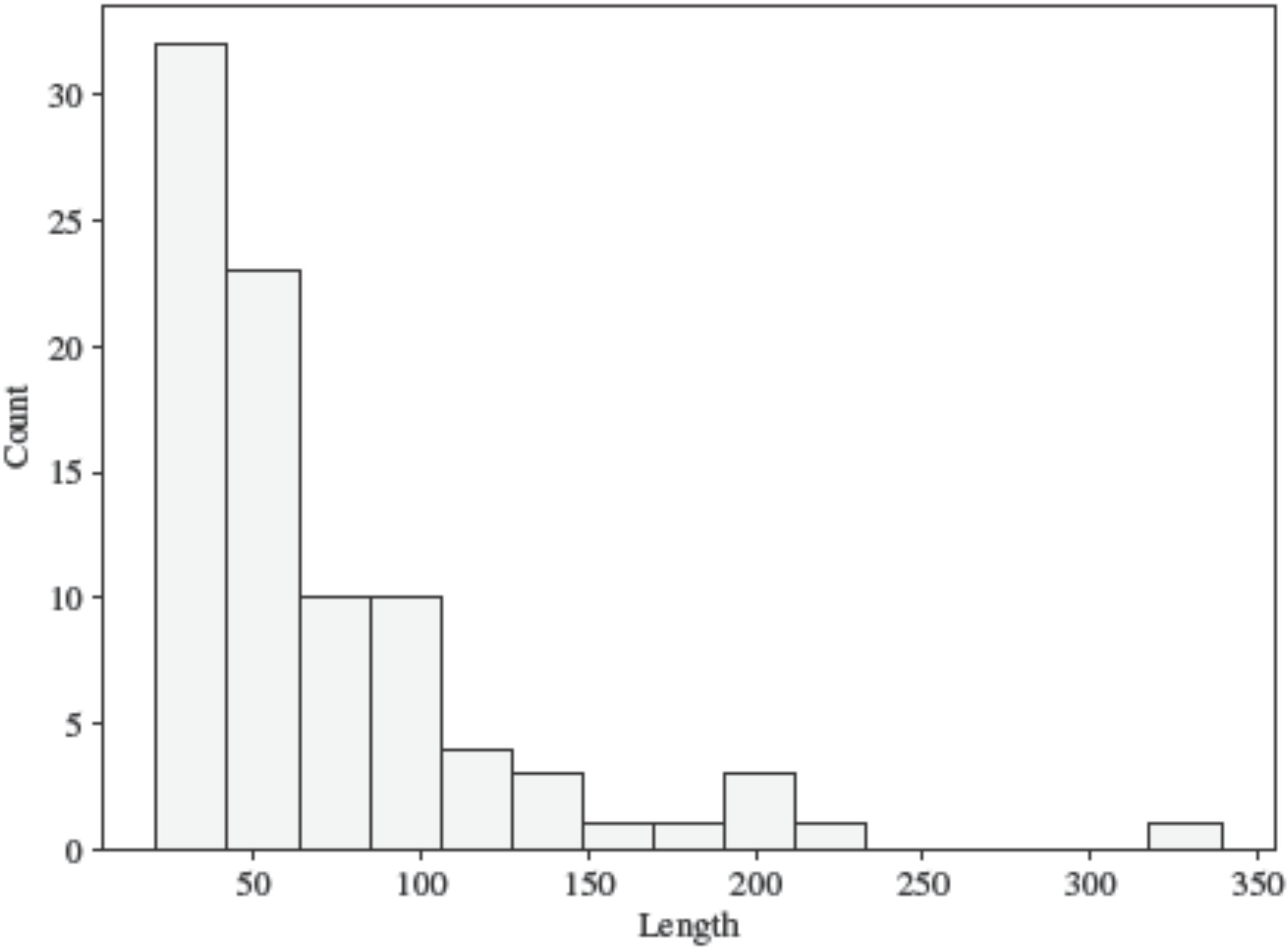


**Supplemental Figure S14**. **Distribution of sequence lengths in our benchmarking dataset.** Note that these are the lengths of the sequences after pre-processing, and not the chain lengths.

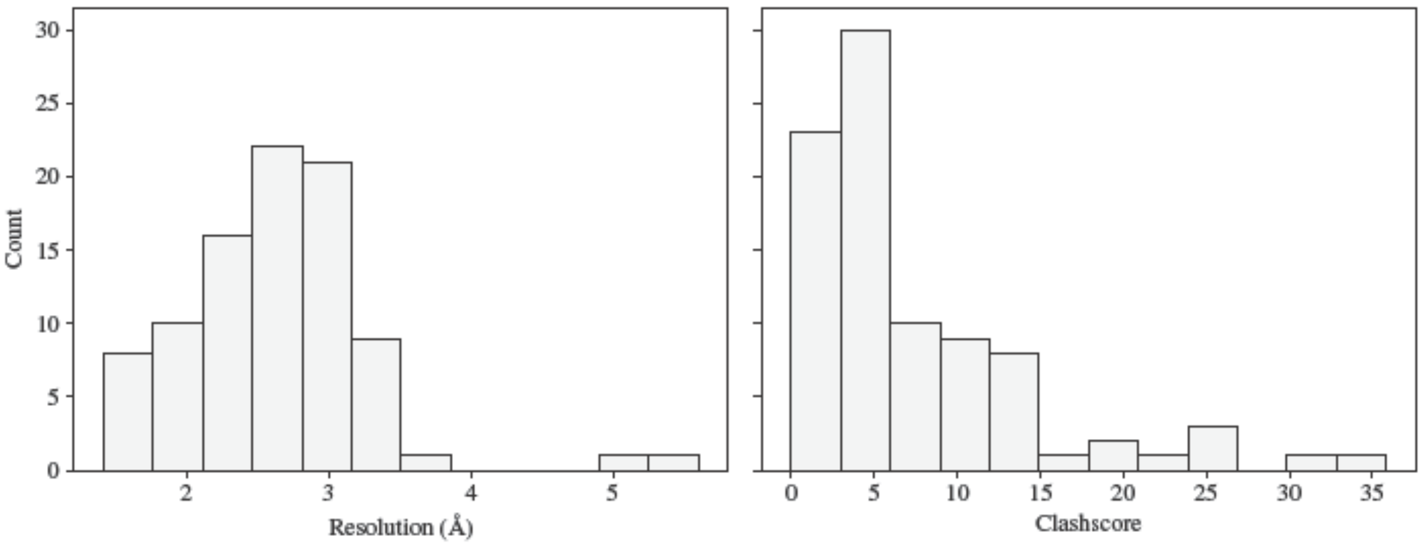


**Supplemental Figure S15**. **Distribution of resolution and clashscore in our benchmarking dataset.**
