## Supplemental Methods for "Deep Learning for RNA Secondary Structure Determination: Gauging Generalizability and Broadening the Scope of Traditional Methods"

Marcell Szikszai[1,2], Ting-Yuan Wang[3], Ryan Krueger[4], David H. Mathews[5,6], Max Ward[1], Sharon Aviran[3,7]

**Author Affiliations**

1 Department of Computer Science and Software Engineering, The University of Western Australia, Crawley, WA 6009, Australia

2 Department of Molecular and Cellular Biology, Harvard University, Cambridge, MA 02138, USA

3 Department of Biomedical Engineering, University of California Davis, Davis, CA 95616, USA

4 School of Engineering and Applied Sciences, Harvard University, Cambridge, MA 02138, USA

5 Department of Biochemistry and Biophysics, University of Rochester Medical Center, Rochester, NY 14642, USA

6 Center for RNA Biology, University of Rochester Medical Center, Rochester, NY 14642, USA

7 Genome Center, University of California Davis, Davis, CA 95616, USA

### Data Collection

Details of the datasets analyzed in this study are summarized below. In total, four datasets were utilized, encompassing bacterial, viral, and human RNAs. Structural motifs shorter than 25 nucleotides were excluded from analysis. Below, we describe each dataset in detail.

*E. coli*

All structured regions and reactivity data were collected from prior work (Mustoe et al., 2018). Mustoe et al. identified 58 structural motifs using low SHAPE/low-entropy methods, 9 of which have been independently verified in earlier studies (Chiaruttini et al., 1996; Desnoyers et al., 2009; Fu et al., 2013). They split *rpmI* 5′ UTR into two well-defined sub-motifs for ease of analysis, yielding a total of 10 motifs that comprise our validated dataset. For the remaining 49 motifs, we excluded the 5′ UTR rpoS region (-) 2865600-2865661 due to its high proportion of missing values. The final putative dataset, therefore, consists of 48 motifs. Reactivity profiles were obtained from RASP v2.0 (Mu et al., 2025).

SARS-CoV-2

We used reactivity profiles from a prior study (Manfredonia et al., 2020), which we downloaded from the Incarnato laboratory website (https://www.incarnatolab.com/datasets/SARS_Manfredonia_2020.php). The conserved motifs reported in this study were included and designated as validated. In addition, we incorporated 5′ UTR stem–loops 1–7 as validated elements, as they are consistently reported across multiple studies, while 5′ UTR stem–loops 1–8 was included as putative since stem-loop 8 was not supported by their data. Additional structures were collected from other works. Hagey et al. identified two conserved structures, both of which were included and classified as validated (Hagey et al., 2022). Lan et al. investigated the frameshifting element (FSE), which we incorporated and designated as putative (Lan et al., 2022). Furthermore, Wacker et al. applied NMR spectroscopy and verified 15 structures (Wacker et al., 2020). Of these, one was excluded because it contained only a single stem within a stem–loop, another was excluded due to a high proportion of missing values, and three were excluded due to redundancy, as two overlapped with stem–loop 5 in the 5′ UTR identified by Manfredonia et al. and one overlapped with the FSE reported by Lan et al. The remaining Wacker et al. structures were included and marked as validated, except for two that were noted in their study as requiring further investigation. Yang et al. applied evolutionary covariance analysis and reported 20 structural elements (Yang et al., 2025). Four of these overlapped with motifs identified by Manfredonia et al. or with the FSE described by Lan et al. and were therefore excluded. The remaining structures were retained and designated as putative. Because different viral strains were analyzed across studies, we performed manual sequence alignment to reconcile coordinate differences, particularly for motifs reported by Hagey et al. and Yang et al.

HIV-1

Structural motifs and reactivity profiles were obtained directly from (Siegfried et al., 2014). In that study, SHAPE-MaP was applied to identify structured elements based on low SHAPE reactivity and low Shannon entropy. We collected the *de novo* structures shown in Figure 4, which highlights regions with low SHAPE reactivity and low Shannon entropy. As these motifs were newly proposed and not well-validated, we designated them as putative. In addition, their Supp. Table 2 lists previously verified functional structures. From this list, we excluded splice donor and acceptor sites as well as all pseudoknots. The remaining motifs were incorporated into our analysis and designated as validated. Reactivity profiles were downloaded from RASP v2.0 (Mu et al., 2025).

*H. sapiens*

The RNAndria dataset comprises 1,456 human mRNA regions and 1,098 pri-miRNA sequences, each probed *in vitro* using DMS-MaPseq (Lajarte et al., 2024). It provides high-resolution structural information for long and diverse RNAs, extending our bacterial and viral datasets. There was no need to delineate independently folding regions or label them as validated or putative since these regions were folded *in vitro* in their entirety and then probed by DMS. The data were downloaded from https://huggingface.co/rouskinlab/datasets.

**Experimental padding**

Some of the sequences in the Ribonanza data are padded prior to probing, such that most probed sequences are of a fixed length. The overwhelming majority of training sequences are 177 nt long. By subtracting 30, which is the barcode length in most training sequences, and further subtracting 47, which is the total length of the constant 5' and 3' flanking regions, we are left with 100 nt as the region of interest. Based on this and on He et al.'s description of their work, we concluded that they predominantly padded training sequences shorter than 100 (He et al., 2024). We therefore padded only sequences shorter than 100 using He et al.'s code (downloaded from https://github.com/DasLab/big_library_design on 10/13/25), which aims to ensure that the added nucleotides do not interfere with the region of interest during probing. A detailed description of padding and barcoding procedures and other design considerations is provided in this site.

**Computational padding**

When training deep learning models, it is common practice to use batch processing to improve efficiency. However, RNA sequences vary in length, requiring padding to achieve uniform input dimensions. During inference, sequences are sorted by length before batching, so sequences of similar size are grouped together, thereby minimizing padding. Nevertheless, padding remains necessary in some cases. In RibonanzaNet, sequences are padded with the token "N" (in addition to A, U, C, and G). In our experiments, sequences were padded to lengths of 177, 207, 307, and 457 nucleotides. The lengths 177 and 207 nt are the most prevalent in the Ribonanza dataset, while 307 and 457 nt were chosen to evaluate the effect of longer padding, as these lengths are also present in the testing set. Padding was applied at inference by modifying the parameter max_len in dataset.py of the RibonanzaNet repository, set to max([max(length), length_to_pad]).

**Barcodes and constant regions**

In the Ribonanza data, barcodes are designed as stem–loop structures containing a conserved UUCG tetraloop and variable stem lengths, with the most common stem length being 13 base-pairs, resulting in a total barcode length of 30 nucleotides. For our analyses, we used barcodes of 30 nucleotides taken from Ribonanza test sequences. All sequences were also flanked by constant sequences: a 5′ sequence (5′-GGGAACGACUCGAGUAGAGUCGAAAA-3′) and a 3′ sequence (5′-AAAAGAAACAACAACAACAAC-3′). For the comparison experiments, sequences were processed in the same manner, with both barcodes and constant regions added to our dataset.
