## Supplemental File 1 for "Deep Learning for RNA Secondary Structure Determination: Gauging Generalizability and Broadening the Scope of Traditional Methods"

### Supplemental File 1: Deep Learning for RNA Secondary Structure Determination: Gauging Generalizability and Broadening the Scope of Traditional Methods

Marcell Szikszai[1, 2], Ting-Yuan Wang[3], Ryan Krueger[4], David H. Mathews[5, 6], Max Ward[1], and Sharon Aviran[3, 7]

[1]Department of Computer Science and Software Engineering, The University of Western Australia, Crawley, WA 6009, Australia
[2]Department of Molecular and Cellular Biology, Harvard University, Cambridge, MA 02138, USA
[3]Department of Biomedical Engineering, University of California Davis, Davis, CA 95616, USA
[4]School of Engineering and Applied Sciences, Harvard University, Cambridge, MA 02138, USA
[5]Department of Biochemistry and Biophysics, University of Rochester Medical Center, Rochester, NY 14642, USA
[6]Center for RNA Biology, University of Rochester Medical Center, Rochester, NY 14642, USA
[7]Genome Center, University of California Davis, Davis, CA 95616, USA

### 1 Aptamers

#### 1.1 Theophyline aptamer

### 1.1.1 8d28_A

Crystal structure of theophylline aptamer in complex with theophylline

**Release date:** 2022-11-30
**Method:** x-ray diffraction
**Resolution:** 1.42 Å
**Chain length:** 33
**Extracted length:** 33

**Total clashes:** 4
**Clashes per residue:** 0.121
**Clashscore:** 1.840

**Description:** RNA (33-MER)
**Organism:** synthetic construct

**Menichelli, E., Lam, B.J., Wang, Y., Wang, V.S., Shaffer, J., Tjhung, K.F., Bursulaya, B., Nguyen, T.N., Vo, T., Alper, P.B., McAllister, C.S., Jones, D.H., Spraggon, G., Michellys, P.Y., Joslin, J., Joyce, G.F., Rogers, J. (2022) Discovery of small molecules that target a tertiary-structured RNA. Proc.Natl.Acad.Sci.USA.**
**DOI:** 10.1073/pnas.2213117119

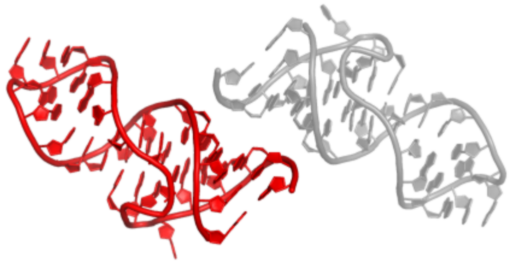

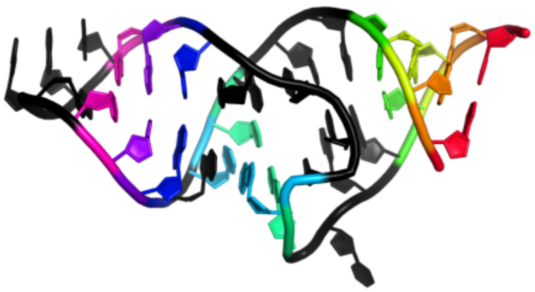

GGCGAUACCAGCCGAAAGGCCCUUGGCAGCGCC
((((...((.(((....)))....))...))))

### 1.1.2 8d29_C

Crystal structure of theophylline aptamer - apo form

**Release date:** 2022-11-30
**Method:** x-ray diffraction
**Resolution:** 1.81 Å
**Chain length:** 34
**Extracted length:** 34

**Total clashes:** 4
**Clashes per residue:** 0.118
**Clashscore:** 4.310

**Description:** RNA (34-MER)
**Organism:** synthetic construct

---

**Menichelli, E., Lam, B.J., Wang, Y., Wang, V.S., Shaffer, J., Tjhung, K.F., Bursulaya, B., Nguyen, T.N., Vo, T., Alper, P.B., McAllister, C.S., Jones, D.H., Spraggon, G., Michellys, P.Y., Joslin, J., Joyce, G.F., Rogers, J. (2022) Discovery of small molecules that target a tertiary-structured RNA. Proc.Natl.Acad.Sci.USA.**
**DOI:** 10.1073/pnas.2213117119

---

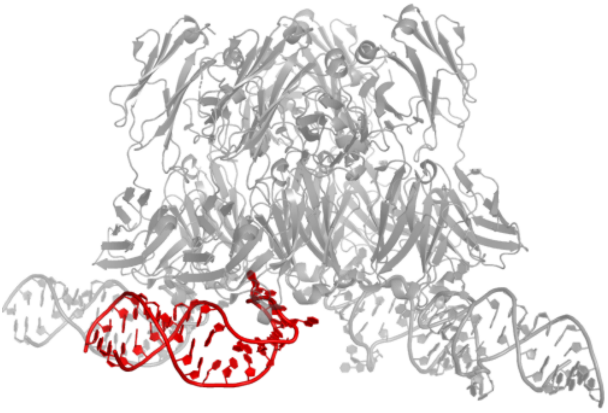

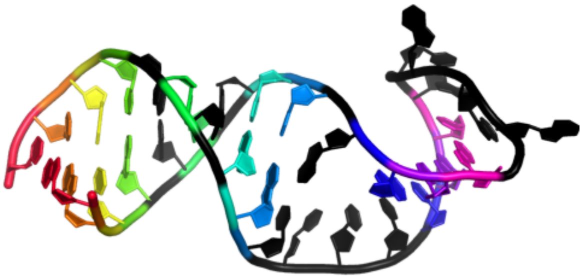

```
GGCGAUACCAGCGAAACACGCCCUUGGCAGCGUC
((((.(.((.(((.....)))....)).).))))
```

### 1.1.3 8dk7_C

Crystal structure of theophylline aptamer soaked with TAL2

**Release date:** 2022-11-30
**Method:** x-ray diffraction
**Resolution:** 2.46 Å
**Chain length:** 34
**Extracted length:** 34

**Total clashes:** 4
**Clashes per residue:** 0.118
**Clashscore:** 24.590

**Description:** RNA (34-MER)
**Organism:** synthetic construct

**Menichelli, E., Lam, B.J., Wang, Y., Wang, V.S., Shaffer, J., Tjhung, K.F., Bursulaya, B., Nguyen, T.N., Vo, T., Alper, P.B., McAllister, C.S., Jones, D.H., Spraggon, G., Michellys, P.Y., Joslin, J., Joyce, G.F., Rogers, J. (2022) Discovery of small molecules that target a tertiary-structured RNA. Proc.Natl.Acad.Sci.USA.**
**DOI:** 10.1073/pnas.2213117119

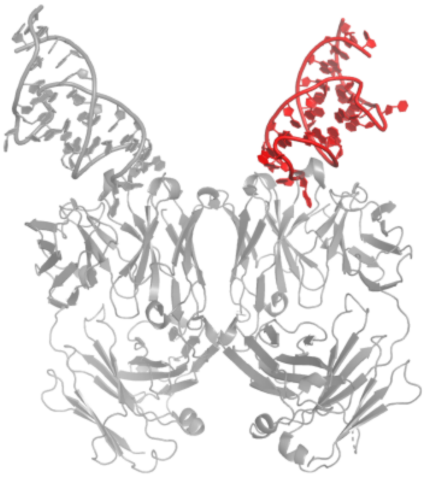

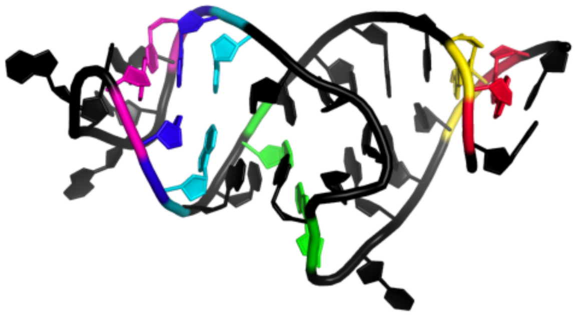

```
GGCGAUACCAGCGAAACACGCCCUUGGCAGCGUC
.((....(..(((.....))).....)....)).
```

#### 1.2 minE/minF aptamer

### 1.2.1 4m4o_B

Crystal structure of the aptamer minE-lysozyme complex

**Release date:** 2013-12-18
**Method:** x-ray diffraction
**Resolution:** 2.00 Å
**Chain length:** 59
**Extracted length:** 59

**Total clashes:** 14
**Clashes per residue:** 0.237
**Clashscore:** 2.830

**Description:** RNA (59-MER)
**Organism:** nan

**Malashkevich, V.N., Padlan, F.C., Toro, R., Girvin, M., Almo, S.C. (N/A) Crystal structure of the aptamer minE-lysozyme complex. to be published.**
**DOI:** nan

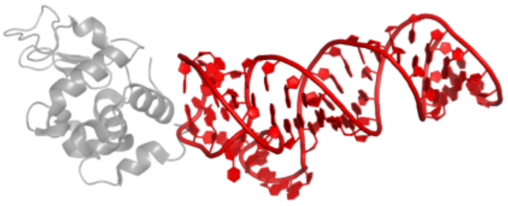

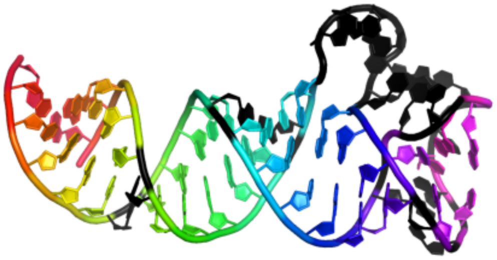

GGGUUCAUCAGGGCUAAAGAGUGCAGAGUUACUUAGUUCACUGCAGACUUGACGAACCC
((((((.(((((.((.......((((.(.(.<..)>..).)))))).))))).))))))

#### 1.2.2 4m6d_L

Crystal structure of the aptamer minF-lysozyme complex.

**Release date:** 2013-12-11
**Method:** x-ray diffraction
**Resolution:** 2.68 Å
**Chain length:** 45
**Extracted length:** 43

**Total clashes:** 62
**Clashes per residue:** 1.378
**Clashscore:** 18.470

**Description:** aptamer
**Organism:** nan

---

**Malashkevich, V.N., Padlan, F.C., Toro, R., Girvin, M., Almo, S.C. (N/A) Crystal structure of the aptamer minF-lysozyme complex. To be Published.**
**DOI:** nan

---

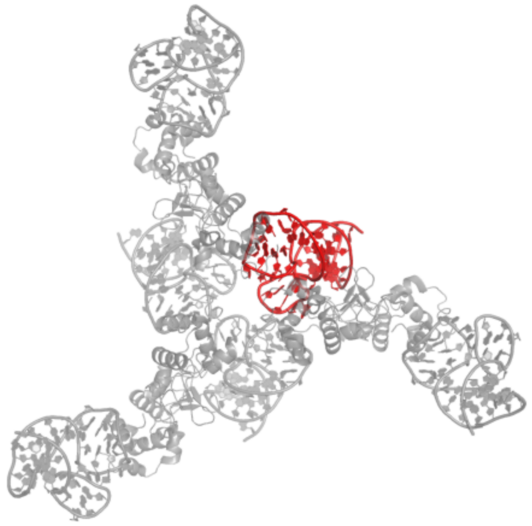

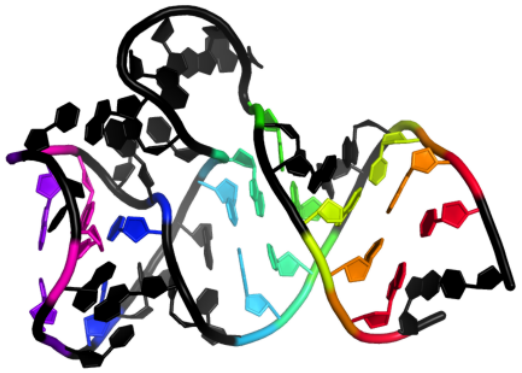

GGGCGGCUAAAGAGUGCAGAGUUACUUAGUUCACUGCAGACGCCC
..(((..(.......((...(.(.<..)>..)...)))..)))..

#### 1.3 RhoBAST aptamer

##### 1.3.1 9bun_A

RhoBAST aptamer RNA in complex with 5(6)-carboxytetramethylrhodamine

**Release date:** 2024-06-05
**Method:** x-ray diffraction
**Resolution:** 2.10 Å
**Chain length:** 48
**Extracted length:** 48

**Total clashes:** 9
**Clashes per residue:** 0.188
**Clashscore:** 11.130

**Description:** RNA (48-MER)
**Organism:** synthetic construct

**Batey, R.T., Siwik, S.H. (N/A) Structure of RhoBAST RNA aptamer in complex with 5(6)-carboxytetramethylrhodamine (TAMRA). To Be Published.**
**DOI:** nan

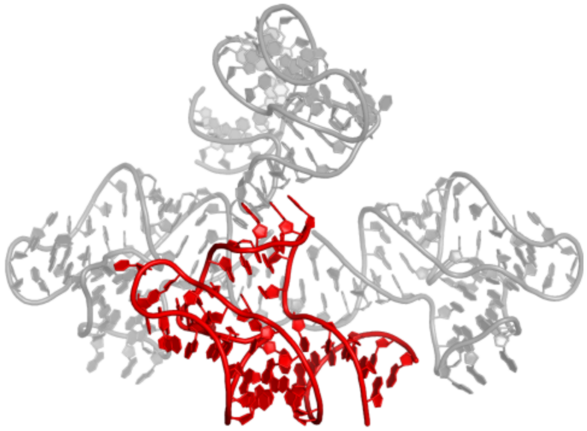
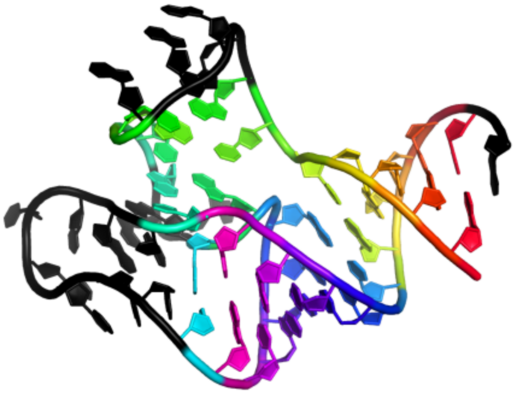

```
GGACUCGGAAACGUGAAGGAGAGGCGCAAGGUUAACCGCCUCAGUCCA
(((((((....))(<...<)((((((>........>))))))))))).
```

### 1.3.2 8jy0_D

Crystal structure of RhoBAST complexed with TMR-DN

**Release date:** 2024-05-29
**Method:** x-ray diffraction
**Resolution:** 2.75 Å
**Chain length:** 64
**Extracted length:** 64

**Total clashes:** 41
**Clashes per residue:** 0.641
**Clashscore:** 5.150

**Description:** RhoBAST
**Organism:** artificial sequences

---

**Zhang, Y., Xu, Z., Xiao, Y., Jiang, H., Zuo, X., Li, X., Fang, X. (2024) Structural mechanisms for binding and activation of a contact-quenched fluorophore by RhoBAST. Nat Commun.**
**DOI:** 10.1038/s41467-024-48478-9

---

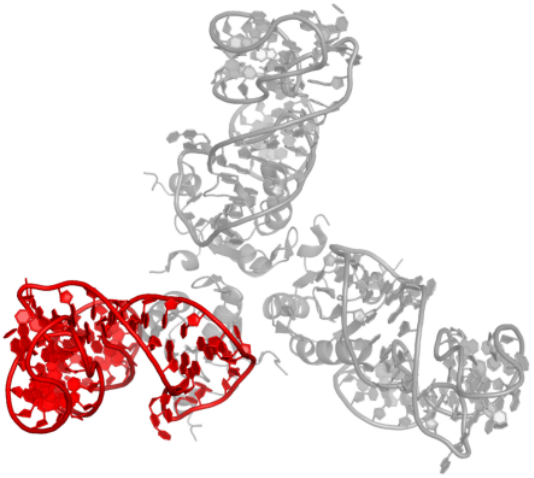

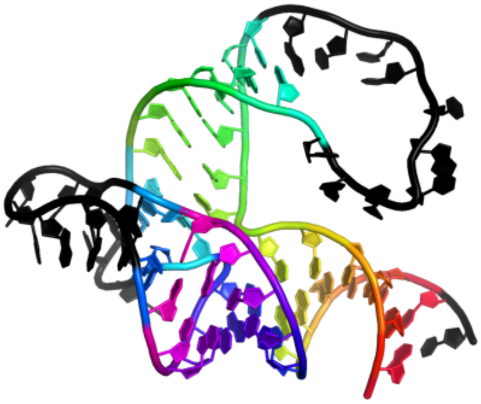

```
GAACCUCCGCCCAUUGCACUCCGGGCGGUGAAGGAGAGGCGCAAGGUUAACCGCCUCAGGUUCC
((((((((((((..........))))))(<...<)((((((>........>)))))))))))).
```

#### 1.4 Spinach aptamer

### 1.4.1 6b14_R

Crystal structure of Spinach RNA aptamer in complex with Fab BL3-6S97N

**Release date:** 2017-12-27
**Method:** x-ray diffraction
**Resolution:** 1.64 Å
**Chain length:** 83
**Extracted length:** 83

**Total clashes:** 0
**Clashes per residue:** 0.000
**Clashscore:** 2.380

**Description:** RNA (86-MER)
**Organism:** synthetic construct

---

**Koirala, D., Shelke, S.A., Dupont, M., Ruiz, S., DasGupta, S., Bailey, L.J., Benner, S.A., Piccirilli, J.A. (2018) Affinity maturation of a portable Fab-RNA module for chaperone-assisted RNA crystallography. Nucleic Acids Res.**
**DOI:** 10.1093/nar/gkx1292

---

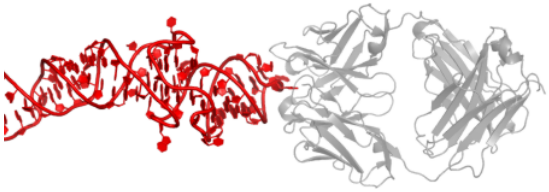

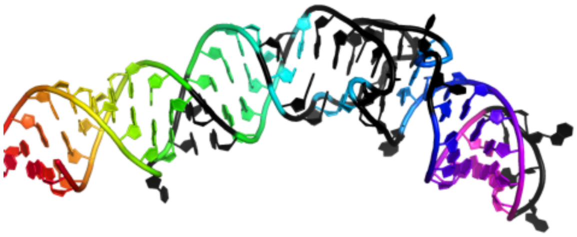

GACGCGACCGAAAUGGUGAAGGACGGGUCCAGUGCGAAACACGCACUGUUGAGUAGAGUGUGAGCUCCGUAACUGGUCGCGUC
((((((((((..((((.(.....(...(((((((((.....)))))))..)).........)..).))))..).))))))))))

### 1.4.2 6b3k_R

Crystal structure of mutant Spinach RNA aptamer in complex with Fab BL3-6

**Release date:** 2017-12-27
**Method:** x-ray diffraction
**Resolution:** 2.09 Å
**Chain length:** 83
**Extracted length:** 83

**Total clashes:** 0
**Clashes per residue:** 0.000
**Clashscore:** 4.330

**Description:** RNA (83-MER),RNA (83-MER)
**Organism:** synthetic construct

---

**Koirala, D., Shelke, S.A., Dupont, M., Ruiz, S., DasGupta, S., Bailey, L.J., Benner, S.A., Piccirilli, J.A. (2018) Affinity maturation of a portable Fab-RNA module for chaperone-assisted RNA crystallography. Nucleic Acids Res.**
**DOI:** 10.1093/nar/gkx1292

---

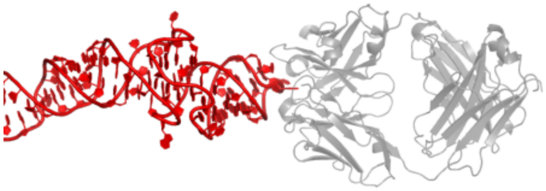

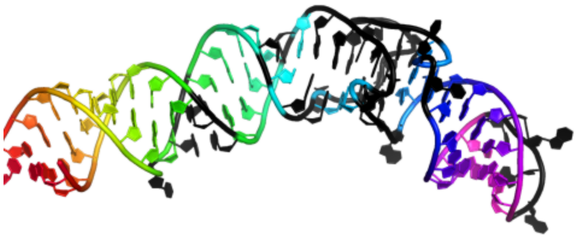

```
GACGCGACCGAAAUGGUGAAGGACGGGUCCAGUGCGAGACCCGCACUGUUGAGUAGAGUGUGAGCUCCGUAACUGGUCGCGUC
((((((((((..((((.(.....(...(((((((((.....)))))))..))..........)..).))))..).)))))))))
```

#### 1.5 Mango aptamer

### 1.5.1 8u5t_A

Structure of Mango II variant aptamer bound to T01-6A-B

**Release date:** 2024-03-27
**Method:** x-ray diffraction
**Resolution:** 2.20 Å
**Chain length:** 36
**Extracted length:** 36

**Total clashes:** 0
**Clashes per residue:** 0.000
**Clashscore:** 2.450

**Description:** Mango II variant
**Organism:** synthetic construct

**Passalacqua, L.F.M., Ferre-D'Amare, A.R. (N/A) Structure of Mango II variant aptamer bound to T01-6A-B. To Be Published.**
**DOI:** nan

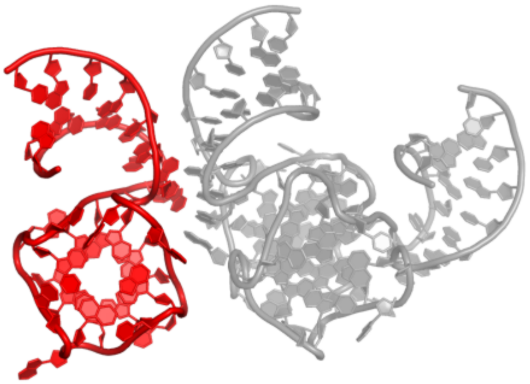

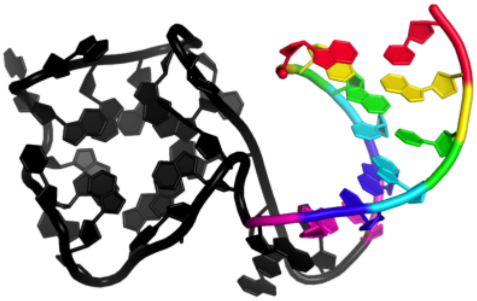

```
GCGUACGAAGGUGAGGAGAGGCGAGGAAGAGUACGC
((((((........................))))))
```

### 1.5.2 6e8u_B

Structure of the Mango-III (A10U) aptamer bound to TO1-Biotin

**Release date:** 2019-04-17
**Method:** x-ray diffraction
**Resolution:** 1.55 Å
**Chain length:** 37
**Extracted length:** 37

**Total clashes:** 0
**Clashes per residue:** 0.000
**Clashscore:** 4.540

**Description:** RNA (37-MER)
**Organism:** synthetic construct

---

**Trachman 3rd., R.J., Autour, A., Jeng, S.C.Y., Abdolahzadeh, A., Andreoni, A., Cojocaru, R., Garipov, R., Dolgosheina, E.V., Knutson, J.R., Ryckelynck, M., Unrau, P.J., Ferre-D'Amare, A.R. (2019) Structure and functional reselection of the Mango-III fluorogenic RNA aptamer. Nat. Chem. Biol.**
**DOI:** 10.1038/s41589-019-0267-9

---

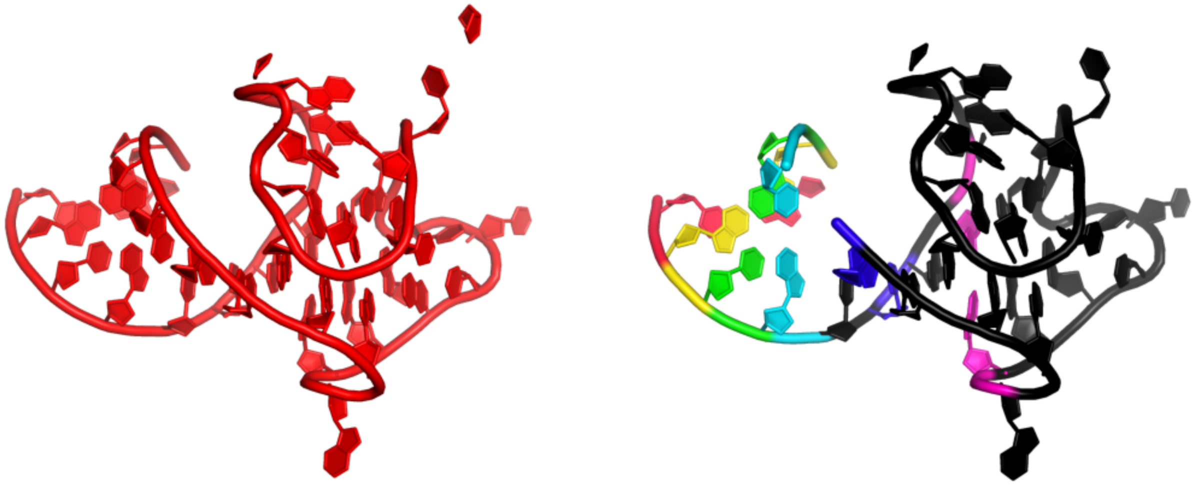

```
GUACGAAGGAAGGUUUGGUAUGGGGUAGUUGUCGUAC
((((.(....(..................).).))))
```

### 1.5.3 6e8t_D

Structure of the Mango-III (A10U) aptamer bound to TO1-Biotin

**Release date:** 2019-04-17
**Method:** x-ray diffraction
**Resolution:** 2.90 Å
**Chain length:** 36
**Extracted length:** 36

**Total clashes:** 31
**Clashes per residue:** 0.861
**Clashscore:** 12.160

**Description:** RNA (35-MER)
**Organism:** synthetic construct

---

**Trachman 3rd., R.J., Autour, A., Jeng, S.C.Y., Abdolahzadeh, A., Andreoni, A., Cojocaru, R., Garipov, R., Dolgosheina, E.V., Knutson, J.R., Ryckelynck, M., Unrau, P.J., Ferre-D'Amare, A.R. (2019) Structure and functional reselection of the Mango-III fluorogenic RNA aptamer. Nat. Chem. Biol.**
**DOI:** 10.1038/s41589-019-0267-9

---

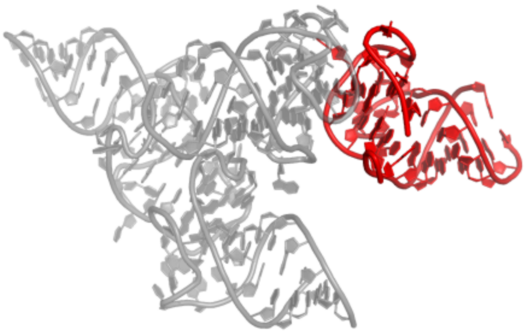

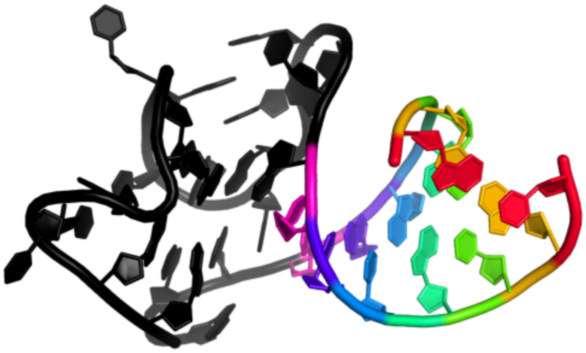

GUACGAAGGAAGGUUUGGUAUGUGGUAUAUUCGUAC
((((((......................))))))

### 1.5.4 8u5j_A

Structure of Mango III variant aptamer bound to T01-07M-B

**Release date:** 2024-03-27
**Method:** x-ray diffraction
**Resolution:** 1.70 Å
**Chain length:** 36
**Extracted length:** 36

**Total clashes:** 20
**Clashes per residue:** 0.556
**Clashscore:** 7.820

**Description:** Mango III variant
**Organism:** synthetic construct

---

**Passalacqua, L.F.M., Ferre-D'Amare, A.R. (N/A) Structure of Mango III variant aptamer bound to T01-07M-B. To Be Published.**
**DOI:** nan

---

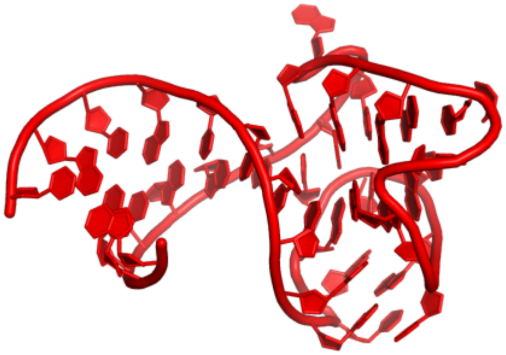

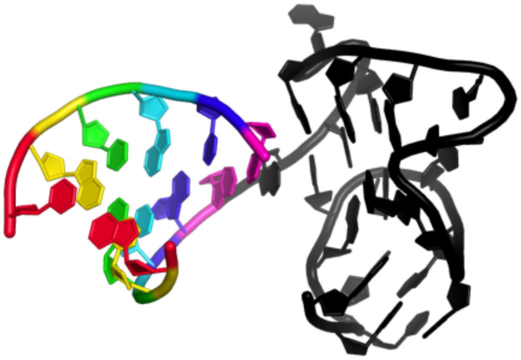

GUACGAAGGAAGGUUUGGUAUGUGGUAGAUUCGUAC
((((((........................))))))

### 1.5.5 6c65_B

Crystal Structure of the Mango-II-A22U Fluorescent Aptamer Bound to TO1-Biotin

**Release date:** 2018-08-08
**Method:** x-ray diffraction
**Resolution:** 2.80 Å
**Chain length:** 36
**Extracted length:** 36

**Total clashes:** 41
**Clashes per residue:** 1.139
**Clashscore:** 23.160

**Description:** RNA (36-MER)
**Organism:** synthetic construct

---

**Trachman 3rd., R.J., Abdolahzadeh, A., Andreoni, A., Cojocaru, R., Knutson, J.R., Ryckelynck, M., Unrau, P.J., Ferre-D'Amare, A.R. (2018) Crystal Structures of the Mango-II RNA Aptamer Reveal Heterogeneous Fluorophore Binding and Guide Engineering of Variants with Improved Selectivity and Brightness. Biochemistry.**
**DOI:** 10.1021/acs.biochem.8b00399

---

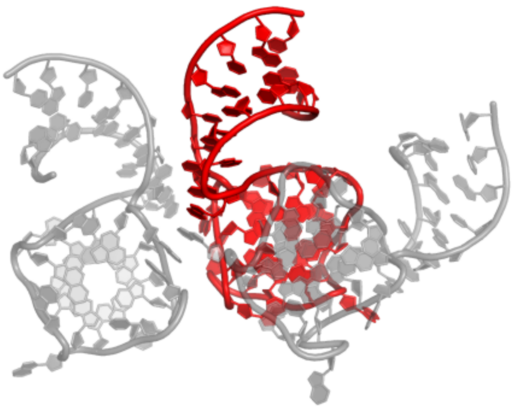

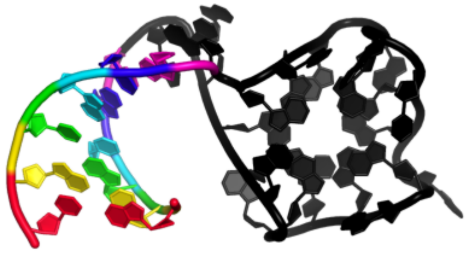

```
GCGUACGAAGGAGAGGAGAGGUAGAGGAGAGUACGC
((((((........................))))))
```

### 1.5.6 6e8s_B

Structure of the iMango-III aptamer bound to TO1-Biotin

**Release date:** 2019-04-17
**Method:** x-ray diffraction
**Resolution:** 2.35 Å
**Chain length:** 38
**Extracted length:** 38

**Total clashes:** 34
**Clashes per residue:** 0.895
**Clashscore:** 12.660

**Description:** iMango-III aptamer
**Organism:** synthetic construct

---

**Trachman 3rd., R.J., Autour, A., Jeng, S.C.Y., Abdolahzadeh, A., Andreoni, A., Cojocaru, R., Garipov, R., Dolgosheina, E.V., Knutson, J.R., Ryckelynck, M., Unrau, P.J., Ferre-D'Amare, A.R. (2019) Structure and functional reselection of the Mango-III fluorogenic RNA aptamer. Nat. Chem. Biol.**
**DOI:** 10.1038/s41589-019-0267-9

---

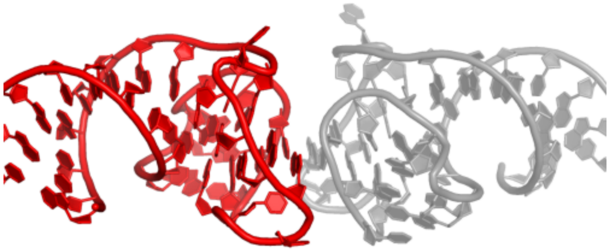

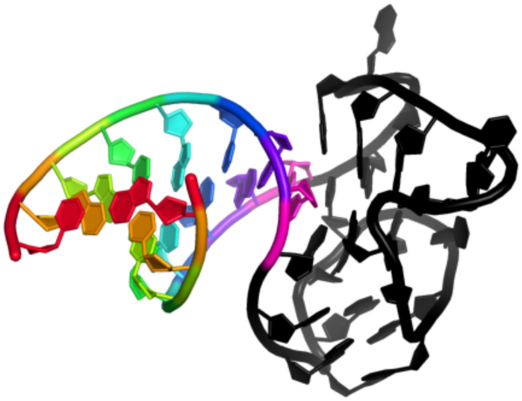

GCUACGAAGGAAGGAUUGGUAUGUGGUAUAUUCGUAGC
((((((((......................))))))))

#### 1.6 Vitamin B12 aptamer

### 1.6.1 1et4_E

CRYSTAL STRUCTURE OF A VITAMIN B12 BINDING RNA APTAMER WITH LIGAND AT 2.3 A

**Release date:** 2000-11-13
**Method:** x-ray diffraction
**Resolution:** 2.30 Å
**Chain length:** 35
**Extracted length:** 35

**Total clashes:** 24
**Clashes per residue:** 0.686
**Clashscore:** 10.070

**Description:** RNA APTAMER, 35-MER
**Organism:** nan

---

**Sussman, D., Wilson, C. (2000) A water channel in the core of the vitamin B(12) RNA aptamer. Structure Fold.Des.**
**DOI:** 10.1016/S0969-2126(00)00159-3

---

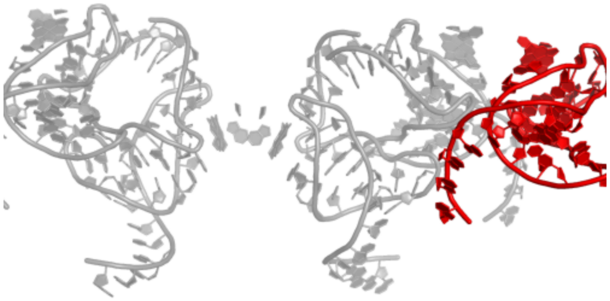

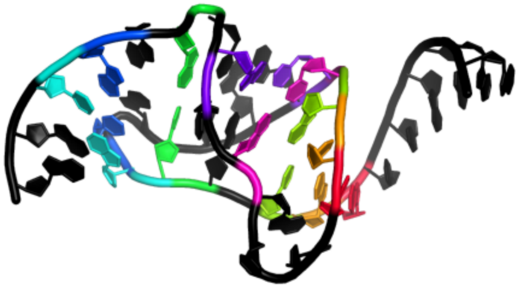

```
GGAACCGGUGCGCAUAACCACCUCAGUGCGAGCAA
......<<<.((....((>>>...).)..))..
```

#### 1.7 Squash aptamer

##### 1.7.1 7kvu_G

Crystal structure of Squash RNA aptamer in complex with DFHBI-1T

**Release date:** 2022-01-19
**Method:** x-ray diffraction
**Resolution:** 2.68 Å
**Chain length:** 83
**Extracted length:** 83

**Total clashes:** 0
**Clashes per residue:** 0.000
**Clashscore:** 3.680

**Description:** Squash RNA aptamer
**Organism:** synthetic construct

---

**Truong, L., Kooshapur, H., Dey, S.K., Li, X., Tjandra, N., Jaffrey, S.R., Ferre-D'Amare, A.R. (2022) The fluorescent aptamer Squash extensively repurposes the adenine riboswitch fold. Nat.Chem.Biol.**
**DOI:** 10.1038/s41589-021-00931-2

---

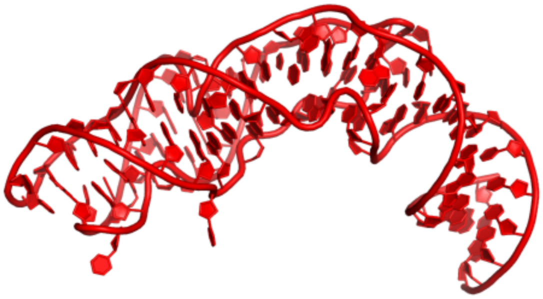

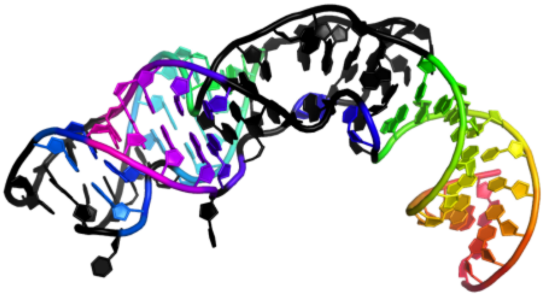

GGGAAGAUACAAGGUGAGCCCAAUAAUAUGGUUUGGGUUAGGAUAGGAAGUAGAGCCUUAAACUCUCUAAGCGGUAUCUUCCC
((((((((((.......((((((......<<.))))))........(..(.((((>>.....)))))....).))))))))))

#### 1.8 Pepper aptamer

##### 1.8.1 7eoh_A

Crystal structure of the Pepper aptamer in complex with HBC

**Release date:** 2021-11-24
**Method:** x-ray diffraction
**Resolution:** 1.64 Å
**Chain length:** 49
**Extracted length:** 49

**Total clashes:** 0
**Clashes per residue:** 0.000
**Clashscore:** 0.620

**Description:** Pepper (49-MER)
**Organism:** synthetic construct

---

**Huang, K., Chen, X., Li, C., Song, Q., Li, H., Zhu, L., Yang, Y., Ren, A. (2021) Structure-based investigation of fluorogenic Pepper aptamer. Nat.Chem.Biol.**
**DOI:** 10.1038/s41589-021-00884-6

---

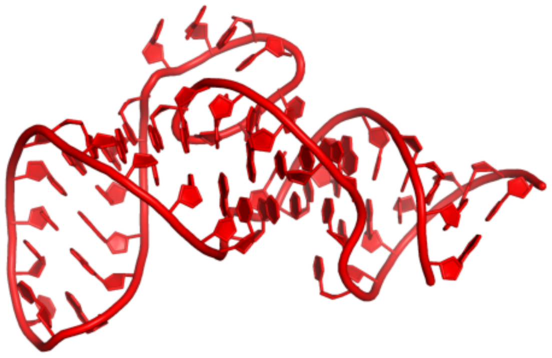

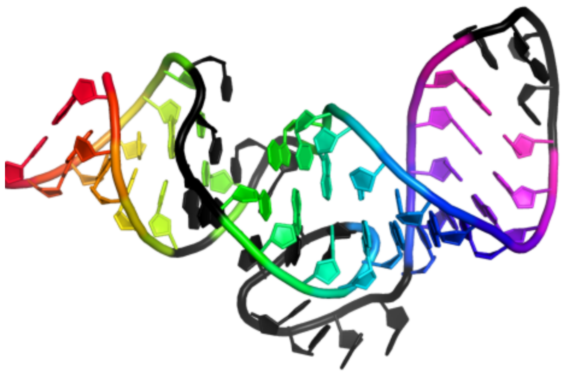

```
GGCGCACUGGCGCUGCGCCUUCGGGCGCCAAUCGUAGCGUGUCGGCGCC
(((((.(..((((((((((....))))).......)))))...))))))
```

#### 1.9 Chili aptamer

##### 1.9.1 7oax_D

Crystal structure of the Chili RNA aptamer in complex with DMHBO+

**Release date:** 2021-06-16
**Method:** x-ray diffraction
**Resolution:** 2.24 Å
**Chain length:** 52
**Extracted length:** 52

**Total clashes:** 4
**Clashes per residue:** 0.077
**Clashscore:** 1.550

**Description:** Chili RNA Aptamer
**Organism:** synthetic construct

---

**Mieczkowski, M., Steinmetzger, C., Bessi, I., Lenz, A.K., Schmiedel, A., Holzapfel, M., Lambert, C., Pena, V., Hobartner, C. (2021) Large Stokes shift fluorescence activation in an RNA aptamer by intermolecular proton transfer to guanine. Nat Commun.**
**DOI:** 10.1038/s41467-021-23932-0

---

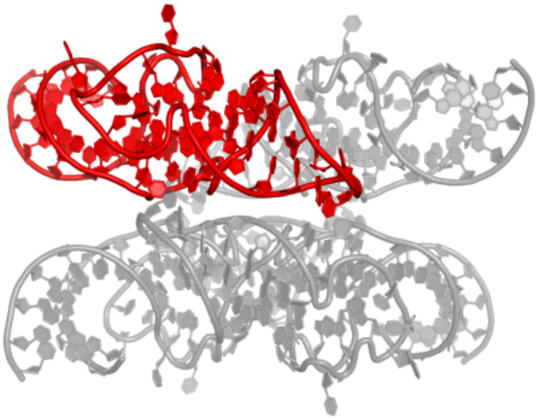

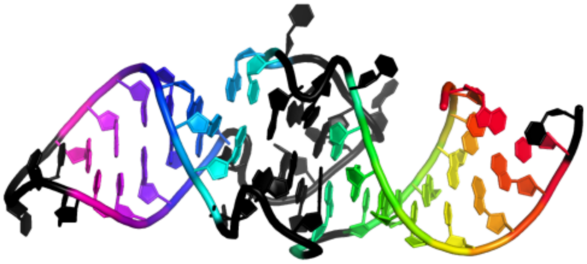

```
GGCUAGCUGGAGGGGCGCCAGUUCGCUGGUGGUUGGGUGCGGUCGGCUAGCC
.(((((((.(....(((((((....)))))........))...)))))))).
```

#### 1.10 Corn aptamer

##### 1.10.1 5bjo_Y

Crystal structure of the Corn RNA aptamer in complex with DFHO, site-specific 5-iodo-U

**Release date:** 2017-09-27
**Method:** x-ray diffraction
**Resolution:** 2.35 Å
**Chain length:** 36
**Extracted length:** 36

**Total clashes:** 0
**Clashes per residue:** 0.000
**Clashscore:** 4.190

**Description:** RNA (36-MER)
**Organism:** synthetic construct

**Warner, K.D., Sjekloca, L., Song, W., Filonov, G.S., Jaffrey, S.R., Ferre-D'Amare, A.R. (2017) A homodimer interface without base pairs in an RNA mimic of red fluorescent protein. Nat. Chem. Biol.**
**DOI:** 10.1038/nchembio.2475

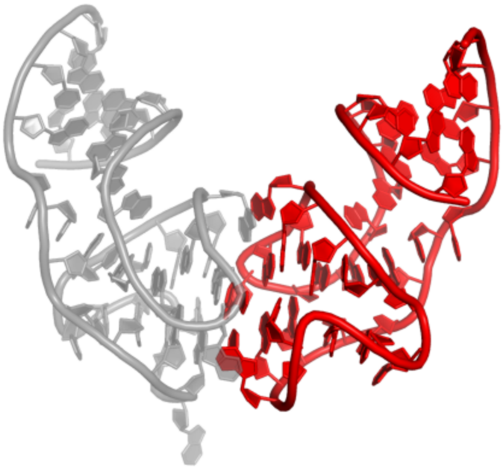

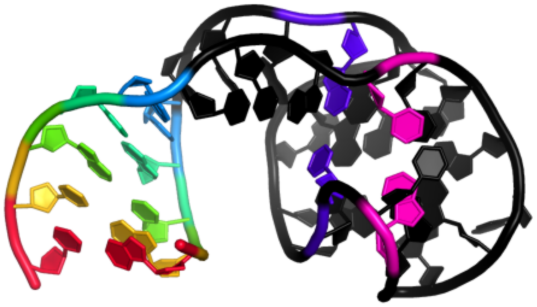

GGCGCGAGGAAGGAGGUCUGAGGAGGUCACUGCGCC
(((((...(........).(.......)...)))))

#### 1.11 DIR2s aptamer

### 1.11.1 6db8_R

Structural basis for promiscuous binding and activation of fluorogenic dyes by DIR2s RNA aptamer

**Release date:** 2018-11-14
**Method:** x-ray diffraction
**Resolution:** 1.87 Å
**Chain length:** 60
**Extracted length:** 60

**Total clashes:** 0
**Clashes per residue:** 0.000
**Clashscore:** 4.480

**Description:** RNA (60-MER)
**Organism:** synthetic construct

---

**Shelke, S.A., Shao, Y., Laski, A., Koirala, D., Weissman, B.P., Fuller, J.R., Tan, X., Constantin, T.P., Waggoner, A.S., Bruchez, M.P., Armitage, B.A., Piccirilli, J.A. (2018) Structural basis for activation of fluorogenic dyes by an RNA aptamer lacking a G-quadruplex motif. Nat Commun.**
**DOI:** 10.1038/s41467-018-06942-3

---

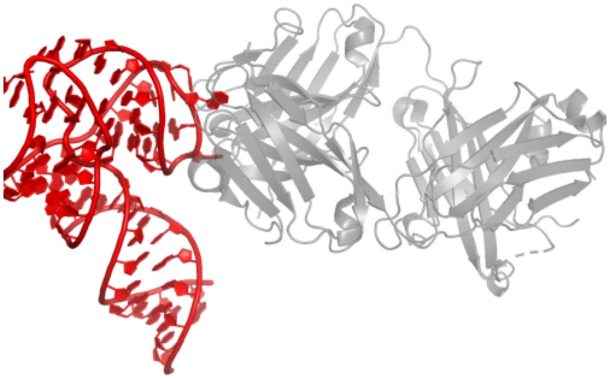

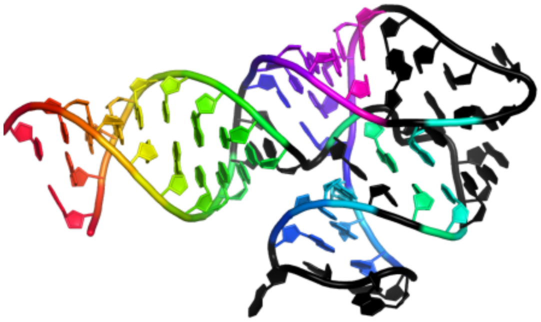

GGAUGCGCCUUGAAAAGCCUGCGAAACACGCAGCUGGUGAAUGACAGCUAUGGCGCAUCC
(((((((((..(.....<).(((.....)))(((((......>.)))))..)))))))))

#### 1.12 Clivia aptamer

##### 1.12.1 8hze_B

A new fluorescent RNA aptamer bound with N

**Release date:** 2024-06-19
**Method:** x-ray diffraction
**Resolution:** 1.59 Å
**Chain length:** 35
**Extracted length:** 35

**Total clashes:** 0
**Clashes per residue:** 0.000
**Clashscore:** 2.480

**Description:** RNA (36-MER)
**Organism:** synthetic construct

**Huang, K., Song, Q., Fang, M., Yao, D., Shen, X., Xu, X., Chen, X., Zhu, L., Yang, Y., Ren, A. (2024) Structural basis of a small monomeric Clivia fluorogenic RNA with a large Stokes shift. Nat.Chem.Biol.**
**DOI:** 10.1038/s41589-024-01633-1

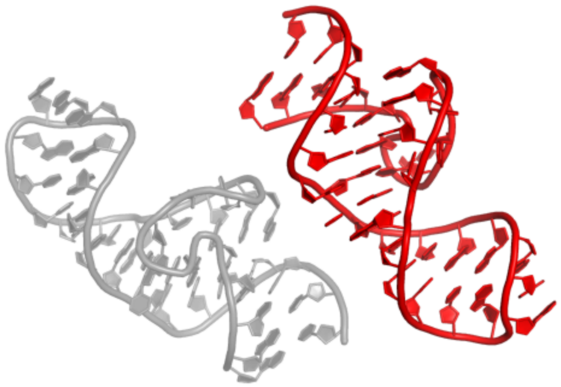

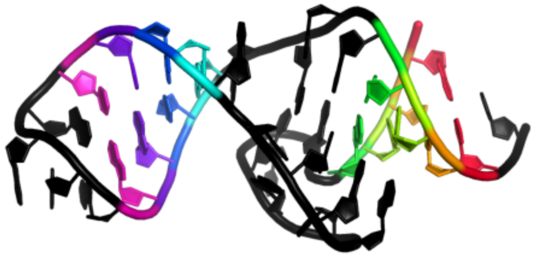

```
GAAGAUUGUAAACAUGCCGAAAGGCAGACACUUCC
((((..........((((....))))....)))).
```

##### 1.12.2 8hzj_A

A new fluorescent RNA aptamer bound with N571

**Release date:** 2024-06-19
**Method:** x-ray diffraction
**Resolution:** 2.60 Å
**Chain length:** 35
**Extracted length:** 35

**Total clashes:** 2
**Clashes per residue:** 0.057
**Clashscore:** 2.500

**Description:** RNA (36-MER)
**Organism:** synthetic construct

**Huang, K., Song, Q., Fang, M., Yao, D., Shen, X., Xu, X., Chen, X., Zhu, L., Yang, Y., Ren, A. (2024) Structural basis of a small monomeric Clivia fluorogenic RNA with a large Stokes shift. Nat.Chem.Biol.**
**DOI:** 10.1038/s41589-024-01633-1

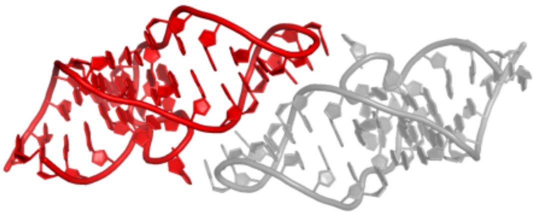

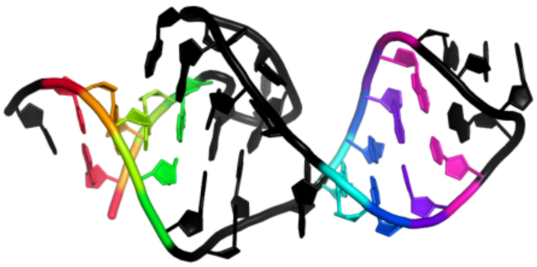

GAAGAUUGUAAACAUGCCGAAAGGCAGACACUUCC
((((..........((((....))))....)))).

##### 1.12.3 8hzl_B

A new fluorescent RNA aptamer_III bound with N

**Release date:** 2024-06-19
**Method:** x-ray diffraction
**Resolution:** 2.60 Å
**Chain length:** 84
**Extracted length:** 84

**Total clashes:** 26
**Clashes per residue:** 0.310
**Clashscore:** 5.170

**Description:** RNA (84-MER)
**Organism:** synthetic construct

---

**Huang, K., Song, Q., Fang, M., Yao, D., Shen, X., Xu, X., Chen, X., Zhu, L., Yang, Y., Ren, A. (2024) Structural basis of a small monomeric Clivia fluorogenic RNA with a large Stokes shift. Nat.Chem.Biol.**
**DOI:** 10.1038/s41589-024-01633-1

---

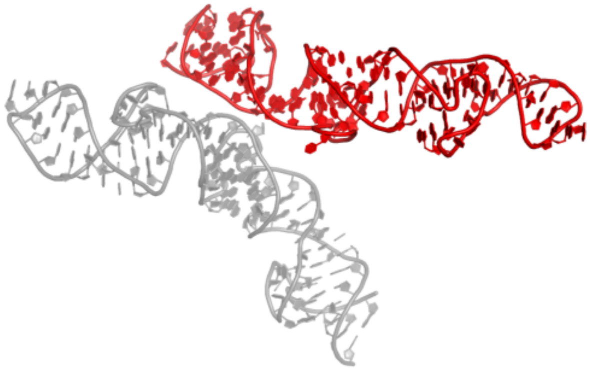

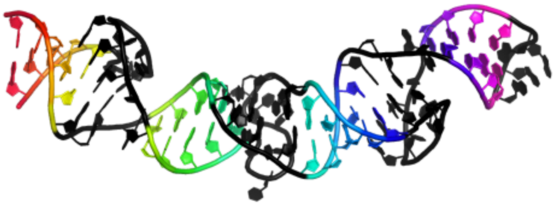

GGAAGAUUGUAAACAGCGAGAUUGUAAACAGCGAGAUUGUAAACAUGCCGAAAGGCAGACACUCGCGACACUCGCGACACUUUC
(((((..........(((((..........(((((..........((((....))))....)))))....)))))....)))))

#### 1.13 A9g aptamer

##### 1.13.1 6rti_X

X-ray structure of human glutamate carboxypeptidase II (GCPII) in complex with aptamer A9g

**Release date:** 2020-06-10
**Method:** x-ray diffraction
**Resolution:** 2.20 Å
**Chain length:** 43
**Extracted length:** 43

**Total clashes:** 0
**Clashes per residue:** 0.000
**Clashscore:** 3.130

**Description:** Aptamer A9g, RNA (43-MER)
**Organism:** synthetic construct

---

**Ptacek, J., Zhang, D., Qiu, L., Kruspe, S., Motlova, L., Kolenko, P., Novakova, Z., Shubham, S., Havlinova, B., Baranova, P., Chen, S.J., Zou, X., Giangrande, P., Barinka, C. (2020) Structural basis of prostate-specific membrane antigen recognition by the A9g RNA aptamer. Nucleic Acids Res.**
**DOI:** 10.1093/nar/gkaa494

---

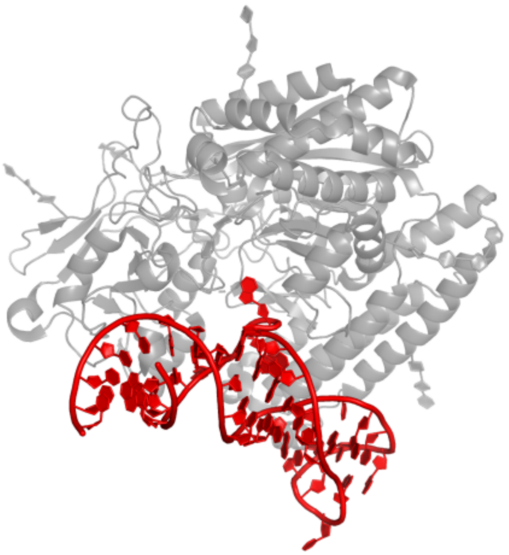

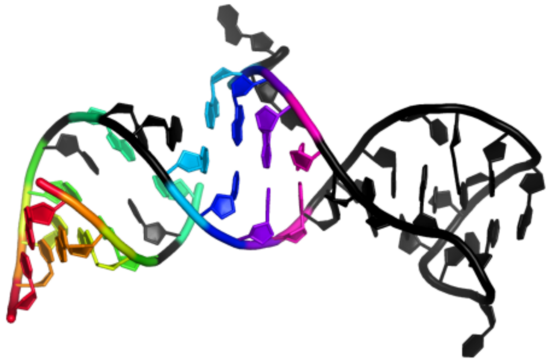

GGGACCGAAAAAGACCUGACUUCUAUACUAAGUCUACGUUCCC
((((.(.....((((................))))..).))))

#### 1.14 Beetroot aptamer

##### 1.14.1 8eyu_B

Structure of Beetroot dimer bound to DFAME

**Release date:** 2023-05-31
**Method:** x-ray diffraction
**Resolution:** 1.95 Å
**Chain length:** 49
**Extracted length:** 49

**Total clashes:** 5
**Clashes per residue:** 0.102
**Clashscore:** 3.240

**Description:** RNA (49-MER)
**Organism:** synthetic construct

**Passalacqua, L.F.M., Starich, M.R., Link, K.A., Wu, J., Knutson, J.R., Tjandra, N., Jaffrey, S.R., Ferre-D'Amare, A.R. (2023) Co-crystal structures of the fluorogenic aptamer Beetroot show that close homology may not predict similar RNA architecture. Nat Commun.**
**DOI:** 10.1038/s41467-023-38683-3

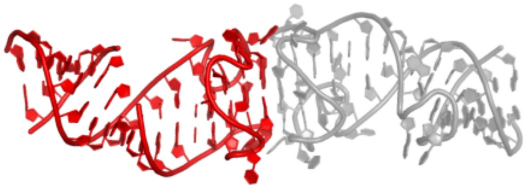

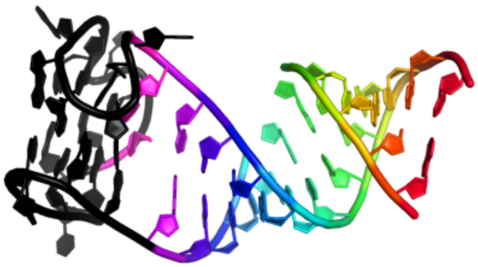

GCGCCGGUUAGGCAGAGGUGGGUGGUGUGGAGGAGUAUCUGUCCGGCGC
((((((....((((((.........(..........)))))))))))))

### 1.14.2 8f0n_B

Wobble Beetroot (A16U-U38G) dimer bound to DFHO

**Release date:** 2023-05-31
**Method:** x-ray diffraction
**Resolution:** 2.85 Å
**Chain length:** 49
**Extracted length:** 49

**Total clashes:** 26
**Clashes per residue:** 0.531
**Clashscore:** 10.100

**Description:** RNA (49-MER)
**Organism:** synthetic construct

---

**Passalacqua, L.F.M., Starich, M.R., Link, K.A., Wu, J., Knutson, J.R., Tjandra, N., Jaffrey, S.R., Ferre-D'Amare, A.R. (2023) Co-crystal structures of the fluorogenic aptamer Beetroot show that close homology may not predict similar RNA architecture. Nat Commun.**
**DOI:** 10.1038/s41467-023-38683-3

---

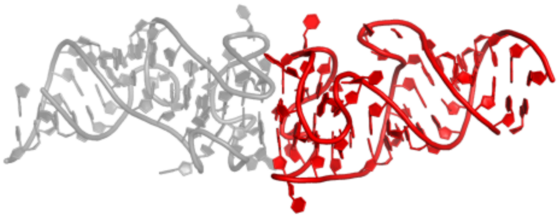

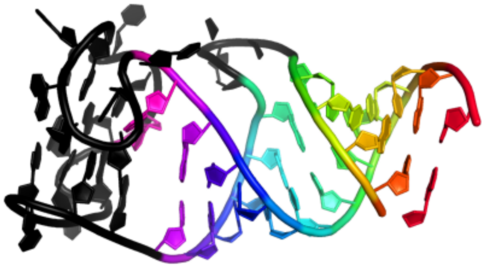

GCGCCGGUUAGGCAGUGGUGGGUGGUGUGGAGGAGUAGCUGUCCGGCGC
((((((....((((((.........(..........))))))))))))

#### 1.15 Malachite green aptamer

### 1.15.1 1f1t_A

CRYSTAL STRUCTURE OF THE MALACHITE GREEN APTAMER COMPLEXED WITH TETRAMETHYL-ROSAMINE

**Release date:** 2000-09-04
**Method:** x-ray diffraction
**Resolution:** 2.80 Å
**Chain length:** 38
**Extracted length:** 38

**Total clashes:** 92
**Clashes per residue:** 2.421
**Clashscore:** 35.830

**Description:** MALACHITE GREEN APTAMER RNA
**Organism:** nan

---

**Baugh, C., Grate, D., Wilson, C. (2000) 2.8 A crystal structure of the malachite green aptamer. J.Mol.Biol.**
**DOI:** 10.1006/jmbi.2000.3951

---

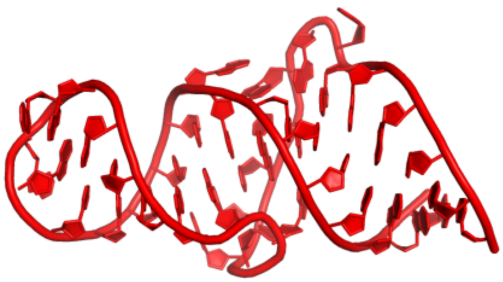

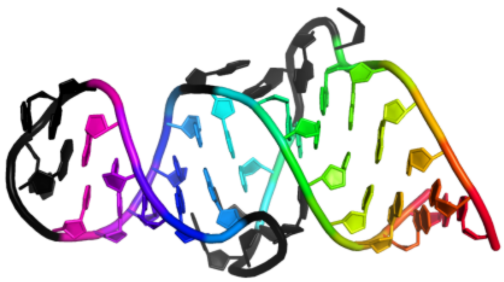

GGAUCCCGACUGGCGAGAGCCAGGUAACGAAUGGAUCC
((((((((.(((((....)))))....))...))))))

#### 1.16 11F7t aptamer

##### 1.16.1 5voe_A

DesGla-XaS195A Bound to Aptamer 11F7t

**Release date:** 2018-06-20
**Method:** x-ray diffraction
**Resolution:** 2.00 Å
**Chain length:** 36
**Extracted length:** 36

**Total clashes:** 0
**Clashes per residue:** 0.000
**Clashscore:** 12.260

**Description:** Aptamer 11F7t (36-MER)
**Organism:** synthetic construct

---

**Gunaratne, R., Kumar, S., Frederiksen, J.W., Stayrook, S., Lohrmann, J.L., Perry, K., Bompiani, K.M., Chabata, C.V., Thalji, N.K., Ho, M.D., Arepally, G., Camire, R.M., Krishnaswamy, S., Sullenger, B.A. (2018) Combination of aptamer and drug for reversible anticoagulation in cardiopulmonary bypass. Nat. Biotechnol.**
**DOI:** 10.1038/nbt.4153

---

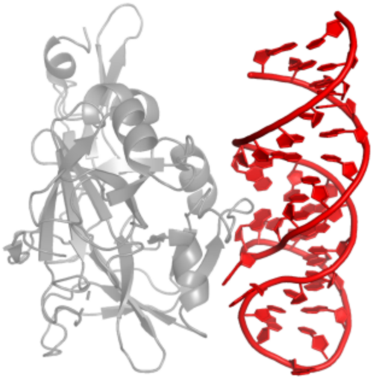

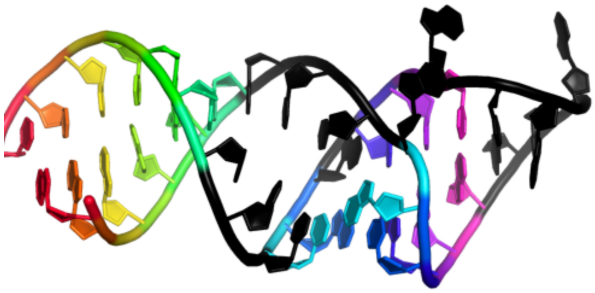

GAGAGCCCCAGCGAGAUAAUACUUGGCCCCGCUCUU
((((((....(((((......))).))...))))))

#### 1.17 K1 aptamer

### 1.17.1 6sy4_C

TetR in complex with the TetR-binding RNA-aptamer K1

**Release date:** 2020-02-05
**Method:** x-ray diffraction
**Resolution:** 2.69 Å
**Chain length:** 43
**Extracted length:** 38

**Total clashes:** 4
**Clashes per residue:** 0.093
**Clashscore:** 3.720

**Description:** TetR-binding aptamer K1 (43-MER)
**Organism:** Escherichia coli

**Grau, F.C., Jaeger, J., Groher, F., Suess, B., Muller, Y.A. (2020) The complex formed between a synthetic RNA aptamer and the transcription repressor TetR is a structural and functional twin of the operator DNA-TetR regulator complex. Nucleic Acids Res.**
**DOI:** 10.1093/nar/gkaa083

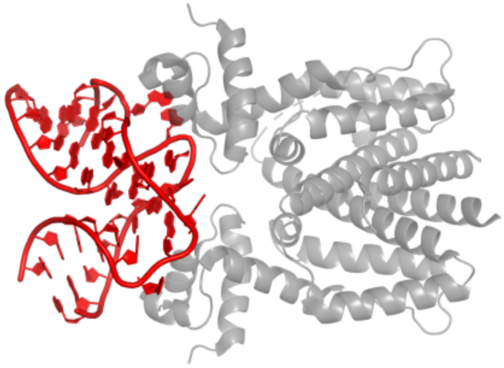

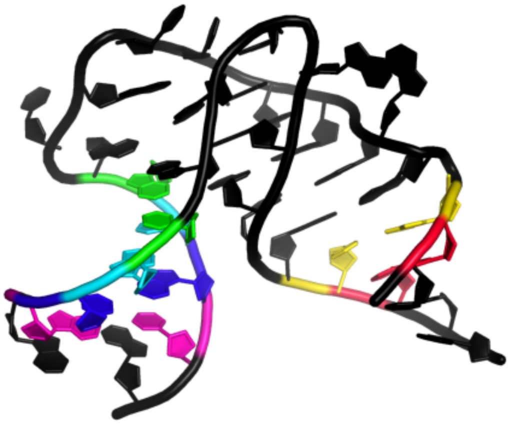

GGCCGGAGAAUGUUAUGGCGCGAAAGCGCAGAGAAAACCGGUC
....((...........((((....))))........))....

#### 1.18 Tetracycline aptamer

##### 1.18.1 3egz_B

Crystal structure of an in vitro evolved tetracycline aptamer and artificial riboswitch

**Release date:** 2008-10-28
**Method:** x-ray diffraction
**Resolution:** 2.20 Å
**Chain length:** 65
**Extracted length:** 65

**Total clashes:** 136
**Clashes per residue:** 2.092
**Clashscore:** 24.650

**Description:** Tetracycline aptamer and artificial riboswitch
**Organism:** nan

---

**Xiao, H., Edwards, T.E., Ferre-D'Amare, A.R. (2008) Structural basis for specific, high-affinity tetracycline binding by an in vitro evolved aptamer and artificial riboswitch. Chem.Biol.**
**DOI:** 10.1016/j.chembiol.2008.09.004

---

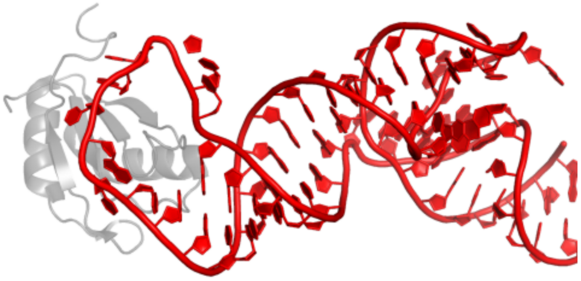

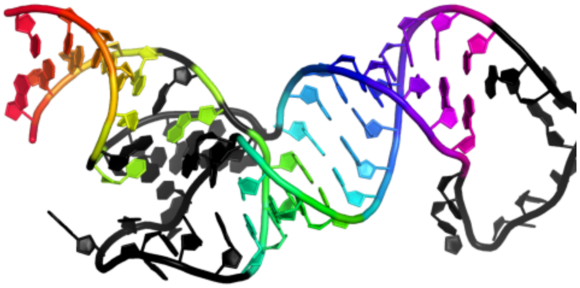

GAGGGAGAGGUGAAGAAUACGACCACCUAGGUACCAUUGCACUCCGGUACCUAAAACAUACCCUC
(((((.(.((((...........))))((((((((..........)))))))).....).)))))

### 2 CRISPR guides

#### 2.1 Cas9 guide

### 2.1.1 7el1_B

Structure of a protein from bacteria

**Release date:** 2021-07-28
**Method:** x-ray diffraction
**Resolution:** 2.22 Å
**Chain length:** 73
**Extracted length:** 53

**Total clashes:** 9
**Clashes per residue:** 0.123
**Clashscore:** 6.160

**Description:** RNA (73-MER)
**Organism:** Staphylococcus aureus

---

**Liu, H., Zhu, Y., Lu, Z., Huang, Z. (2021) Structural basis of Staphylococcus aureus Cas9 inhibition by AcrIIA14. Nucleic Acids Res.**
**DOI:** 10.1093/nar/gkab487

---

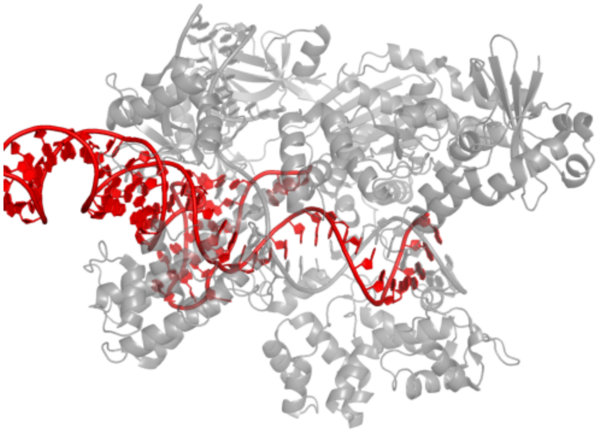

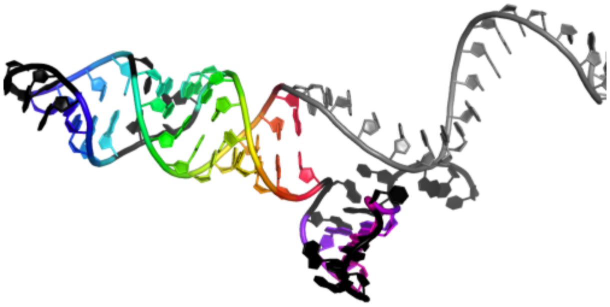

GGAAAUUAGGUGCGCUUGGCGUUUUAGUACUCUGGAAACAGAAUCUACUAAAACAAGGCAAAAUGCCGUGUUU
....................(((((((((.((((....))))...)))))))))..(((.....)))......

##### 2.1.2 6wbr_B

Crystal structure of AceCas9 bound with guide RNA and DNA with 5'-NNNCC-3' PAM

**Release date:** 2020-11-18
**Method:** x-ray diffraction
**Resolution:** 2.91 Å
**Chain length:** 94
**Extracted length:** 74

**Total clashes:** 73
**Clashes per residue:** 0.777
**Clashscore:** 11.650

**Description:** RNA (94-MER)
**Organism:** Acidothermus cellulolyticus 11B

---

**Das, A., Hand, T.H., Smith, C.L., Wickline, E., Zawrotny, M., Li, H. (2020) The molecular basis for recognition of 5'-NNNCC-3' PAM and its methylation state by Acidothermus cellulolyticus Cas9. Nat Commun.**
**DOI:** 10.1038/s41467-020-20204-1

---

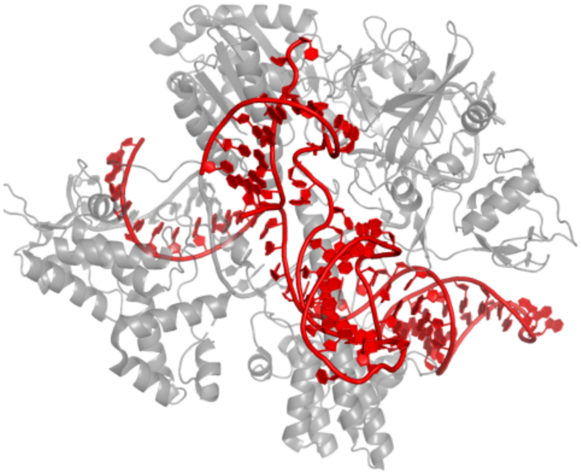

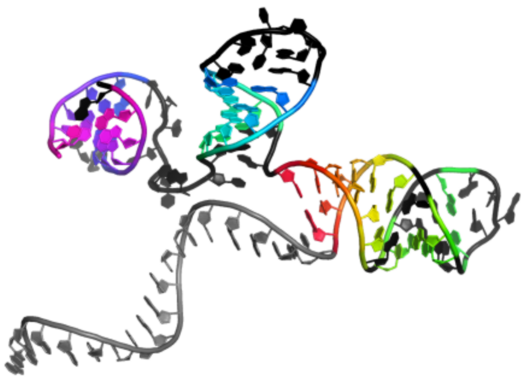

GGAUGGCAAGAUCCUGGUAUGCUGGGGAGCCUGAAAAGGCUACCUAGCAAGACCCCUUCGUGGGGUCGCAUUCUUCACCCCCAGCAGGGGGUUC
....................((((((.((..(....)..)).))))))..(((((.......))))).........((((((....))))))..

##### 2.1.3 8umf_B

Structure of PsCas9 in complex with gRNA and DNA in product state

**Release date:** 2024-10-02
**Method:** electron microscopy
**Resolution:** 2.90 Å
**Chain length:** 131
**Extracted length:** 101

**Total clashes:** 50
**Clashes per residue:** 0.382
**Clashscore:** 4.930

**Description:** RNA (121-MER)
**Organism:** synthetic construct

---

**Bravo, J.P.K., Taylor, D.W. (N/A) Structure-guided engineering of PsCas9 yields a high fidelity and activity enzyme for in vivo gene editing. To Be Published.**
**DOI:** nan

---

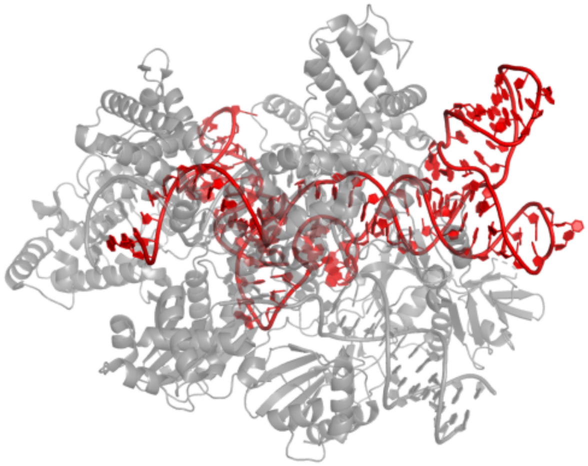

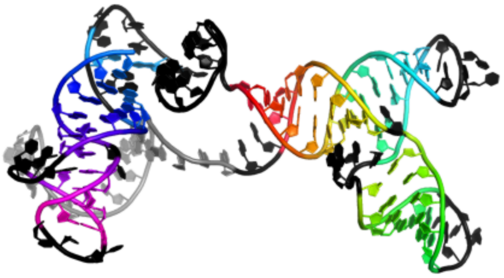

AUGUCACCUCCAAUGACUAGGGGUUUCAGUUAUUCGUGAAAACGAAUGAAGUCACUCUUAAAGUGAGCUGAAAUCACUAAAAAUUAAGAUUGAACCCGGCUACUGACUCUGUCAUCCGGGUUUACUUAUUU

......................((((((((.((((((....))))))....(((((.....)))))))))))))...........(....)..((((((....((((...)))).)))))))..........

##### 2.1.4 8hud_B

Cryo-EM structure of the EvCas9-sgRNA-target DNA ternary complex

**Release date:** 2023-12-27
**Method:** electron microscopy
**Resolution:** 3.43 Å
**Chain length:** 75
**Extracted length:** 52

**Total clashes:** 19
**Clashes per residue:** 0.253
**Clashscore:** 10.270

**Description:** sgRNA
**Organism:** synthetic construct

---

**Tang, N., Wu, Z., Gao, Y., Chen, W., Wang, Z., Su, M., Ji, W., Ji, Q. (2024) Molecular Basis and Genome Editing Applications of a Compact Eubacterium ventriosum CRISPR-Cas9 System. Acs Synth Biol.**
**DOI:** 10.1021/acssynbio.3c00501

---

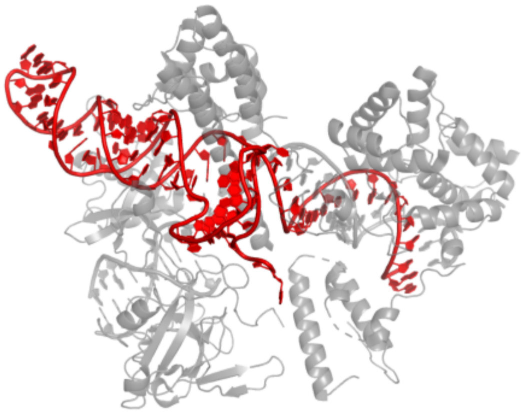

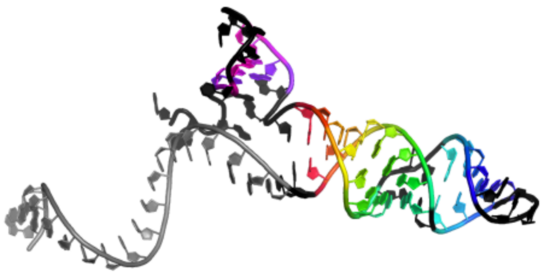

GGUAAUCGCUCUCCUCCGGCGAUUUUAGUACCUGAGAAAUCAGAUCUACUAAAACAAGGCUUUAUGCCGAAAUCA
......................((((((((.((((....))))...))))))))...(((.....))).......

#### 2.2 sgRNA guide

##### 2.2.1 8rdu_1

Conformational Landscape of the Type V-K CRISPR-associated TransposonIntegration Assembly CAST V-K composite map

**Release date:** 2024-06-19
**Method:** electron microscopy
**Resolution:** 2.30 Å
**Chain length:** 261
**Extracted length:** 227

**Total clashes:** 16
**Clashes per residue:** 0.061
**Clashscore:** 3.440

**Description:** sgRNA
**Organism:** Scytonema hofmannii

**Tenjo-Castano, F., Sofos, N., Stutzke, L.S., Temperini, P., Fuglsang, A., Pape, T., Mesa, P., Montoya, G. (2024) Conformational landscape of the type V-K CRISPR-associated transposon integration assembly. Mol.Cell.**
**DOI:** 10.1016/j.molcel.2024.05.005

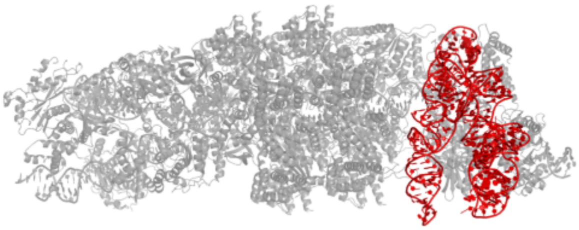

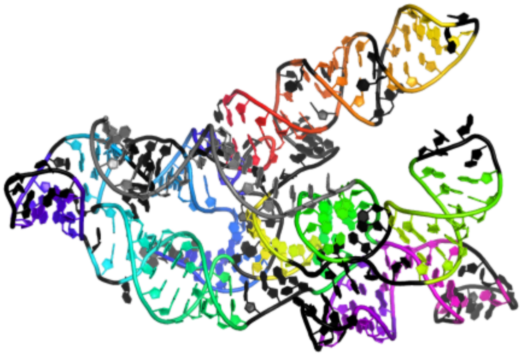

GGAUAUUAAUAGCGCCGCAAUUCAUGCUGCUUGCAGCCUCUGAAUUUUGUUAAAUGAGGGUUAGUUUGACUGUAUAAAUACAGUCUUGCUUUCUGACCCUGGUAGCUGCUCACCCUGAUGCUGCUGUCAAUAGACAGGAUAGGUGCGCUCCCAGCAAUAAGGGCGCGGAUGUACUGCUGUAGUGGCUACUGAAUCACCCCCGAUCAAGGGGGAACCCUCCAAAAGGUGGGUUGAAAGGAGAAGUCAUUUAAUAAGGCCACU

...........<<<<((.(.(((..(((((..)))))....))).).))........<<<<......((((((....))))))...(((((((>>>>.((((((((((.(((....(((((((((....)))))....((.>>>>.)))))).....)).((((.....)))).).))))))))))....))(((((......)))))(((((..(....)..))))))))))........................

### 2.2.2 8x5v_B

BlCas9-sgRNA-target DNA complex

**Release date:** 2024-07-10
**Method:** x-ray diffraction
**Resolution:** 2.00 Å
**Chain length:** 110
**Extracted length:** 91

**Total clashes:** 2
**Clashes per residue:** 0.018
**Clashscore:** 0.940

**Description:** RNA (110-mer)
**Organism:** Brevibacillus laterosporus

**Nakane, T., Nakagawa, R., Ishiguro, S., Okazaki, S., Mori, H., Shuto, Y., Yamashita, K., Yachie, N., Nishimasu, H., Nureki, O. (2024) Structure and engineering of Brevibacillus laterosporus Cas9. Commun Biol.**
**DOI:** 10.1038/s42003-024-06422-z

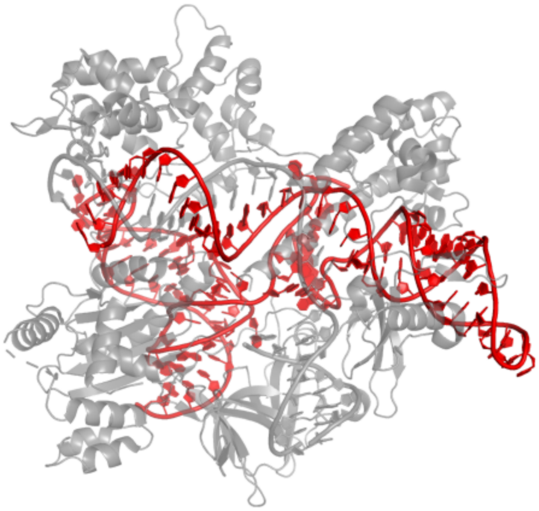

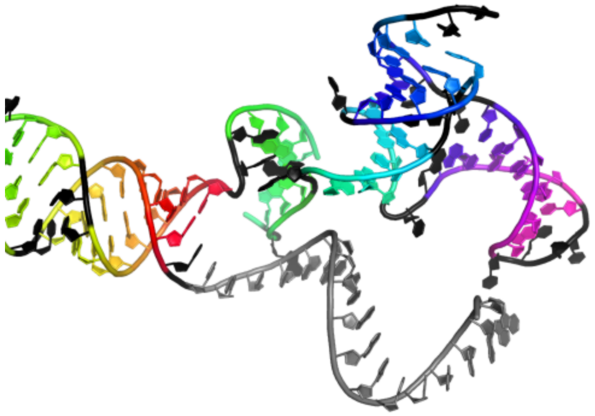

```
GGAAAUUAGGUGCGCUUGGCGCUAUAGUUCCUUGAAAAAGUUGCUAUAGUAAGGGCAACAGACCCGAGGCGUUGGGGAUCGCCUAGCCCGUUUUUACGGGCUCUCCCCAU
....................((((((((..(((....)))..))))))))..((((....).)))..<<<<..(((((.>>>>..((((((....))))))..)))))..
```

### 2.2.3 7c7l_C

Cryo-EM structure of the Cas12f1-sgRNA-target DNA complex

**Release date:** 2020-12-23
**Method:** electron microscopy
**Resolution:** 3.30 Å
**Chain length:** 180
**Extracted length:** 110

**Total clashes:** 29
**Clashes per residue:** 0.161
**Clashscore:** 4.790

**Description:** sgRNA
**Organism:** uncultured archaeon

**Takeda, S.N., Nakagawa, R., Okazaki, S., Hirano, H., Kobayashi, K., Kusakizako, T., Nishizawa, T., Yamashita, K., Nishimasu, H., Nureki, O. (2021) Structure of the miniature type V-F CRISPR-Cas effector enzyme. Mol.Cell.**
**DOI:** 10.1016/j.molcel.2020.11.035

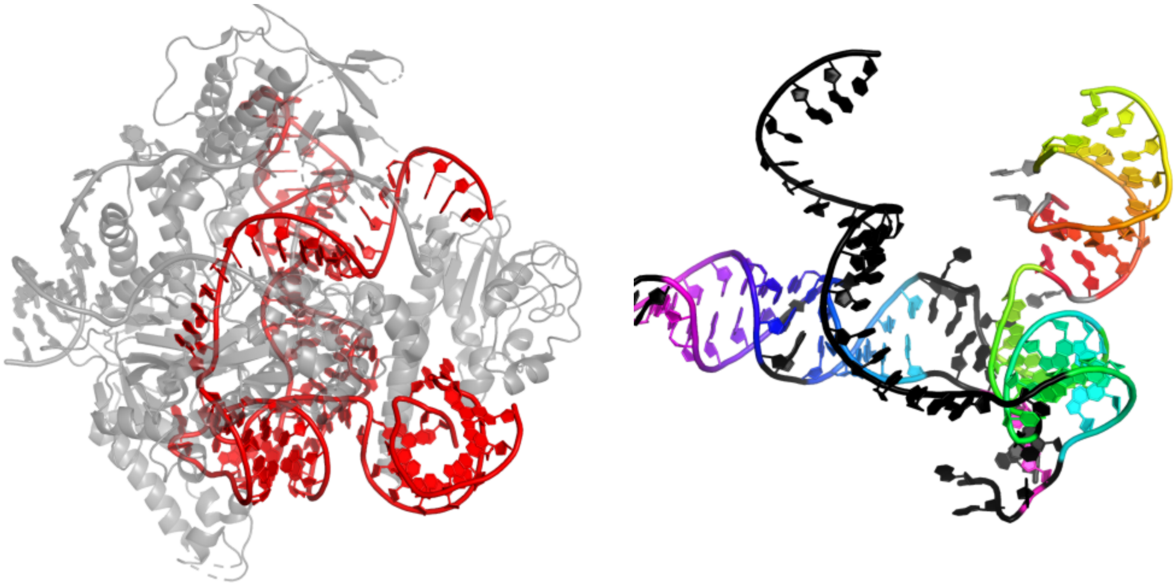

UUCACUGAUAAAGUGGAGAACCGCUUCACCAAAAGCUGUCCCUUAGGGGAUUAGAACUUGAGUGAAGGUGGGCUGCUUGCAUCAGCCUAAUGUCGAGAAGUGCUUUCUUCGGAAAGUAACCCUCGAAACAAAUUCAUUUGAAAGAAUGAAGGAAUGCAACGGAAAUUAGGUGCGCUUGGC

....................((.((((((((...........................)).))))))).)<<<<<.((((((>>>>>.....(((((..((((((((....)))))).)).)))))..(......................).)))))).....................

###### 2.2.4 5wti_B

Crystal structure of the CRISPR-associated protein in complex with crRNA and DNA

**Release date:** 2017-11-01
**Method:** x-ray diffraction
**Resolution:** 2.68 Å
**Chain length:** 123
**Extracted length:** 104

**Total clashes:** 0
**Clashes per residue:** 0.000
**Clashscore:** 7.540

**Description:** RNA (123-MER)
**Organism:** RNA transcription vector pBRDI1

---

**Wu, D., Guan, X., Zhu, Y., Ren, K., Huang, Z. (2017) Structural basis of stringent PAM recognition by CRISPR-C2c1 in complex with sgRNA. Cell Res.**
**DOI:** 10.1038/cr.2017.46

---

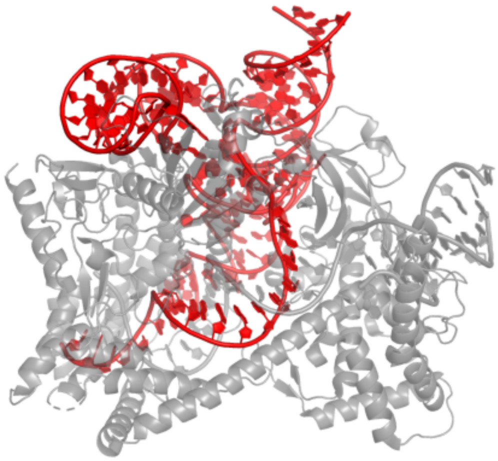

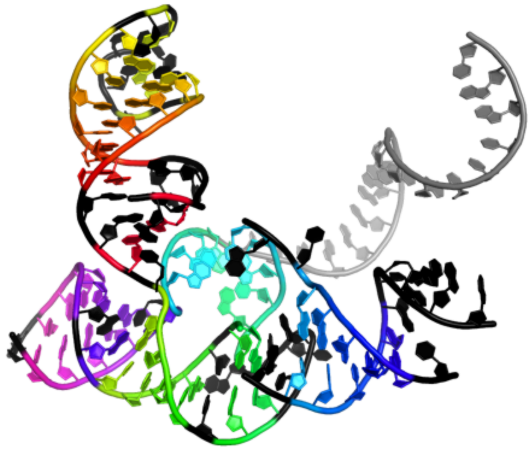

```
GGCGAGGUUCUGUCUUUUGGUCAGGACAACCGUCUAGCUAUAAGUGCUGCAGGGGUGUGAGAAACUCCUAUUGCUGGACGAUGUCUCUUUCGAGGCAUUAGCACCGGGGAGAAGUCAUUUAAU
.(....((((((.(.....).))))))..).((((.((.....<<<<<..(((((.........)))))...)).))))(.(((((......))))).)>>>>>...................
```

#### 2.3 Cas12 guide

### 2.3.1 8bf8_B

ISDra2 TnpB in complex with reRNA

**Release date:** 2023-04-12
**Method:** electron microscopy
**Resolution:** 2.80 Å
**Chain length:** 150
**Extracted length:** 123

**Total clashes:** 49
**Clashes per residue:** 0.327
**Clashscore:** 8.730

**Description:** Deinococcus radiodurans R1 chromosome 1
**Organism:** Deinococcus radiodurans R1 = ATCC 13939 = DSM 20539

---

**Sasnauskas, G., Tamulaitiene, G., Druteika, G., Carabias, A., Silanskas, A., Kazlauskas, D., Venclovas, C., Montoya, G., Karvelis, T., Siksnys, V. (2023) TnpB structure reveals minimal functional core of Cas12 nuclease family. Nature.**
**DOI:** 10.1038/s41586-023-05826-x

---

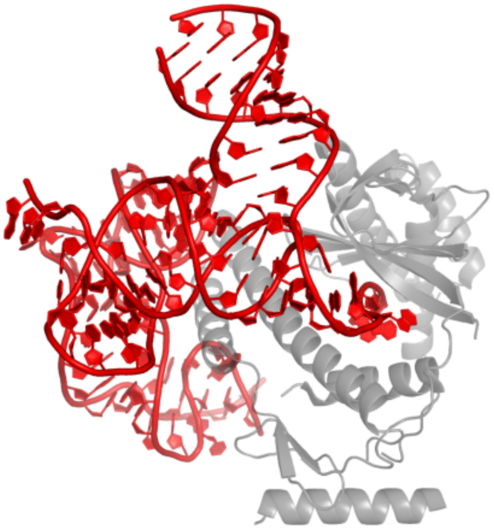

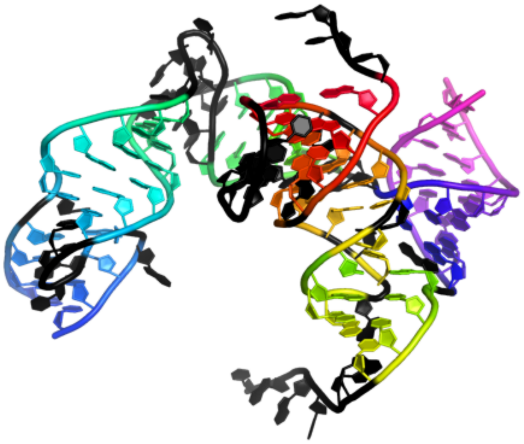

```
CAUUCGGCGUGAAGCGUUGGUGGCUGCGGGAAUCUCAGACACCUUAAACGCUCAUGGAGGCUAUGUCAGACCUGCUUCGGCGGGCAAUGGUCUGCGAAGUGAGAAUCACGCGACUUUAGUCGUGUGAGGUUCAAGAGUCCCUUGGCGCCC
..................((((.(((.(.<<<<).))).)))).......(((((...((((((....(.(((........)))).))))))......)))))..(((((((((....))))))))).>>>>..................
```

### 2.3.2 8j3r_C

Cryo-EM structure of the AsCas12f-HKRA-sgRNAS3-5v7-target DNA

**Release date:** 2023-09-27
**Method:** electron microscopy
**Resolution:** 2.95 Å
**Chain length:** 118
**Extracted length:** 109

**Total clashes:** 18
**Clashes per residue:** 0.153
**Clashscore:** 5.620

**Description:** RNA (118-MER)
**Organism:** Sulfoacidibacillus thermotolerans

---

**Hino, T., Omura, S.N., Nakagawa, R., Togashi, T., Takeda, S.N., Hiramoto, T., Tasaka, S., Hirano, H., Tokuyama, T., Uosaki, H., Ishiguro, S., Kagieva, M., Yamano, H., Ozaki, Y., Motooka, D., Mori, H., Kirita, Y., Kise, Y., Itoh, Y., Matoba, S., Aburatani, H., Yachie, N., Karvelis, T., Siksnys, V., Ohmori, T., Hoshino, A., Nureki, O. (2023) An AsCas12f-based compact genome-editing tool derived by deep mutational scanning and structural analysis. Cell.**
**DOI:** 10.1016/j.cell.2023.08.031

---

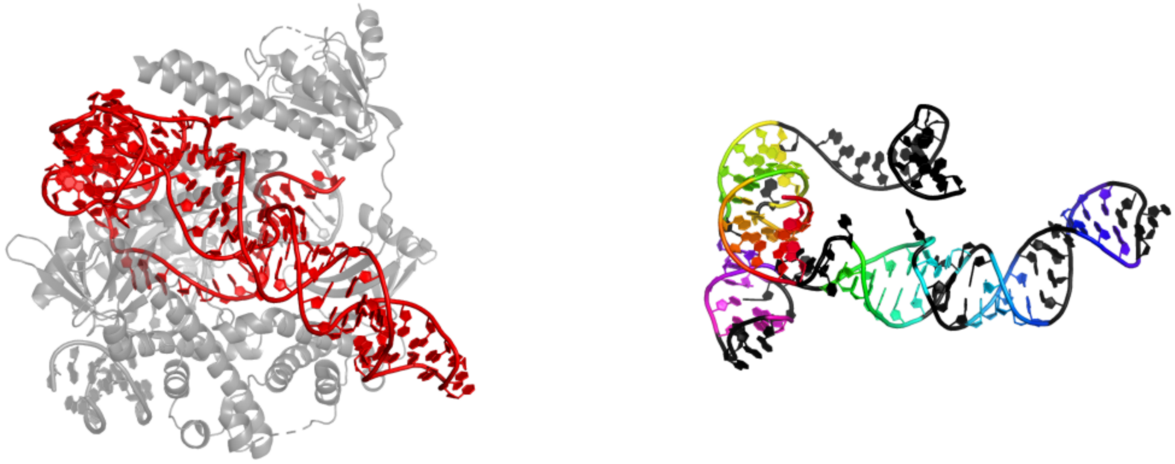

GGAUUCGUCGGUUCAGCGACGAUAAGCCGAGAAGUGCCAAUAAAACUGUUAAGUGGUUUGGUAACGCUCGGUAAGGUCCGAAAGGAGAACCACUGAACGGAAAUUAGGCGCGCUUGGC

....((((((.<<<<<))))))...((((((..(..((((....(((....)))...))))...).))))))..(((((....))...))).>>>>>.....................

##### 2.3.3 6xmf_C

Cryo-EM structure of Cas12g binary complex

**Release date:** 2021-01-13
**Method:** electron microscopy
**Resolution:** 3.10 Å
**Chain length:** 122
**Extracted length:** 119

**Total clashes:** 0
**Clashes per residue:** 0.000
**Clashscore:** 8.600

**Description:** RNA (116-MER)
**Organism:** metagenome

**Li, Z., Zhang, H., Xiao, R., Han, R., Chang, L. (2021) Cryo-EM structure of the RNA-guided ribonuclease Cas12g. Nat.Chem.Biol.**
**DOI:** 10.1038/s41589-020-00721-2

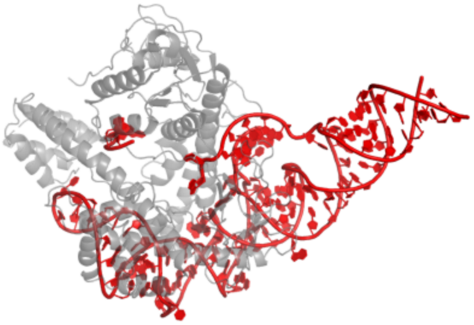

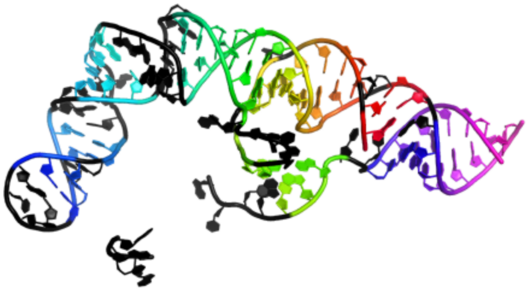

GGGAUGCUUACUUAGUCAUCUGGUUGGCAAACCUCCGCGGACCUUCGGGACCAAUGGAGAGGAACCCAGCCGAGAAGCAUCGAGCCGGUAAAUGUUUACCGGCUCUGACACCAACUGGUGAA

..((.((((.(((.<<<....((((((.....(((...((.((....)).))....)))......))))))))))))).)).((((((((......))))))))..>>>.............

### 2.3.4 8dc2_B

Cryo-EM structure of CasLambda (Cas12l) bound to crRNA and DNA

**Release date:** 2022-12-14
**Method:** electron microscopy
**Resolution:** 2.99 Å
**Chain length:** 52
**Extracted length:** 36

**Total clashes:** 51
**Clashes per residue:** 0.981
**Clashscore:** 11.650

**Description:** RNA (51-MER)
**Organism:** uncultured virus

**Al-Shayeb, B., Skopintsev, P., Soczek, K.M., Stahl, E.C., Li, Z., Groover, E., Smock, D., Eggers, A.R., Pausch, P., Cress, B.F., Huang, C.J., Staskawicz, B., Savage, D.F., Jacobsen, S.E., Banfield, J.F., Doudna, J.A. (2022) Diverse virus-encoded CRISPR-Cas systems include streamlined genome editors. Cell.**
**DOI:** 10.1016/j.cell.2022.10.020

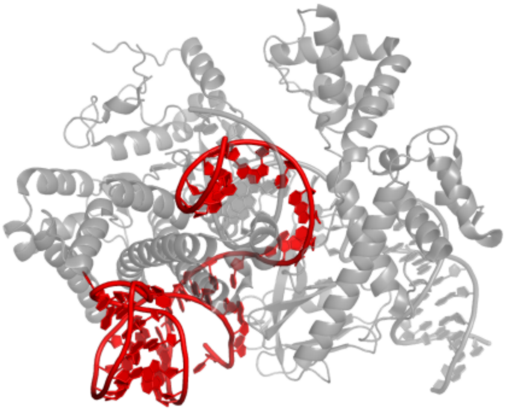

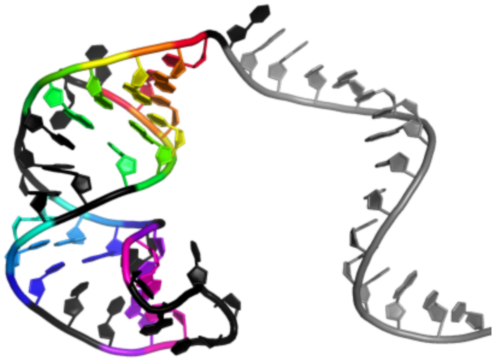

```
AUUGUUGUAACUCUUAUUUUGUAUGGAGUAAACAACUAGCAUCACCUUCACC
...(((((..(((..((.....)).)))...))))).................
```

#### 2.4 Cas13 guide

##### 2.4.1 6dtd_C

High-resolution crystal structure of Cas13b from Prevotella buccae

**Release date:** 2019-02-20
**Method:** x-ray diffraction
**Resolution:** 1.65 Å
**Chain length:** 37
**Extracted length:** 37

**Total clashes:** 0
**Clashes per residue:** 0.000
**Clashscore:** 4.840

**Description:** RNA (37-MER)
**Organism:** Segatella buccae

**Slaymaker, I.M., Mesa, P., Kellner, M.J., Kannan, S., Brignole, E., Koob, J., Feliciano, P.R., Stella, S., Abudayyeh, O.O., Gootenberg, J.S., Strecker, J., Montoya, G., Zhang, F. (2019) High-Resolution Structure of Cas13b and Biochemical Characterization of RNA Targeting and Cleavage. Cell Rep.**
**DOI:** 10.1016/j.celrep.2019.02.094

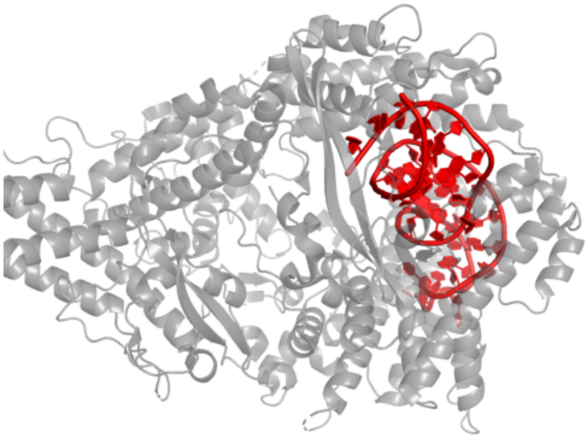

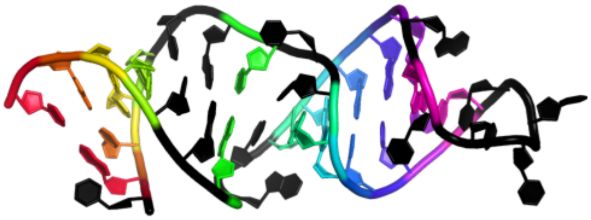

```
UGUUGCAUCUGCCUUCUUUUUGAAAGGUAAAAACAAC
.((((..(..((((((.....).)))))..)..))))
```

##### 2.4.2 8wcs_G

Cryo-EM structure of Cas13h1-crRNA binary complex

**Release date:** 2024-05-22
**Method:** electron microscopy
**Resolution:** 3.10 Å
**Chain length:** 66
**Extracted length:** 66

**Total clashes:** 0
**Clashes per residue:** 0.000
**Clashscore:** 4.750

**Description:** 66-nt crRNA
**Organism:** synthetic construct

**Chen, F., Zhang, C., Xue, J., Wang, F., Li, Z. (2024) Molecular mechanism for target RNA recognition and cleavage of Cas13h. Nucleic Acids Res.**
**DOI:** 10.1093/nar/gkae324

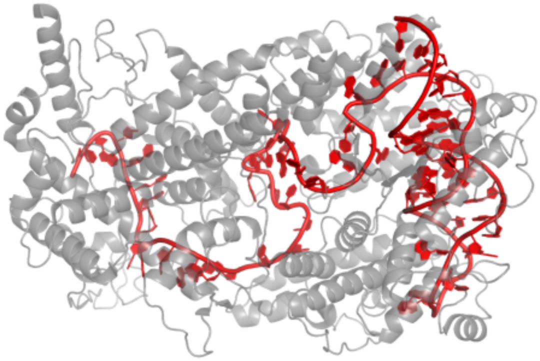

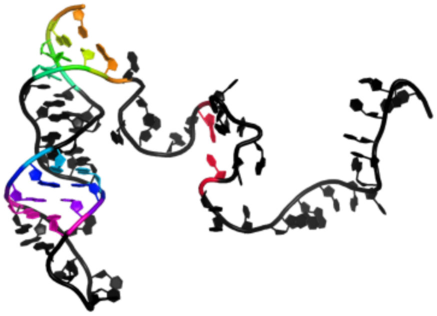

```
UGCUUCACGUAGGCCUUGGAGCCGUACAUGGUUGUAACAAGCCUAAGUUUGAAAGGUAAAAACAAC
................(......)......((((......((((.........)))).....))))
```

##### 2.4.3 6aay_B

the Cas13b binary complex

**Release date:** 2019-03-13
**Method:** x-ray diffraction
**Resolution:** 2.79 Å
**Chain length:** 59
**Extracted length:** 59

**Total clashes:** 49
**Clashes per residue:** 0.831
**Clashscore:** 26.170

**Description:** RNA (52-MER)
**Organism:** Bergeyella zoohelcum

---

**Zhang, B., Ye, W.W., Ye, Y.M., Zhou, H., Saeed, A.F.U.H., Chen, J., Lin, J.Y., Perculija, V., Chen, Q., Chen, C.J., Chang, M.X., Choudhary, M.I., Ouyang, S.Y. (2018) Structural insights into Cas13b-guided CRISPR RNA maturation and recognition. Cell Res.**
**DOI:** 10.1038/s41422-018-0109-4

---

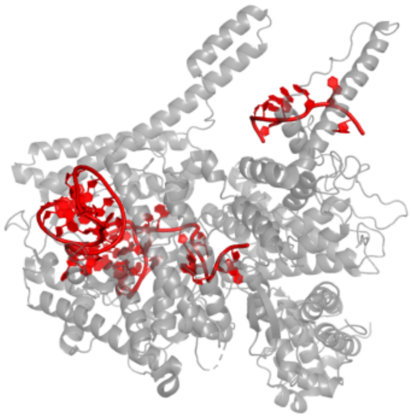

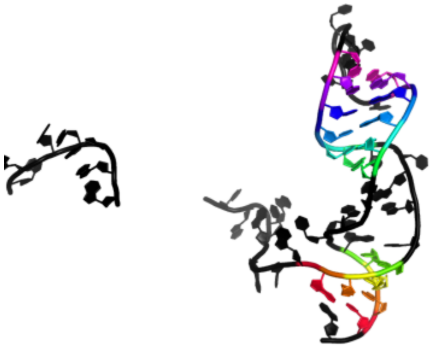

```
AAAAAGGGUUUAAAAAAUGAAAGUUGGAACUGCUCUCAUUUUGGAGGGUAAUCACAACA
......................((((.....((((((......)))))).....)))).
```

##### 2.4.4 8ewg_B

Cryo-EM structure of a riboendonclease

**Release date:** 2023-08-30
**Method:** electron microscopy
**Resolution:** 2.90 Å
**Chain length:** 60
**Extracted length:** 57

**Total clashes:** 27
**Clashes per residue:** 0.450
**Clashscore:** 5.360

**Description:** RNA (56-MER)
**Organism:** Thermoclostridium caenicola

---

**Wang, F., Zhang, C., Xu, H., Zeng, W., Ma, L., Li, Z. (2023) Structural Basis for the Ribonuclease Activity of a Thermostable CRISPR-Cas13a from Thermoclostridium caenicola. J.Mol.Biol.**
**DOI:** 10.1016/j.jmb.2023.168197

---

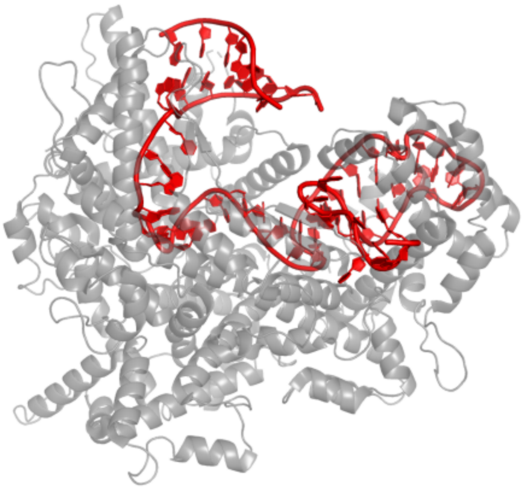

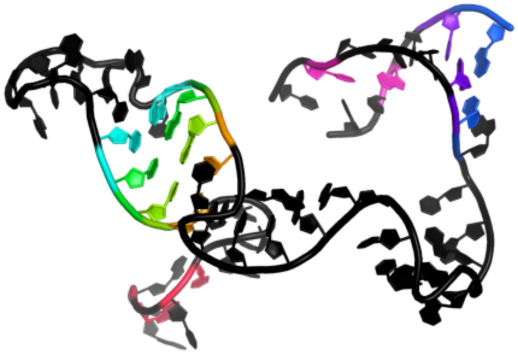

```
UAGGGUCACAACUCCCAUGUAGGCGGAGACUGCAACCCGAAGGUGUGACUUCCAUGCCAA
....(.)....((((.........)))).................((...(....).)).
```

#### 2.5 OMEGA effector guide

##### 2.5.1 8gkh_W

Structure of the Spizellomyces punctatus Fanzor (SpuFz) in complex with omega RNA and target DNA

**Release date:** 2023-07-05
**Method:** electron microscopy
**Resolution:** 2.70 Å
**Chain length:** 78
**Extracted length:** 63

**Total clashes:** 0
**Clashes per residue:** 0.000
**Clashscore:** 5.490

**Description:** RNA (78-MER)
**Organism:** Spizellomyces punctatus

**Saito, M., Xu, P., Faure, G., Maguire, S., Kannan, S., Altae-Tran, H., Vo, S., Desimone, A., Macrae, R.K., Zhang, F. (2023) Fanzor is a eukaryotic programmable RNA-guided endonuclease. Nature.**
**DOI:** 10.1038/s41586-023-06356-2

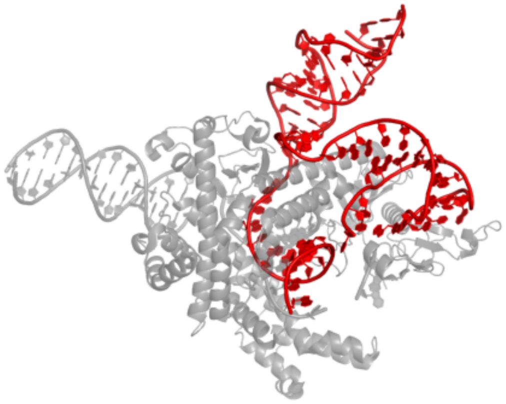

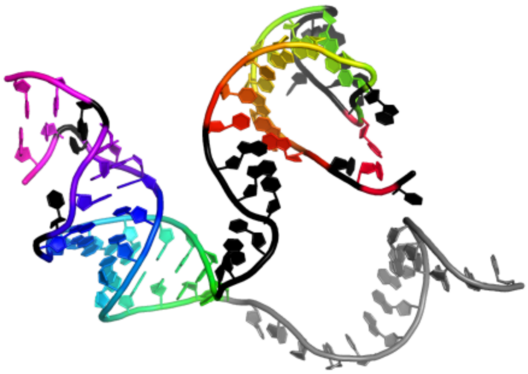

UCCGAGCCGGUAGUCGCGCGGUUCAAUCCCUGGUGCGGGUGCUAGUGCACCGGCUCCGCACUAUCUAUAGGUUAUGAA
.<.(((((((.....>).))))))......(((((((((.(((.((..)).))))))))))))...............

#### 2.6 Fanzor guide

### 2.6.1 9cf2_W

Parasitella parasitica Fanzor (PpFz) State 3

**Release date:** 2024-09-11
**Method:** electron microscopy
**Resolution:** 3.15 Å
**Chain length:** 61
**Extracted length:** 42

**Total clashes:** 0
**Clashes per residue:** 0.000
**Clashscore:** 3.490

**Description:** Parasitella parasitica Fanzor 1 omegaRNA
**Organism:** Parasitella parasitica

**Xu, P., Saito, M., Faure, G., Maguire, S., Chau-Duy-Tam Vo, S., Wilkinson, M.E., Kuang, H., Wang, B., Rice, W.J., Macrae, R.K., Zhang, F. (2024) Structural insights into the diversity and DNA cleavage mechanism of Fanzor. Cell.**
**DOI:** 10.1016/j.cell.2024.07.050

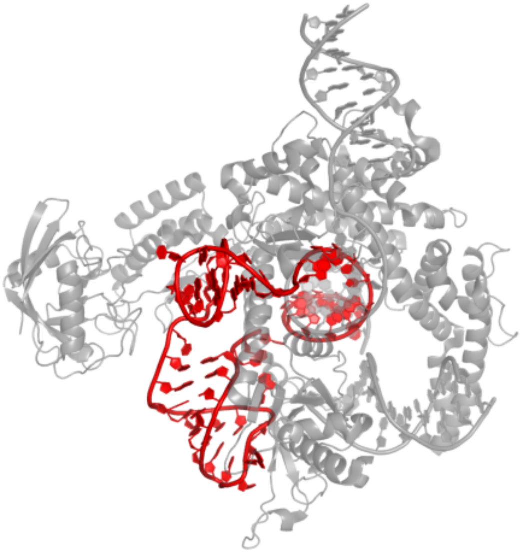

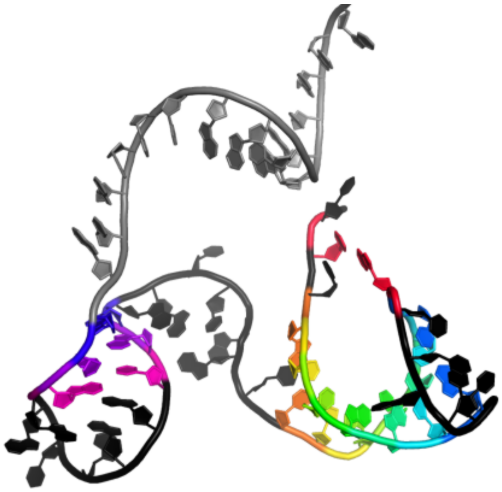

UUAUCCACCAAAGUUAUCGCUUUGGUCAAUUAAUGCAGGUAAGCAACAUCCAGCAAACAGA
....<.(((((((.....>))))))).......(((......)))................

### 3 IRES

#### 3.1 IAPV IRES

### 3.1.1 6p5i_1

Structure of a mammalian 80S ribosome in complex with the Israeli Acute Paralysis Virus IRES (Class 1)

**Release date:** 2019-09-18
**Method:** electron microscopy
**Resolution:** 3.10 Å
**Chain length:** 253
**Extracted length:** 205

**Total clashes:** 0
**Clashes per residue:** 0.000
**Clashscore:** 2.800

**Description:** IAPV-IRES
**Organism:** Israeli acute paralysis virus

---

**Acosta-Reyes, F., Neupane, R., Frank, J., Fernandez, I.S. (2019) The Israeli acute paralysis virus IRES captures host ribosomes by mimicking a ribosomal state with hybrid tRNAs. Embo J.**
**DOI:** 10.15252/embj.2019102226

---

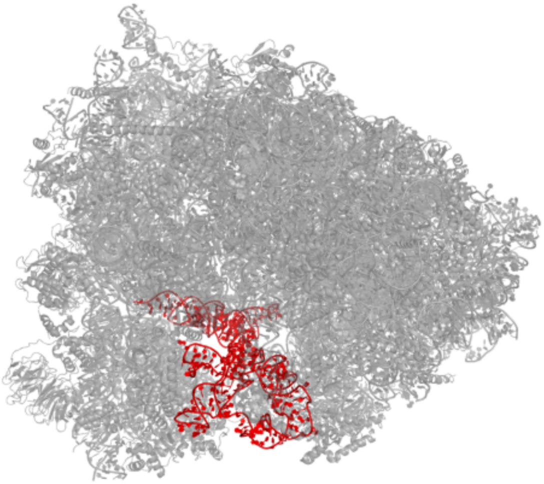

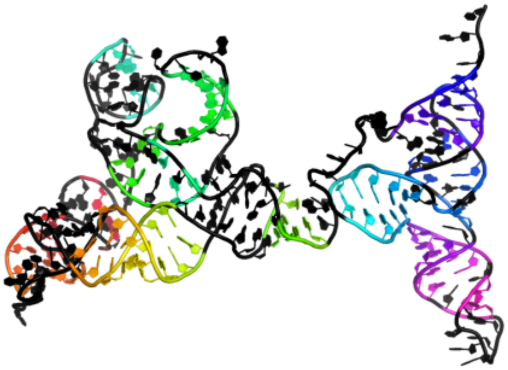

GAGCGGUUUCUGGAAUACUAUAUGUAAGUAUAGUGUUCUGGAGGCAUCAUUCUAUGGUUACCCAUCAUUAGAGGAAAUUUCCAAUAAACUCUGGUGUAAGGCUUAGAGUGAUGGUCGAGGUGCCCUAUUUAGGGUGAGGAGCCUCGGUGGCAGCCCCACCAAAUCCUCUAUUGGAUAGGAACAGCUGUACUGGGCAGUUACAGCAGUCGUAUGGUAACACAUGCGGCGUUCCGAAAUACCAUGCCUGGCGAUG

.............................................(...(((((...........((((((((......<<<......))))))))........)))))...)........((.((((....))))...<<.))......((.....))........>.>.....>>>..(((((.(.((.(..<<<<<..).).)).(((((.((......)).))))))))))).........>>>>>........

### 3.1.2 6p5n_1

Structure of a mammalian 80S ribosome in complex with a single translocated Israeli Acute Paralysis Virus IRES and eRF1

**Release date:** 2019-09-25
**Method:** electron microscopy
**Resolution:** 3.20 Å
**Chain length:** 251
**Extracted length:** 207

**Total clashes:** 0
**Clashes per residue:** 0.000
**Clashscore:** 2.150

**Description:** IAPV-IRES
**Organism:** Israeli acute paralysis virus

---

**Acosta-Reyes, F., Neupane, R., Frank, J., Fernandez, I.S. (2019) The Israeli acute paralysis virus IRES captures host ribosomes by mimicking a ribosomal state with hybrid tRNAs. Embo J.**
**DOI:** 10.15252/embj.2019102226

---

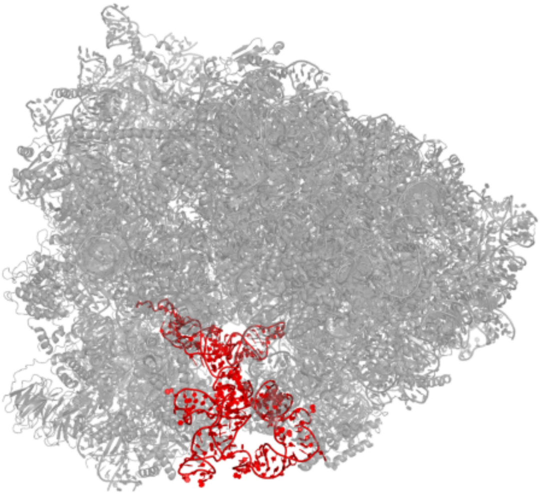
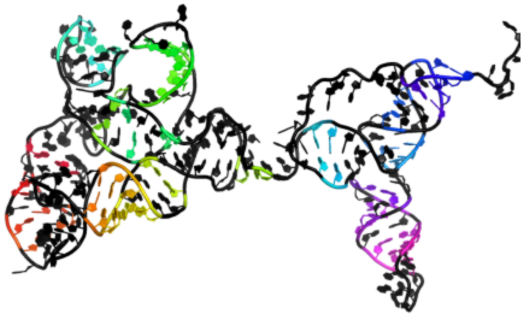

GAGCGGUUUCUGGAAUACUAUAUGUAAGUAUAGUGUUCUGGAGGCAUCAUUCUAUGGUUACCCAUCAUUAGAGGAAAUUUCCAAUAAACUCUGGUGUAAGGCUUAGAGUGAUGGUCGAGGUGCCCUAUUUAGGGUGAGGAGCCUCGGUGGCAGCCCCACCAAAUCCUCUAUUGGAUAGGAACAGCUGUACUGGGCAGUUACAGCAGUCGUAUGGUAACACAUGCGGCGUUCCGAAAUACCAUGCCUUGAUA

...............................................(..(.((...........(((((..(........<......)..))))).......).).).).........<..((((....))))...(.(.>.....(((.....))).....)..)......>....((....((..(..<<<<..)...))..(.((.(((......))).)).)..))...........>>>>.....

#### 3.2 CrPV 5'UTR IRES

### 3.2.1 6w2t_A

Structure of the Cricket Paralysis Virus 5-UTR IRES (CrPV 5-UTR-IRES) bound to the small ribosomal subunit in the closed state (Class 2)

**Release date:** 2020-04-22
**Method:** electron microscopy
**Resolution:** 3.36 Å
**Chain length:** 377
**Extracted length:** 339

**Total clashes:** 0
**Clashes per residue:** 0.000
**Clashscore:** 5.380

**Description:** CrPV 5'-UTR IRES
**Organism:** Cricket paralysis virus

---

**Neupane, R., Pisareva, V.P., Rodriguez, C.F., Pisarev, A.V., Fernandez, I.S. (2020) A complex IRES at the 5'-UTR of a viral mRNA assembles a functional 48S complex via an uAUG intermediate. Elife.**
**DOI:** 10.7554/eLife.54575

---

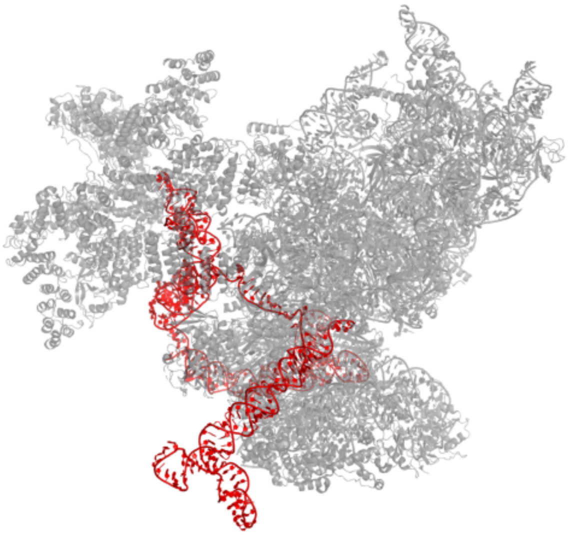

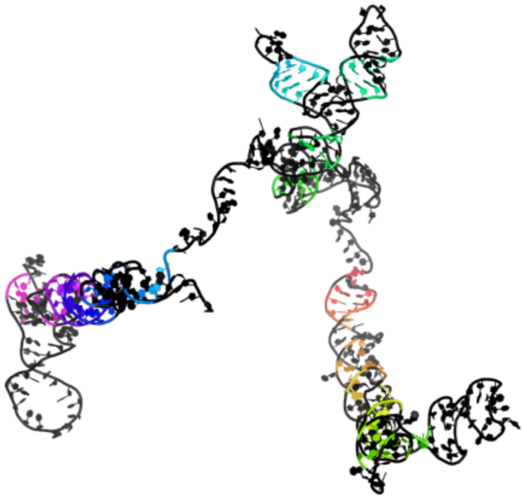

UUCGAUACCAAGAGCUGGUGUCUGAGGGAUUACGACUCGCGUGGUCGGACAGCACAUGUUAGUCAUGUAUUAUGGUGUUUCCCAACACGUAUCUUUGCUCUAGUAUUGAACUUUAGUGAUGAGUGUACUUGGCAUGUACACCUGAUUAGUUAGCCCUUGUGGAUACGCCACUGCCGCUCAGGUACAGUAGGACAUACCCCAAAUAUUGGGUUUUUUUUUUAGAGAAGAAAAUUAGUGGCGAAACGUACGGUUUAUGUCAAAGGAGAAGGUGAUUUUAAUUGCACCUGAAGCAACCCUAGGUCUUCCCUUUGAUGUGAGCUAAUAAGGAGUAACCGAUUCUUACAAUGUGAUCAUGUCUUUUCAACAAACAAAGGAUC

(.((((........((.((...(...(((.....(..(.....).)......(.......................)..)))......)...))..).)......)))).).............(((.........)))........................(.(......((..(........)..))......((((.....))))...).)....................((.....(....((((((((((...(.............................((........)...)...)..))))))))))).....)......)).........................................

#### 3.3 TSV IRES

##### 3.3.1 8evp_EC

Hypopseudouridylated yeast 80S bound with Taura syndrome virus (TSV) internal ribosome entry site (IRES), Structure I

**Release date:** 2023-09-06
**Method:** electron microscopy
**Resolution:** 2.38 Å
**Chain length:** 202
**Extracted length:** 196

**Total clashes:** 0
**Clashes per residue:** 0.000
**Clashscore:** 5.910

**Description:** Internal ribosome entry site
**Organism:** Taura syndrome virus

---

**Zhao, Y., Rai, J., Li, H. (2023) Regulation of translation by ribosomal RNA pseudouridylation. Sci Adv.**
**DOI:** 10.1126/sciadv.adg8190

---

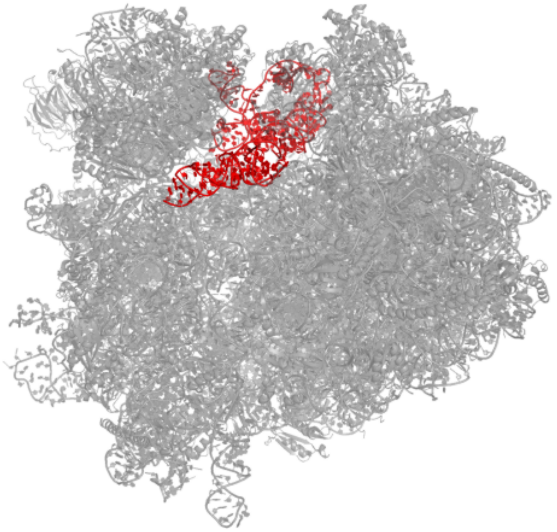

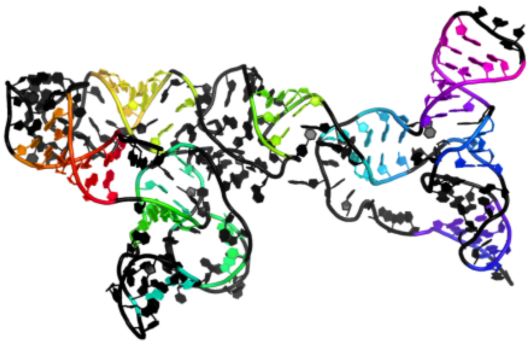

AAACUCCAUGUAUUGGUUACCCAUCUGCAUCGAAAACUCUCCGAACACUAGGUGCAGUAAGGCUUUCAUGGAGUGGUUUGCUAUUUAGCGUACGUGUACCAUAGGCAGCCCCAAAAACACGUGUGAGGAGAAAGUCCCAGUCACUUUGGGCAAAGUAGACAGCCGCGCUUGCGUGGUGGGACUUAAUUAAUGCCUGCUAACC

...((((((...............((((.(........<<<..........).))))..........)))))).(((..((......)).....<.<)))...(.......)....>.>........>>>...((((..(((.....<<<<......))).((((((....)))))).))).)........>>>>.......

#### 3.4 PSIV IGR IRES

### 3.4.1 4v83_CV

Crystal structure of a complex containing domain 3 from the PSIV IGR IRES RNA bound to the 70S ribosome.

**Release date:** 2014-07-09
**Method:** x-ray diffraction
**Resolution:** 3.50 Å
**Chain length:** 35
**Extracted length:** 35

**Total clashes:** 0
**Clashes per residue:** 0.000
**Clashscore:** 20.140

**Description:** domain 3 of PSIC IGR IRES RNA
**Organism:** Thermus thermophilus HB27

**Zhu, J., Korostelev, A., Costantino, D.A., Donohue, J.P., Noller, H.F., Kieft, J.S. (2011) Crystal structures of complexes containing domains from two viral internal ribosome entry site (IRES) RNAs bound to the 70S ribosome. Proc.Natl.Acad.Sci.USA.**
**DOI:** 10.1073/pnas.1018582108

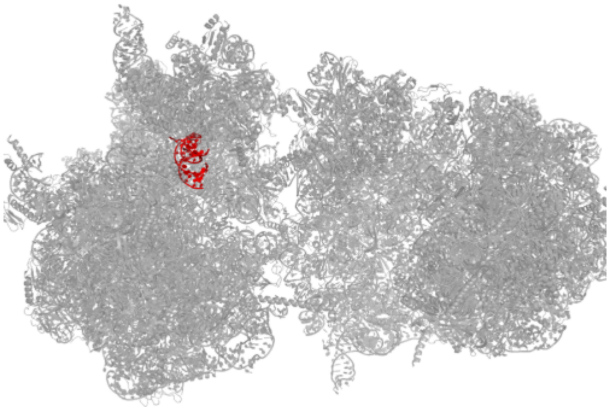

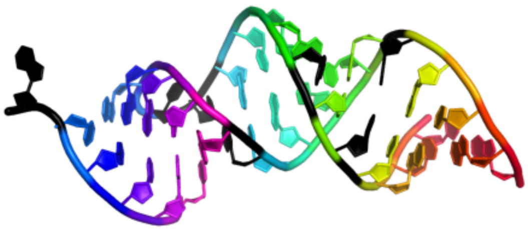

```
UCGCUCAAACAUUAAGUGGUGUUGUGCGACACUUA
((((.(.((((..<<<<<.))))).))))>>>>>.
```

### 4 Ribozymes

#### 4.1 Synthetic ligase ribozyme

##### 4.1.1 3hhn_E

Crystal structure of class I ligase ribozyme self-ligation product, in complex with U1A RBD

**Release date:** 2009-11-24
**Method:** x-ray diffraction
**Resolution:** 2.99 Å
**Chain length:** 137
**Extracted length:** 137

**Total clashes:** 58
**Clashes per residue:** 0.423
**Clashscore:** 12.640

**Description:** Class I ligase ribozyme, self-ligation product
**Organism:** nan

---

**Shechner, D.M., Grant, R.A., Bagby, S.C., Koldobskaya, Y., Piccirilli, J.A., Bartel, D.P. (2009)**
**Crystal structure of the catalytic core of an RNA-polymerase ribozyme. Science.**
**DOI:** 10.1126/science.1174676

---

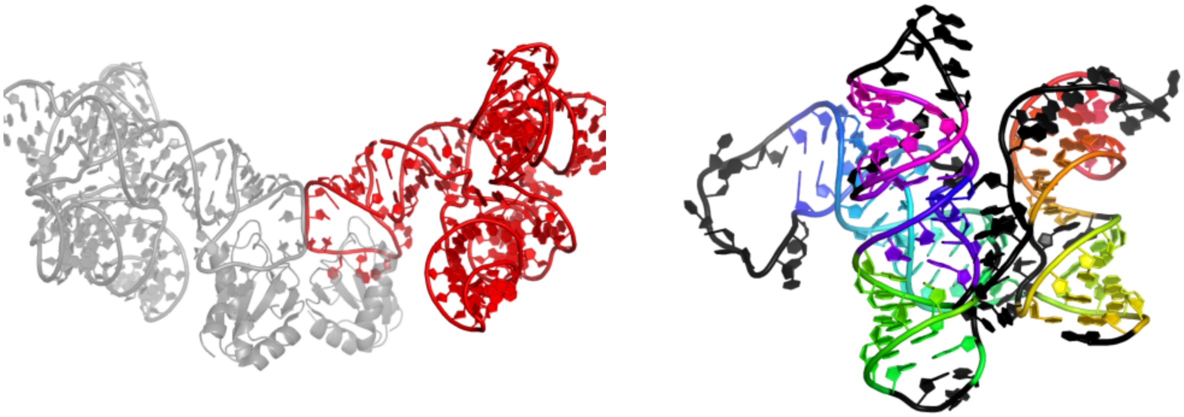

```
UCCAGUAGGAACACUAUACUACUGGAUAAUCAAAGACAAAUCUGCCCGAAGGGCUUGAGAACAUACCCAUUGCACUCCGGGUAUGCAGAGGUGGCAGCCUCCGGUGGGUUAAAACCCAACGUUCUCAACAAUAGUGA
((((((((...<<<<<<.))))))))...............<<<<<(...).(.((((((((((((((..........)))))))..((((.>>>>>))))((.((((......)))).)))))))))).>>>>>>.
```

##### 4.1.2 3ivk_C

Crystal Structure of the Catalytic Core of an RNA Polymerase Ribozyme Complexed with an Antigen Binding Antibody Fragment

**Release date:** 2010-03-02
**Method:** x-ray diffraction
**Resolution:** 3.10 Å
**Chain length:** 128
**Extracted length:** 128

**Total clashes:** 0
**Clashes per residue:** 0.000
**Clashscore:** 31.380

**Description:** class I ligase product
**Organism:** nan

**Shechner, D.M., Grant, R.A., Bagby, S.C., Koldobskaya, Y., Piccirilli, J.A., Bartel, D.P. (2009)**
**Crystal structure of the catalytic core of an RNA-polymerase ribozyme. Science.**
**DOI:** 10.1126/science.1174676

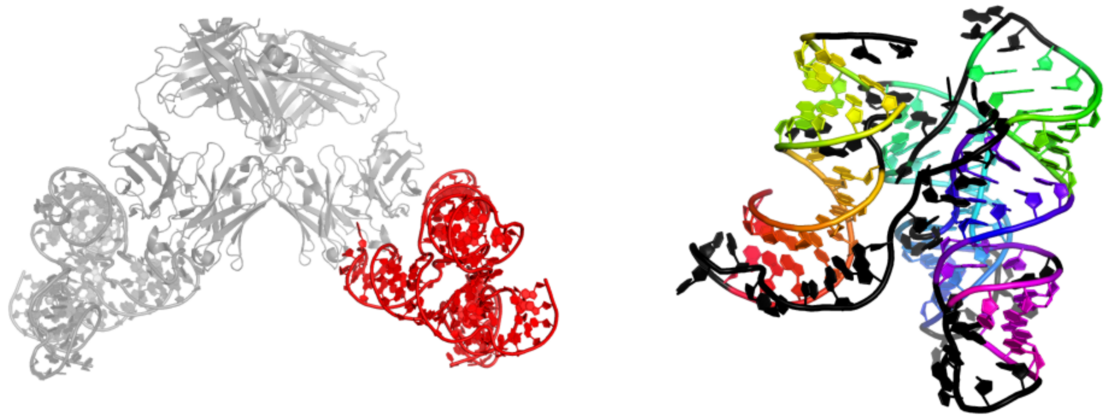

UCCAGUAGGAACACUAUACUACUGGAUAAUCAAAGACAAAUCUGCCCGAAGGGCUUGAGAACAUCGAAACACGAUGCAGAGGUGGCAGCCUCCGGUGGGUUAAAACCCAACGUUCUCAACAAUAGUGA

((((((((....<<<<<.))))))))...............<<<<<(...)...((((((((((((.....)))))..((((.>>>>>))))((.((((......)))).)))))))))..>>>>>..

### 4.1.3 8t2p_B

5TU-t1 - heterodimeric triplet polymerase ribozyme

**Release date:** 2024-01-24
**Method:** electron microscopy
**Resolution:** 5.00 Å
**Chain length:** 152
**Extracted length:** 152

**Total clashes:** 0
**Clashes per residue:** 0.000
**Clashscore:** 0.000

**Description:** RNA (152-MER)
**Organism:** synthetic construct

---

**McRae, E.K.S., Wan, C.J.K., Kristoffersen, E.L., Hansen, K., Gianni, E., Gallego, I., Curran, J.F., Attwater, J., Holliger, P., Andersen, E.S. (2024) Cryo-EM structure and functional landscape of an RNA polymerase ribozyme. Proc.Natl.Acad.Sci.USA.**
**DOI:** 10.1073/pnas.2313332121

---

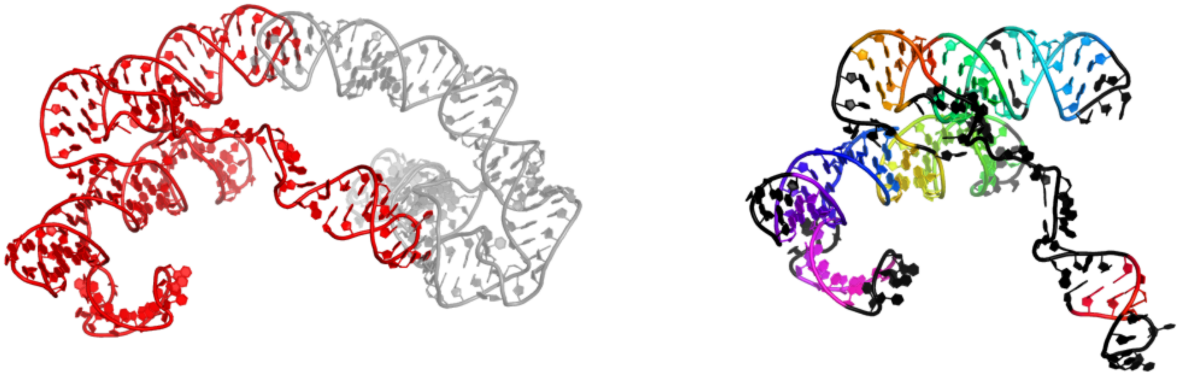

GGAUCUUCUCGAUCUAACAAAAAAGACAAAUCUGCCACAAAGCUUGAGAGCAUCUUCGGAUGCAGAGGCGGCAGCCUUCGGUGGCGCGAUAGCGCCAACGUUCUCAACUAUGACACGCAAAACGCGUGCUCCGUUGAAUGGAGUUUAUCAUG

.(((.......))).................<<<<<.....(.((((((((((.(...))))).((((.>>>>>))))((.((((((....)))))).)).)))))))((((((((((.....)))))(((((.....)))))....)))))

#### 4.2 VS ribozyme

### 4.2.1 4r4v_A

Crystal structure of the VS ribozyme - G638A mutant

**Release date:** 2015-09-30
**Method:** x-ray diffraction
**Resolution:** 3.07 Å
**Chain length:** 186
**Extracted length:** 186

**Total clashes:** 38
**Clashes per residue:** 0.204
**Clashscore:** 3.260

**Description:** VS ribozyme RNA
**Organism:** Neurospora

**Suslov, N.B., DasGupta, S., Huang, H., Fuller, J.R., Lilley, D.M., Rice, P.A., Piccirilli, J.A. (2015) Crystal structure of the Varkud satellite ribozyme. Nat.Chem.Biol.**
**DOI:** 10.1038/nchembio.1929

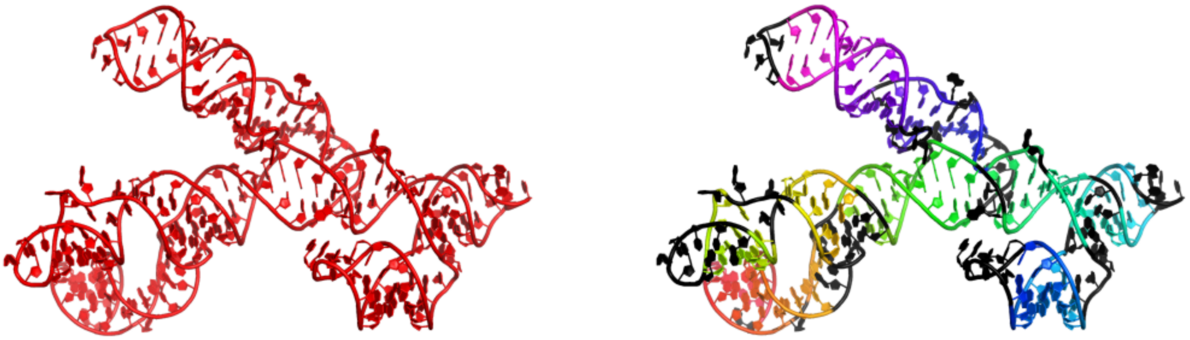

GGCGCUGUGUCGCAAUCUGCGAAGGGCGUCGUCGGCCCAAGCGGUAGUAAGCAGGGAACUCACCUCCAAUGAAACACAUUGUCGUAGCAGUUGACUACUGUUAUGUGAUUGGUAGAGGCUAAGUGACGGUAUUGGCGUAAGCCAAUACCGCAGCACAGCACAAGCCCGCUUGCGAGAUUACAGCGC

.((((((((((.....((((...((((.......))))..))))..((((((.(((..((((((..(((((.....)))))..(((((((.......)))))))......))).)))(((..(((.((((((((((....))))))))))...)))))).....)))))))))..))).)))))))

#### 4.3 Self-alkylating ribozyme

##### 4.3.1 6xjq_A

Crystal structure of a self-alkylating ribozyme - alkylated form with biotinylated epoxide substrate

**Release date:** 2022-01-19
**Method:** x-ray diffraction
**Resolution:** 1.71 Å
**Chain length:** 58
**Extracted length:** 58

**Total clashes:** 10
**Clashes per residue:** 0.172
**Clashscore:** 2.170

**Description:** Self-alkylating ribozyme (58-MER)
**Organism:** Aeropyrum pernix

**Krochmal, D., Shao, Y., Li, N.S., DasGupta, S., Shelke, S.A., Koirala, D., Piccirilli, J.A. (2022) Structural basis for substrate binding and catalysis by a self-alkylating ribozyme. Nat.Chem.Biol.**
**DOI:** 10.1038/s41589-021-00950-z

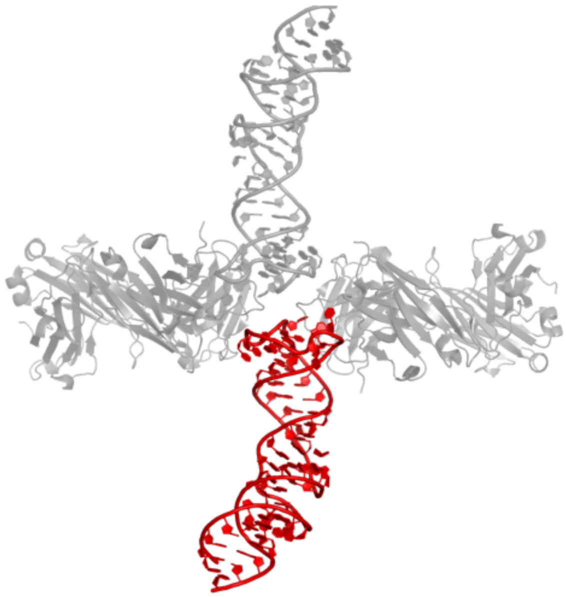

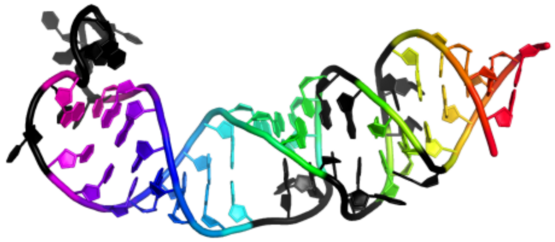

```
GGCCGCUCCAGAAGAGGGCCCCCUUGCCCGUUAUCGGGGGCUAGGCUCGAUGUCGGCC
(((((...((...((((((((((..(........))))))))...)))..)).)))))
```

#### 4.4 Diels-Alder ribozyme

##### 4.4.1 1ykv_D

Crystal structure of the Diels-Alder ribozyme complexed with the product of the reaction between N-pentylmaleimide and covalently attached 9-hydroxymethylanthracene

**Release date:** 2005-02-22
**Method:** x-ray diffraction
**Resolution:** 3.30 Å
**Chain length:** 38
**Extracted length:** 31

**Total clashes:** 26
**Clashes per residue:** 0.684
**Clashscore:** 12.520

**Description:** Diels-Alder ribozyme
**Organism:** nan

---

**Serganov, A., Keiper, S., Malinina, L., Tereshko, V., Skripkin, E., Hobartner, C., Polonskaia, A., Phan, A.T., Wombacher, R., Micura, R., Dauter, Z., Jaschke, A., Patel, D.J. (2005) Structural basis for Diels-Alder ribozyme-catalyzed carbon-carbon bond formation. Nat.Struct.Mol.Biol.**
**DOI:** 10.1038/nsmb906

---

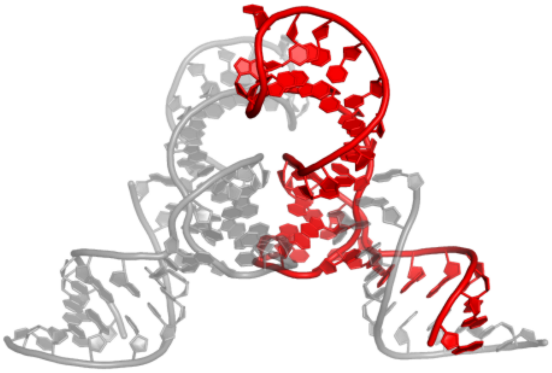

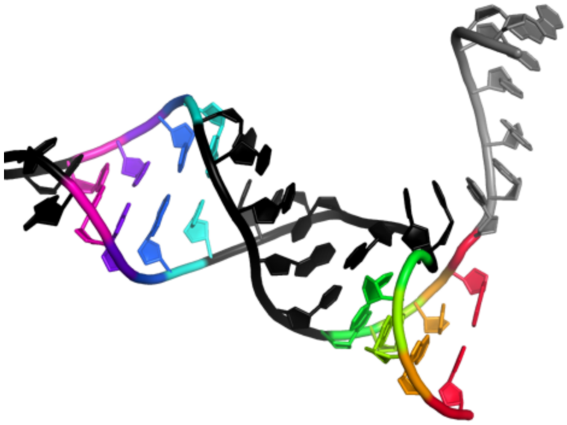

GGGCGAGGCCGUGCCAGCUCUUCGGAGCAAUACUCGGC
.......((((.....((((....))))......))))

##### 4.4.2 1yls_D

Crystal structure of selenium-modified Diels-Alder ribozyme complexed with the product of the reaction between N-pentylmaleimide and covalently attached 9-hydroxymethylanthracene

**Release date:** 2005-02-22
**Method:** x-ray diffraction
**Resolution:** 3.00 Å
**Chain length:** 38
**Extracted length:** 31

**Total clashes:** 43
**Clashes per residue:** 1.132
**Clashscore:** 14.760

**Description:** RNA Diels-Alder ribozyme
**Organism:** nan

**Serganov, A., Keiper, S., Malinina, L., Tereshko, V., Skripkin, E., Hobartner, C., Polonskaia, A., Phan, A.T., Wombacher, R., Micura, R., Dauter, Z., Jaschke, A., Patel, D.J. (2005) Structural basis for Diels-Alder ribozyme-catalyzed carbon-carbon bond formation. Nat.Struct.Mol.Biol.**
**DOI:** 10.1038/nsmb906

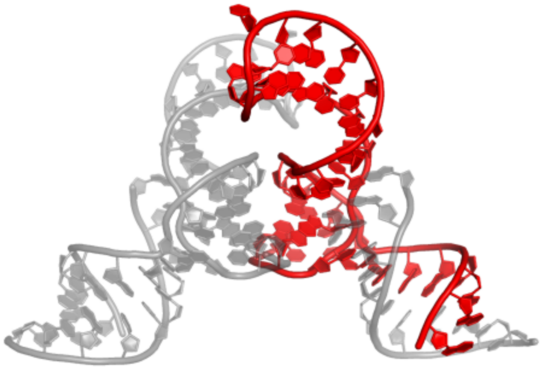

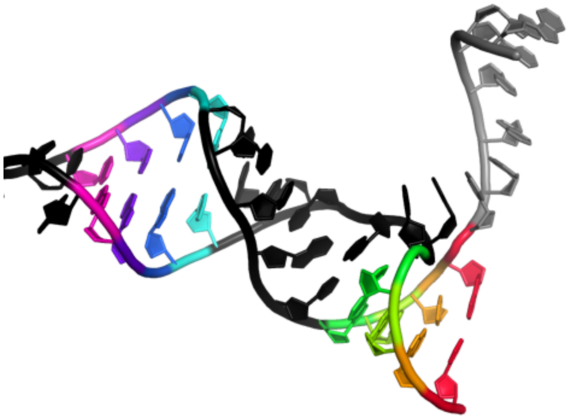

GGGCGAGGCCGUGCCGGCUCUUCGGAGCAAUACUCGGC
.......((((.....((((....)))).......))))

#### 4.5 Methyltransferase ribozyme

##### 4.5.1 7dlz_Y

Crystal Structure of Methyltransferase Ribozyme

**Release date:** 2021-10-27
**Method:** x-ray diffraction
**Resolution:** 3.00 Å
**Chain length:** 45
**Extracted length:** 45

**Total clashes:** 0
**Clashes per residue:** 0.000
**Clashscore:** 8.240

**Description:** RNA (45-MER)
**Organism:** synthetic construct

**Jiang, H.Y., Gao, Y.Q., Zhang, L., Chen, D.R., Gan, J.H., Murchie, A.I.H. (2021) The identification and characterization of a selected SAM-dependent methyltransferase ribozyme that is present in natural sequences. Nat Catal.**
**DOI:** 10.1038/s41929-021-00685-z

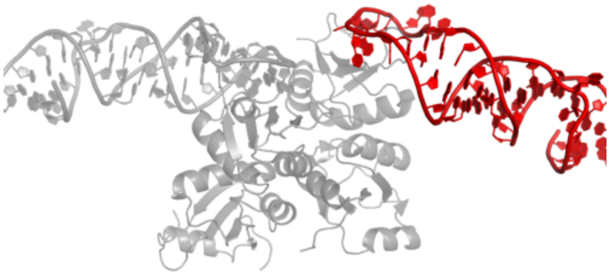

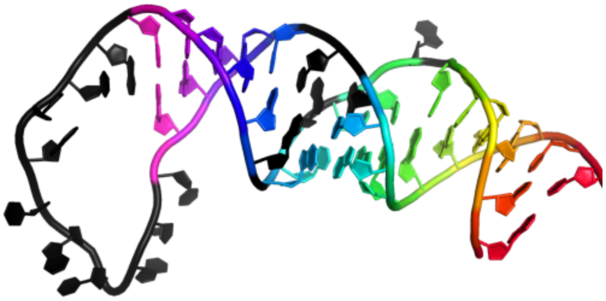

```
GGACCUACUACGAGCGCCAUUGCACUCCGGCGCCACGGGGGGUCC
((((((.((.((.(((((..........)))))..))))))))))
```

### 4.5.2 7v9e_A

Crystal structure of a methyl transferase ribozyme

**Release date:** 2022-03-23
**Method:** x-ray diffraction
**Resolution:** 2.30 Å
**Chain length:** 68
**Extracted length:** 68

**Total clashes:** 4
**Clashes per residue:** 0.059
**Clashscore:** 1.360

**Description:** RNA (68-MER)
**Organism:** Homo sapiens

**Deng, J., Wilson, T.J., Wang, J., Peng, X., Li, M., Lin, X., Liao, W., Lilley, D.M.J., Huang, L. (2022) Structure and mechanism of a methyltransferase ribozyme. Nat.Chem.Biol.**
**DOI:** 10.1038/s41589-022-00982-z

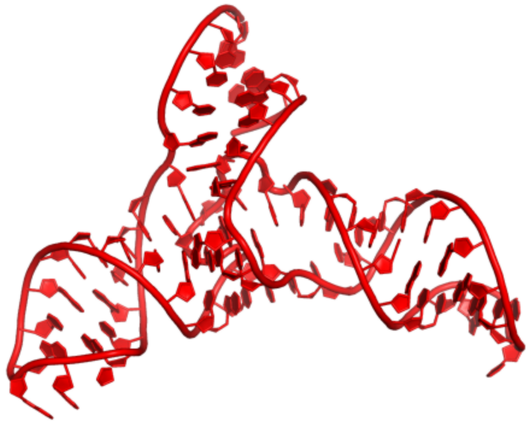

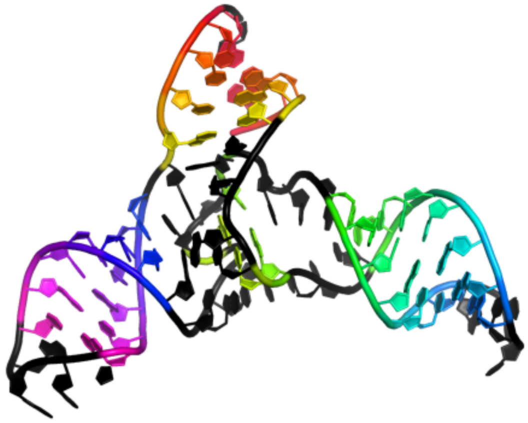

CGGGCUGACCGACCCCCCGAGUUCGCUCGGGGACAACUAGACAUACAGUAUGAAAAUACUGAGCCCGC
(((((....((...(((((((....)))))))....)..).....((((((....)))))).))))).

### 5 Repeats

#### 5.1 r(CUG)

##### 5.1.1 4pcj_A

Modifications to toxic CUG RNAs induce structural stability and rescue mis-splicing in Myotonic Dystrophy

**Release date:** 2014-10-29
**Method:** x-ray diffraction
**Resolution:** 1.90 Å
**Chain length:** 35
**Extracted length:** 35

**Total clashes:** 0
**Clashes per residue:** 0.000
**Clashscore:** 0.000

**Description:** trCUG-3('5)
**Organism:** synthetic construct

---

**deLorimier, E., Coonrod, L.A., Copperman, J., Taber, A., Reister, E.E., Sharma, K., Todd, P.K., Guenza, M.G., Berglund, J.A. (2014) Modifications to toxic CUG RNAs induce structural stability, rescue mis-splicing in a myotonic dystrophy cell model and reduce toxicity in a myotonic dystrophy zebrafish model. Nucleic Acids Res.**
**DOI:** 10.1093/nar/gku941

---

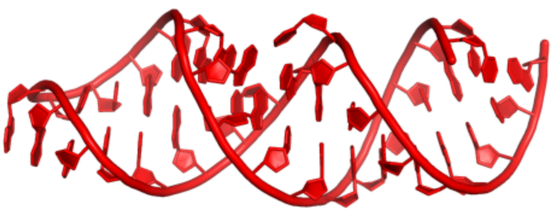

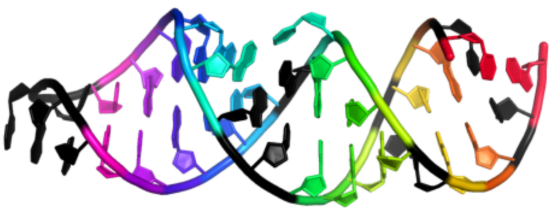

CUGCUGGCUAAGGCAUGAAAGUGCUAUGCCUGCUG
(.((.(((..((((((....))))).)))).)).)

#### 5.2 r(CCUG)

### 5.2.1 4k27_U

Myotonic Dystrophy Type 2 RNA: Structural Studies and Designed Small Molecules that Modulate RNA Function

**Release date:** 2013-11-27
**Method:** x-ray diffraction
**Resolution:** 2.35 Å
**Chain length:** 55
**Extracted length:** 55

**Total clashes:** 0
**Clashes per residue:** 0.000
**Clashscore:** 1.170

**Description:** Myotonic Dystrophy Type 2 RNA
**Organism:** nan

---

**Childs-Disney, J., Yildirim, I., Park, H., Lohman, J., Guan, L., Tran, T., Sarkar, P., Schatz, G.C., Disney, M.D. (2013) Myotonic Dystrophy Type 2 RNA: Structural Studies and Designed Small Molecules that Modulate RNA Function. ACS CHEM.BIOL.**
**DOI:** nan

---

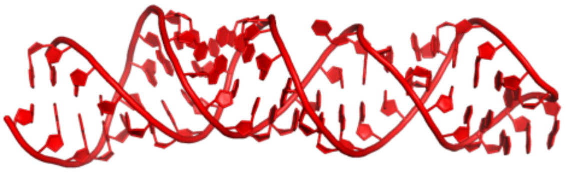

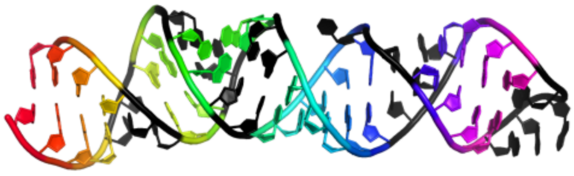

GCCCCUGCCUGCCUGCAGCUAAGGAUGAAAGUCUAUGCUGCCUGCCUGCCUGGGC
((((..((..((..(((((...((((....))))..)))))..))..))..))))

#### 5.3 r(AUUCU)

##### 5.3.1 5btm_A

Crystal structure of AUUCU repeating RNA that causes spinocerebellar ataxia type 10 (SCA10)

**Release date:** 2015-07-15
**Method:** x-ray diffraction
**Resolution:** 2.78 Å
**Chain length:** 55
**Extracted length:** 43

**Total clashes:** 6
**Clashes per residue:** 0.109
**Clashscore:** 2.200

**Description:** RNA (55-mer)
**Organism:** Homo sapiens

---

**Park, H., Gonzalez, A.L., Yildirim, I., Tran, T., Lohman, J.R., Fang, P., Guo, M., Disney, M.D. (2015) Crystallographic and Computational Analyses of AUUCU Repeating RNA That Causes Spinocerebellar Ataxia Type 10 (SCA10). Biochemistry.**
**DOI:** 10.1021/acs.biochem.5b00551

---

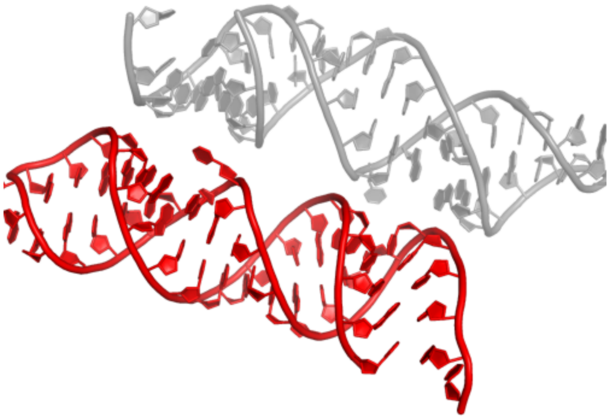

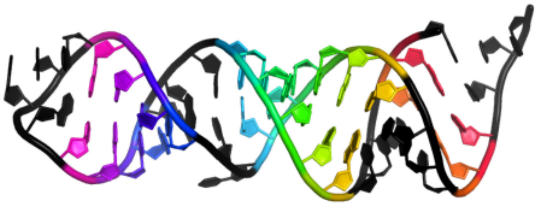

```
GUCAUUCUAUUCUAUCGGCUAAGGAUGAAAGUCUAUGCCGAUUCUAUUCUAUGGC
........((...((((((...((((....))))..))))))...))........
```

### 6 Miscellaneous (synthetic)

#### 6.1 Nanoarchitecture 1

##### 6.1.1 7jrr_A

Crystal structures of artificially designed homomeric RNA nanoarchitectures

**Release date:** 2021-09-08
**Method:** x-ray diffraction
**Resolution:** 2.16 Å
**Chain length:** 51
**Extracted length:** 51

**Total clashes:** 14
**Clashes per residue:** 0.275
**Clashscore:** 4.790

**Description:** RNA (50-MER)
**Organism:** synthetic construct

---

**Liu, D., Shao, Y., Piccirilli, J.A., Weizmann, Y. (2021) Structures of artificially designed discrete RNA nanoarchitectures at near-atomic resolution. Sci Adv.**
**DOI:** 10.1126/sciadv.abf4459

---

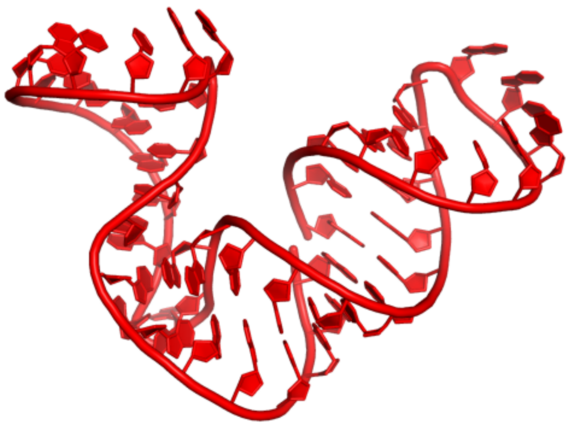

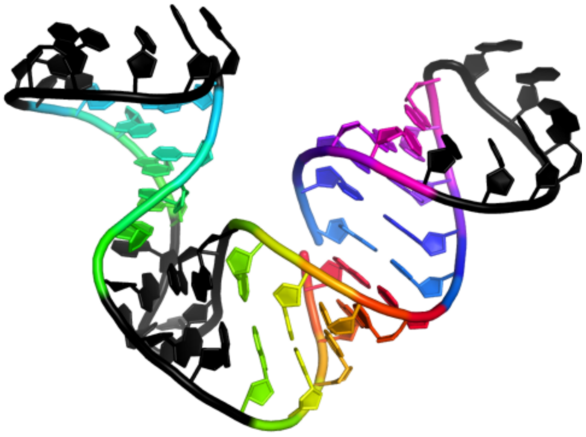

```
GGACGGGAGCUGAACCAUCCAGCGAAGAACGUCCCGACGGAUGGUUCGUCG
(((((...((((.......))))......)))))(((((.......)))))
```

#### 6.2 Nanoarchitecture 2

##### 6.2.1 7jrs_A

Crystal structures of artificially designed homomeric RNA nanoarchitectures

**Release date:** 2021-09-08
**Method:** x-ray diffraction
**Resolution:** 3.21 Å
**Chain length:** 129
**Extracted length:** 129

**Total clashes:** 19
**Clashes per residue:** 0.147
**Clashscore:** 3.950

**Description:** RNA 3D nanocage
**Organism:** synthetic construct

---

**Liu, D., Shao, Y., Piccirilli, J.A., Weizmann, Y. (2021) Structures of artificially designed discrete RNA nanoarchitectures at near-atomic resolution. Sci Adv.**
**DOI:** 10.1126/sciadv.abf4459

---

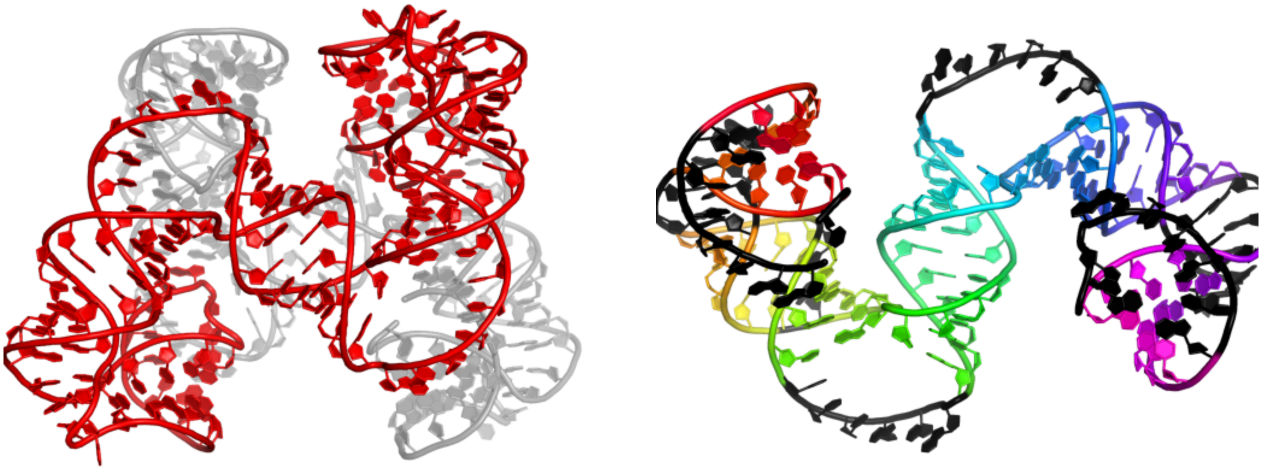

GGACGCGGUAGCGAAGAACCAUCGACUGCGACUUAGCGAGCCUGCCCUGGCAGUGGGAGCCUGAAGCAGGCACGGGCGAAGAACACUGUCAGGUCGCUGAGUCGGCGUCCCAGUUGAUGGGGAGCUACC

.......(((((......(((((.(((((((((((((((......((((((((((...(((((.........)))))......))))))))))))))))))))).......)))).)))))...))))).

#### 6.3 Synthetic hairpin

### 6.3.1 6az4_A

RNA hairpin complex with guanosine dinucleotide ligand G(5')ppp(5')G

**Release date:** 2018-02-21
**Method:** x-ray diffraction
**Resolution:** 2.98 Å
**Chain length:** 32
**Extracted length:** 21

**Total clashes:** 0
**Clashes per residue:** 0.000
**Clashscore:** 0.720

**Description:** RNA (32-MER)
**Organism:** synthetic construct

---

**Zhang, W., Tam, C.P., Zhou, L., Oh, S.S., Wang, J., Szostak, J.W. (2018) Structural Rationale for the Enhanced Catalysis of Nonenzymatic RNA Primer Extension by a Downstream Oligonucleotide. J. Am. Chem. Soc.**
**DOI:** 10.1021/jacs.7b11750

---

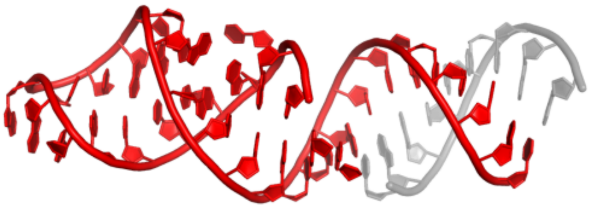

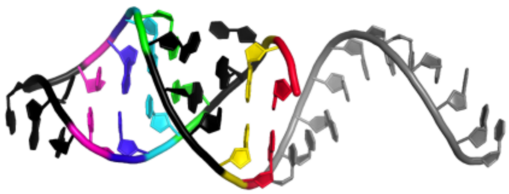

CUGCUGCUGCCGCUAAGGAUGAAAGUCUAUGC
...........((...((((....))))..))

#### 6.4 Synthetic tetraloop-tetraloop receptor

##### 6.4.1 6dvk_H

Computationally designed mini tetraloop-tetraloop receptor by the RNAMake program - construct 6 (miniTTR 6)

**Release date:** 2019-06-26
**Method:** x-ray diffraction
**Resolution:** 2.55 Å
**Chain length:** 95
**Extracted length:** 95

**Total clashes:** 0
**Clashes per residue:** 0.000
**Clashscore:** 1.980

**Description:** RNA (95-MER)
**Organism:** synthetic construct

---

**Yesselman, J.D., Eiler, D., Carlson, E.D., Gotrik, M.R., d'Aquino, A.E., Ooms, A.N., Kladwang, W., Carlson, P.D., Shi, X., Costantino, D.A., Herschlag, D., Lucks, J.B., Jewett, M.C., Kieft, J.S., Das, R. (2019) Computational design of three-dimensional RNA structure and function. Nat Nanotechnol.**
**DOI:** 10.1038/s41565-019-0517-8

---

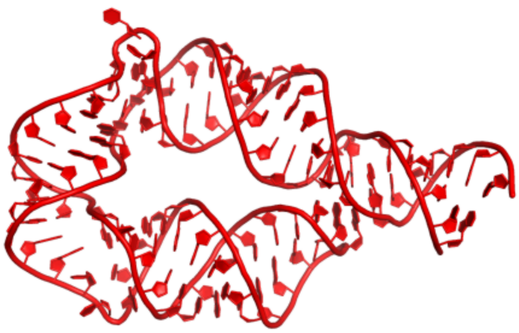

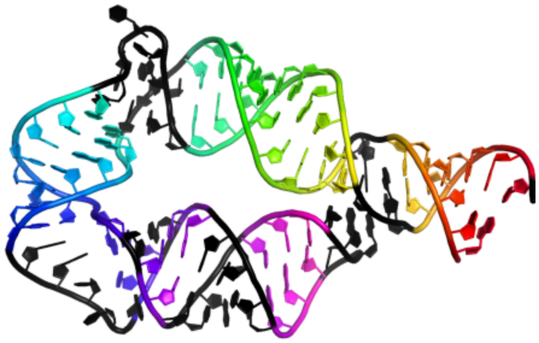

```
GUUGAUAUGGAUUUACUCCGAGGAGACGAACUACCACGAACAGGGGAAACUCUACCCGUGGCGUCUCCGUUUGACGAGUAAGUCCUAAGUCAACA
((((((..((((((((((...(((((((.....((.((....(((....)))....)).))))))))).......))))))))))...)))))).
```

#### 6.5 Synthetic G-quadruplex

##### 6.5.1 5dea_C

Crystal structure of the complex between human FMRP RGG motif and G-quadruplex RNA, cesium bound form.

**Release date:** 2015-09-23
**Method:** x-ray diffraction
**Resolution:** 2.80 Å
**Chain length:** 35
**Extracted length:** 35

**Total clashes:** 4
**Clashes per residue:** 0.114
**Clashscore:** 3.840

**Description:** sc1
**Organism:** synthetic construct

**Vasilyev, N., Polonskaia, A., Darnell, J.C., Darnell, R.B., Patel, D.J., Serganov, A. (2015) Crystal structure reveals specific recognition of a G-quadruplex RNA by a beta-turn in the RGG motif of FMRP. Proc.Natl.Acad.Sci.USA.**
**DOI:** 10.1073/pnas.1515737112

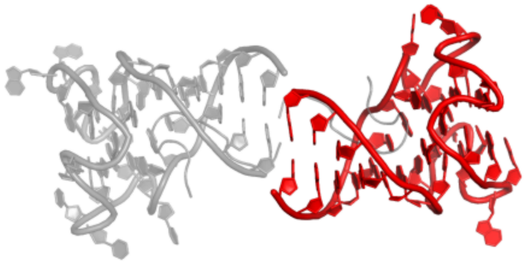

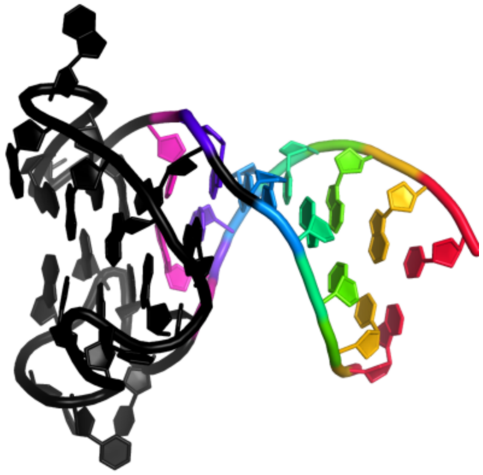

GCUGCGGUGUGGAAGGAGUGGCUGGGUUGCGCAGC
(((((.((....................)))))))

### 7 Miscellaneous

#### 7.1 ToXI

### 7.1.1 7d8o_L

Crystal structure of E. coli ToxIN type III toxin-antitoxin complex

**Release date:** 2022-01-05
**Method:** x-ray diffraction
**Resolution:** 2.10 Å
**Chain length:** 37
**Extracted length:** 37

**Total clashes:** 4
**Clashes per residue:** 0.108
**Clashscore:** 2.640

**Description:** Antitoxin RNA
**Organism:** Escherichia coli

**Manikandan, P., Sandhya, S., Nadig, K., Paul, S., Srinivasan, N., Rothweiler, U., Singh, M. (2022) Identification, functional characterization, assembly and structure of ToxIN type III toxin-antitoxin complex from E. coli. Nucleic Acids Res.**
**DOI:** 10.1093/nar/gkab1264

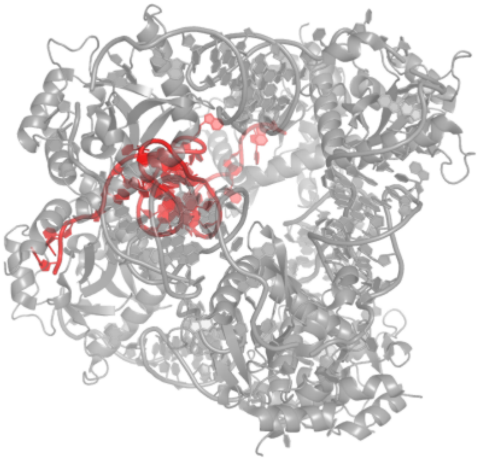

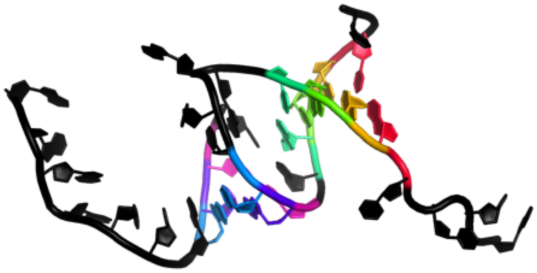

AUUUAGGUGAUUUGCUACCUUUAAGUGCAGCUAGAAA
....((((....<<<.))))......>>>........

##### 7.1.2 2xdb_G

A processed non-coding RNA regulates a bacterial antiviral system

**Release date:** 2011-01-12
**Method:** x-ray diffraction
**Resolution:** 2.55 Å
**Chain length:** 40
**Extracted length:** 36

**Total clashes:** 0
**Clashes per residue:** 0.000
**Clashscore:** 12.800

**Description:** TOXI
**Organism:** Pectobacterium atrosepticum

---

**Blower, T.R., Pei, X.Y., Short, F.L., Fineran, P.C., Humphreys, D.P., Luisi, B.F., Salmond, G.P.C. (2011) A Processed Noncoding RNA Regulates an Altruistic Bacterial Antiviral System. Nat.Struct.Mol.Biol.**
**DOI:** 10.1038/NSMB.1981

---

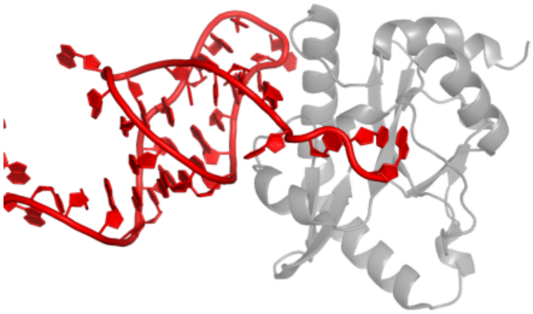

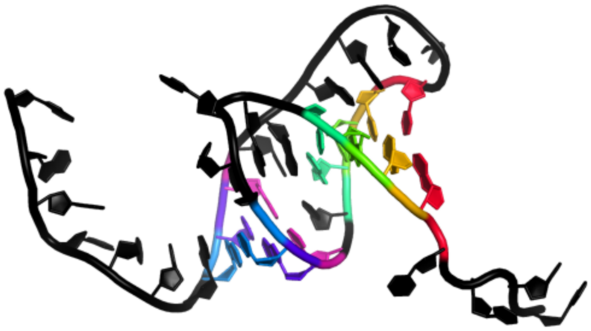

```
AUUCAGGUGAUUUGCUACCUUUAAGUGCAGCUAGAAAUUC
....((((....<<<.)))).......>>>...........
```

##### 7.1.3 4rmo_H

Crystal Structure of the CptIN Type III Toxin-Antitoxin System from Eubacterium rectale

**Release date:** 2015-09-30
**Method:** x-ray diffraction
**Resolution:** 2.20 Å
**Chain length:** 45
**Extracted length:** 45

**Total clashes:** 19
**Clashes per residue:** 0.422
**Clashscore:** 6.710

**Description:** RNA (45-MER)
**Organism:** synthetic construct

---

**Rao, F., Short, F.L., Voss, J.E., Blower, T.R., Orme, A.L., Whittaker, T.E., Luisi, B.F., Salmond, G.P. (2015) Co-evolution of quaternary organization and novel RNA tertiary interactions revealed in the crystal structure of a bacterial protein-RNA toxin-antitoxin system. Nucleic Acids Res.**
**DOI:** 10.1093/nar/gkv868

---

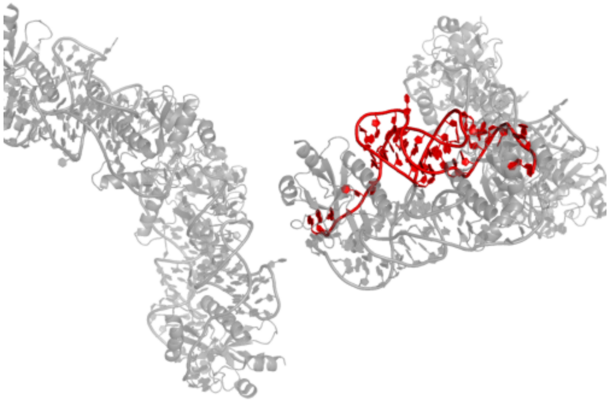

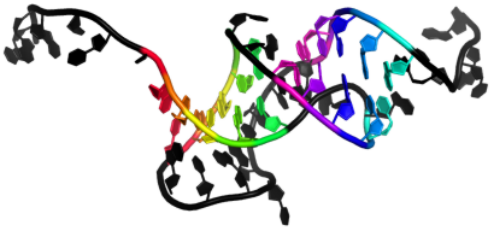

```
AAGUUUACCACUGACCGAUAUGUGGUAUAUAAAUGGUCGGGUUGA
......(((((...<<<<<..)))))..........>>>>>.....
```

#### 7.2 Structured part of acrIF8-aca2 5' UTR

### 7.2.1 8w35_C

Aca2 from Pectobacterium phage ZF40 bound to RNA

**Release date:** 2024-07-24
**Method:** electron microscopy
**Resolution:** 2.61 Å
**Chain length:** 42
**Extracted length:** 37

**Total clashes:** 0
**Clashes per residue:** 0.000
**Clashscore:** 0.620

**Description:** IR2 and IR-RBS RNA
**Organism:** Pectobacterium phage ZF40

---

**Birkholz, N., Kamata, K., Feussner, M., Wilkinson, M.E., Cuba Samaniego, C., Migur, A., Kimanius, D., Ceelen, M., Went, S.C., Usher, B., Blower, T.R., Brown, C.M., Beisel, C.L., Weinberg, Z., Fagerlund, R.D., Jackson, S.A., Fineran, P.C. (2024) Phage anti-CRISPR control by an RNA- and DNA-binding helix-turn-helix protein. Nature.**
**DOI:** 10.1038/s41586-024-07644-1

---

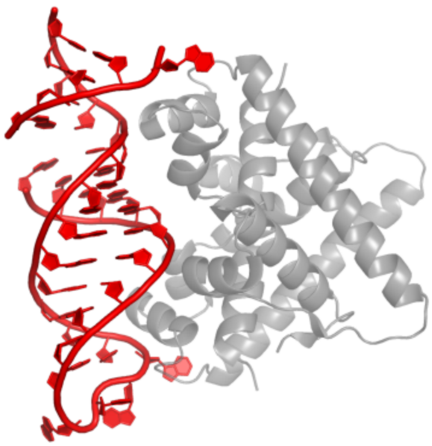

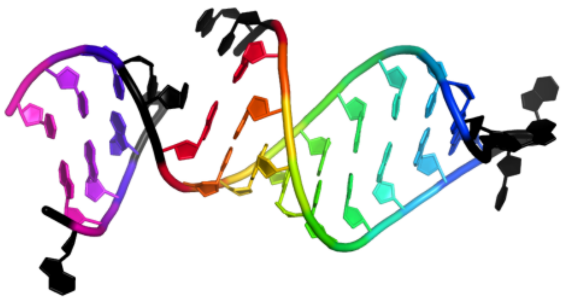

```
AUCGGUUCGAGAUGGCUCGAAUCGCUCCUAACGAGGAUUCCA
..(((((((((....)))))))))..(((....)))......
```

#### 7.3 Saguaro cactus viral mRNA structure

### 7.3.1 8t2a_R

Crystal structure of SCV PTE G18A mutant RNA in complex with Fab BL3-6

**Release date:** 2024-01-10
**Method:** x-ray diffraction
**Resolution:** 3.17 Å
**Chain length:** 90
**Extracted length:** 90

**Total clashes:** 55
**Clashes per residue:** 0.611
**Clashscore:** 9.760

**Description:** RNA (90-MER)
**Organism:** Saguaro cactus virus

---

**Ojha, M., Vogt, J., Das, N.K., Redmond, E., Singh, K., Banna, H.A., Sadat, T., Koirala, D. (2024) Structure of saguaro cactus virus 3' translational enhancer mimics 5' cap for eIF4E binding. Proc.Natl.Acad.Sci.USA.**
**DOI:** 10.1073/pnas.2313677121

---

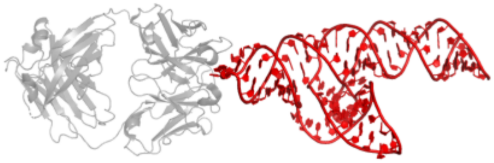

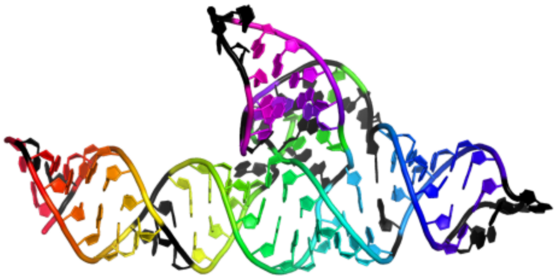

```
GGUUGCUCGACUGUGAGAGGACCUACCCACUGUGGAAACACCACAGGAACUUCCAACCUUCGGGUGGCGAGGUAGGGCAGAAGAGUGACC
 (.((((((..((((..<...(((((((.((((((.....))))))...>..((.(((....)))))).))))))))))..)))))).)
```

### 7.3.2 8t29_R

Crystal structure of SCV PTE RNA in complex with Fab BL3-6

**Release date:** 2024-01-10
**Method:** x-ray diffraction
**Resolution:** 3.13 Å
**Chain length:** 90
**Extracted length:** 90

**Total clashes:** 61
**Clashes per residue:** 0.678
**Clashscore:** 10.320

**Description:** RNA (90-MER)
**Organism:** Saguaro cactus virus

---

**Ojha, M., Vogt, J., Das, N.K., Redmond, E., Singh, K., Banna, H.A., Sadat, T., Koirala, D. (2024) Structure of saguaro cactus virus 3' translational enhancer mimics 5' cap for eIF4E binding. Proc.Natl.Acad.Sci.USA.**
**DOI:** 10.1073/pnas.2313677121

---

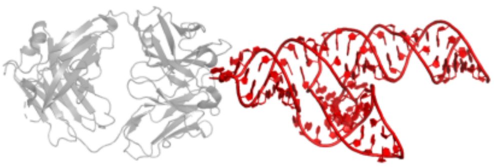

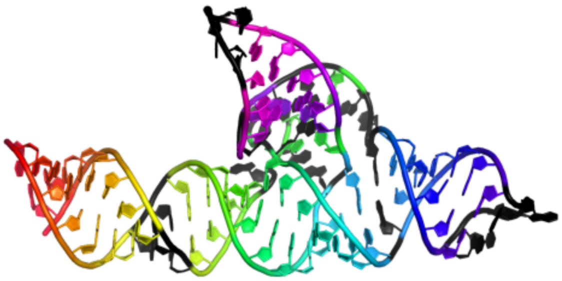

GGUUGCUCGACUGUGAGGGGACCUACCCACUGUGGAAACACCACAGGAACUUCCAACCUUCGGGUGGCGAGGUAGGGCAGAAGAGUGACC
((((((((..((((..<....(((((((.((((((.....))))))...>..((.(((....)))))).))))))))))..))))))))

### 7.3.3 8t2b_R

Crystal structure of SCV PTE G18C mutant RNA in complex with Fab BL3-6

**Release date:** 2024-01-10
**Method:** x-ray diffraction
**Resolution:** 3.18 Å
**Chain length:** 90
**Extracted length:** 90

**Total clashes:** 49
**Clashes per residue:** 0.544
**Clashscore:** 9.440

**Description:** RNA (90-MER)
**Organism:** Saguaro cactus virus

**Ojha, M., Vogt, J., Das, N.K., Redmond, E., Singh, K., Banna, H.A., Sadat, T., Koirala, D. (2024) Structure of saguaro cactus virus 3' translational enhancer mimics 5' cap for eIF4E binding. Proc.Natl.Acad.Sci.USA.**
**DOI:** 10.1073/pnas.2313677121

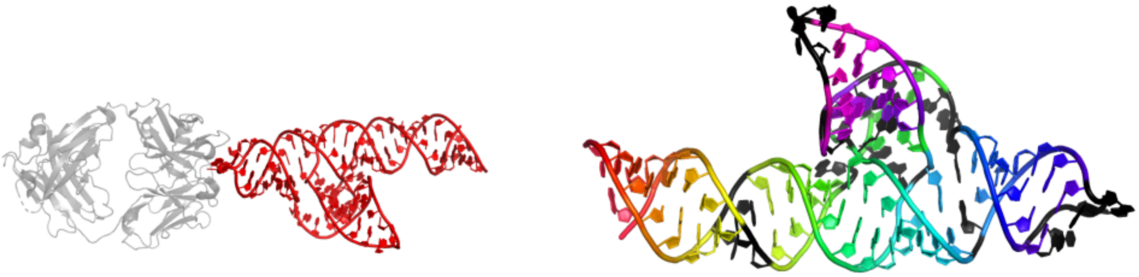

GGUUGCUCGACUGUGAGCGGACCUACCCACUGUGGAAACACCACAGGAACUUCCAACCUUCGGGUGGCGAGGUAGGGCAGAAGAGUGACC
((((((((..((((..<...(((((((.((((((.....))))))...>..((.(((....))))).))))))))))..))))))))

#### 7.4 ITS-2

### 7.4.1 7r6q_6

State E2 nucleolar 60S ribosome biogenesis intermediate - Foot region model

**Release date:** 2022-11-09
**Method:** electron microscopy
**Resolution:** 2.98 Å
**Chain length:** 87
**Extracted length:** 87

**Total clashes:** 0
**Clashes per residue:** 0.000
**Clashscore:** 6.420

**Description:** ITS-2
**Organism:** Saccharomyces cerevisiae BY4741

**Cruz, V.E., Sekulski, K., Peddada, N., Sailer, C., Balasubramanian, S., Weirich, C.S., Stengel, F., Erzberger, J.P. (2022) Sequence-specific remodeling of a topologically complex RNP substrate by Spb4. Nat.Struct.Mol.Biol.**
**DOI:** 10.1038/s41594-022-00874-9

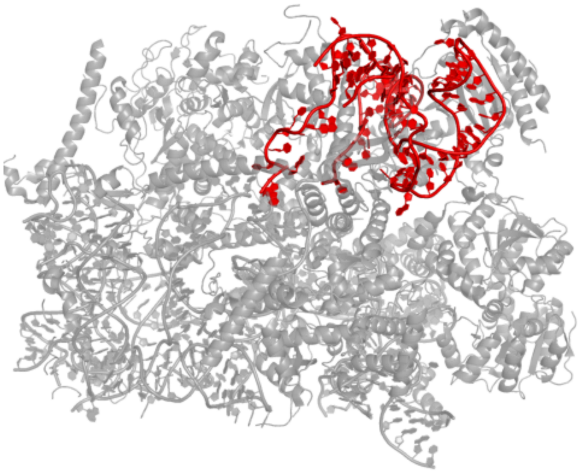

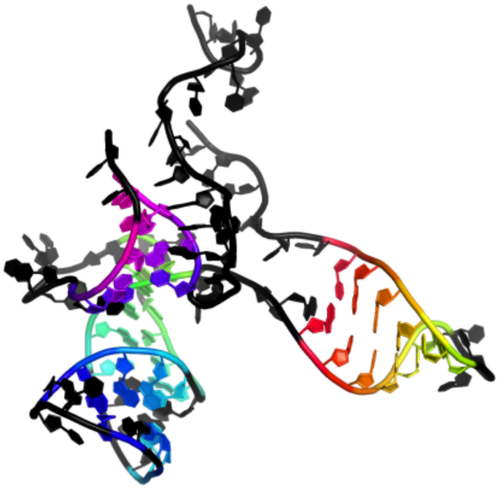

```
CCUUCUCAAACAUUCUGUUUGGUAGUGAGUGAUACUCUUUGGAGUUAACUUGAAAUUGCUGGCCUUUAGGCGAACAAUGUUCUUAAA
......((((((...))))))....(((((...(((.(...))))..))))..).......((((..))))................
```

### 7.4.2 7u0h_6

State NE1 nucleolar 60S ribosome biogenesis intermediate - Overall model

**Release date:** 2022-12-14
**Method:** electron microscopy
**Resolution:** 2.76 Å
**Chain length:** 87
**Extracted length:** 87

**Total clashes:** 60
**Clashes per residue:** 0.690
**Clashscore:** 7.920

**Description:** ITS2 rRNA
**Organism:** Saccharomyces cerevisiae BY4741

**Cruz, V.E., Sekulski, K., Peddada, N., Sailer, C., Balasubramanian, S., Weirich, C.S., Stengel, F., Erzberger, J.P. (2022) Sequence-specific remodeling of a topologically complex RNP substrate by Spb4. Nat.Struct.Mol.Biol.**
**DOI:** 10.1038/s41594-022-00874-9

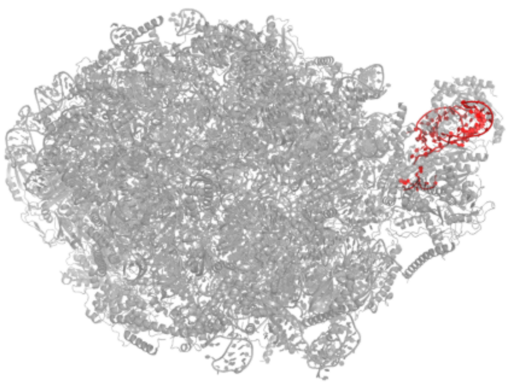

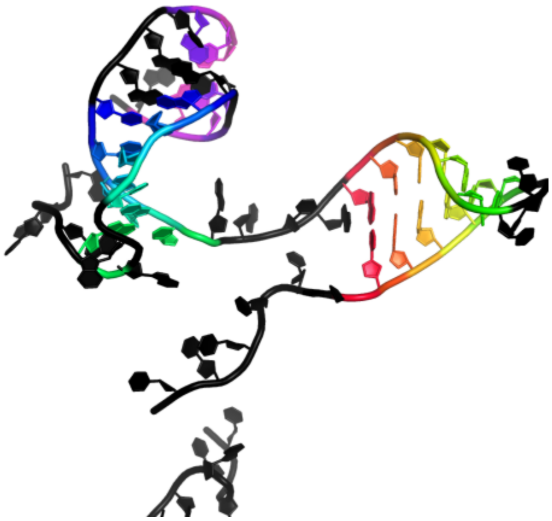

```
CCUUCUCAAACAUUCUGUUUGGUAGUGAGUGAUACUCUUUGGAGUUAACUUGAAAUUGCUGGCCUUUAGGCGAACAAUGUUCUUAAA
......((((((...))))))....(((((...(((......)))..))))..)..................................
```

#### 7.5 ENE

##### 7.5.1 7ljy_B

Cryo-EM structure of the B dENE construct complexed with a 28-mer poly(A)

**Release date:** 2021-04-14
**Method:** electron microscopy
**Resolution:** 5.60 Å
**Chain length:** 76
**Extracted length:** 76

**Total clashes:** 5
**Clashes per residue:** 0.066
**Clashscore:** 1.860

**Description:** B dENE construct
**Organism:** Oryza sativa

---

**Torabi, S.F., Chen, Y.L., Zhang, K., Wang, J., DeGregorio, S.J., Vaidya, A.T., Su, Z., Pabit, S.A., Chiu, W., Pollack, L., Steitz, J.A. (2021) Structural analyses of an RNA stability element interacting with poly(A). Proc.Natl.Acad.Sci.USA.**
**DOI:** 10.1073/pnas.2026656118

---

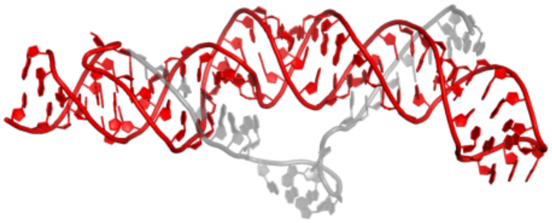

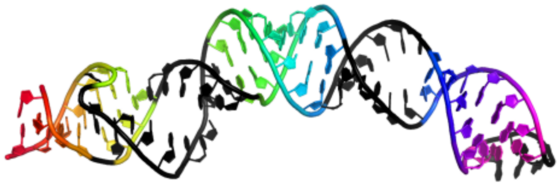

GGGUACUCUUUUCUUUGUCAUGGUUUUCUCAGGCGAAAGUCUGAGUUUUUACAUGACAAAGUUUUUAACGAGGCCC
((((.(((......((((((((.....(((((((....)))))))......))))))))..........)))))))

### 7.5.2 3p22_A

Crystal structure of the ENE, a viral RNA stability element, in complex with A9 RNA

**Release date:** 2010-12-08
**Method:** x-ray diffraction
**Resolution:** 2.50 Å
**Chain length:** 40
**Extracted length:** 40

**Total clashes:** 3
**Clashes per residue:** 0.075
**Clashscore:** 3.530

**Description:** Core ENE hairpin from KSHV PAN RNA
**Organism:** nan

**Mitton-Fry, R.M., DeGregorio, S.J., Wang, J., Steitz, T.A., Steitz, J.A. (2010) Poly(A) tail recognition by a viral RNA element through assembly of a triple helix. Science.**
**DOI:** 10.1126/science.1195858

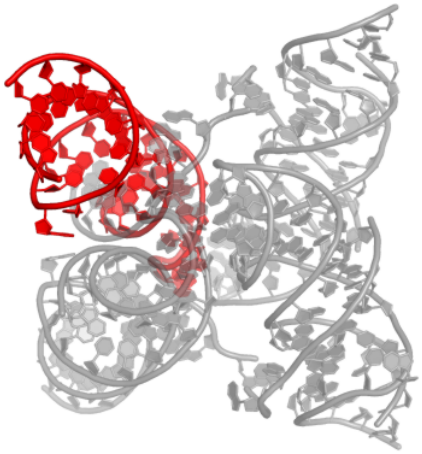

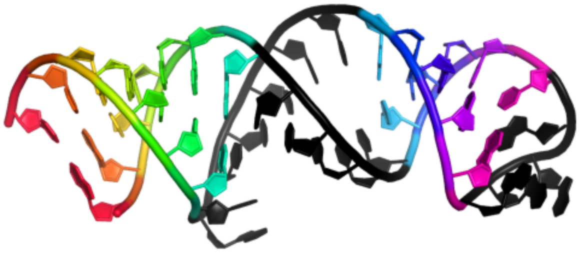

GGCUGGGUUUUUCCUUCGAAAGAAGGUUUUUAUCCCAGUC
((((((.....(((((....)))))......))))))

##### 7.5.3 7jnh_B

Crystal structure of a double-ENE RNA stability element in complex with a 28-mer poly(A) RNA

**Release date:** 2021-01-20
**Method:** x-ray diffraction
**Resolution:** 2.89 Å
**Chain length:** 86
**Extracted length:** 86

**Total clashes:** 62
**Clashes per residue:** 0.721
**Clashscore:** 13.540

**Description:** Core double ENE RNA (Xtal construct) from Oryza sativa transposon,Core double ENE RNA (Xtal construct) from Oryza sativa transposon
**Organism:** Oryza sativa

**Torabi, S.F., Vaidya, A.T., Tycowski, K.T., DeGregorio, S.J., Wang, J., Shu, M.D., Steitz, T.A., Steitz, J.A. (2021) RNA stabilization by a poly(A) tail 3'-end binding pocket and other modes of poly(A)-RNA interaction. Science.**

**DOI:** 10.1126/science.abe6523

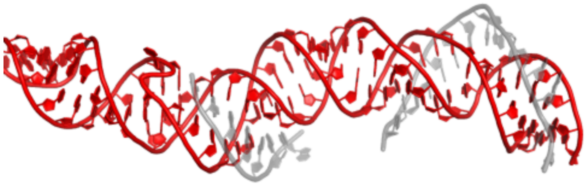

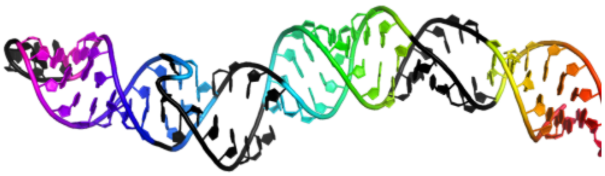

GGGCUGAGUUUUUACAUGACAAAGUUUUUAACGAGGCAGCGGCGAAAGUCGCUGUACUCUUUUCUUUGUCAUGGUUUUCUCAGCCC
((((((((......((((((((((........(((((((((((....)))))))).)))....))))))))))).....))))))))

### 7.5.4 3p22_E

Crystal structure of the ENE, a viral RNA stability element, in complex with A9 RNA

**Release date:** 2010-12-08
**Method:** x-ray diffraction
**Resolution:** 2.50 Å
**Chain length:** 40
**Extracted length:** 40

**Total clashes:** 8
**Clashes per residue:** 0.200
**Clashscore:** 3.530

**Description:** Core ENE hairpin from KSHV PAN RNA
**Organism:** nan

**Mitton-Fry, R.M., DeGregorio, S.J., Wang, J., Steitz, T.A., Steitz, J.A. (2010) Poly(A) tail recognition by a viral RNA element through assembly of a triple helix. Science.**
**DOI:** 10.1126/science.1195858

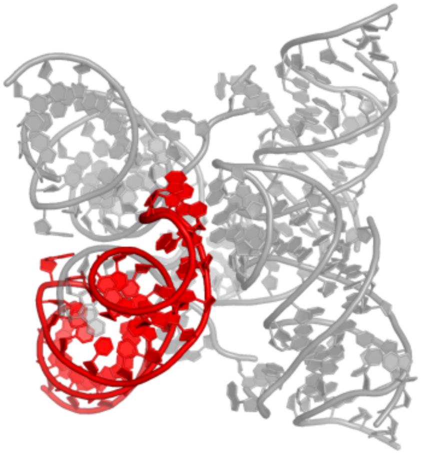

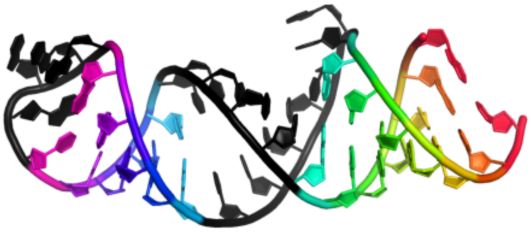

GGCUGGGUUUUUCCUUCGAAAGAAGGUUUUUAUCCCAGUC
((((((.....(((((....))))).......))))))

#### 7.6 ASH1 mRNA E3-localization element

### 7.6.1 5m0h_A

Crystal structure of the central flexible region of ASH1 mRNA E3-localization element

**Release date:** 2017-01-18
**Method:** x-ray diffraction
**Resolution:** 2.65 Å
**Chain length:** 42
**Extracted length:** 42

**Total clashes:** 10
**Clashes per residue:** 0.238
**Clashscore:** 3.700

**Description:** ASH1 E3 (42 nt-TL/TLR)
**Organism:** Saccharomyces cerevisiae

**Edelmann, F.T., Schlundt, A., Heym, R.G., Jenner, A., Niedner-Boblenz, A., Syed, M.I., Paillart, J.C., Stehle, R., Janowski, R., Sattler, M., Jansen, R.P., Niessing, D. (2017) Molecular architecture and dynamics of ASH1 mRNA recognition by its mRNA-transport complex. Nat. Struct. Mol. Biol.**
**DOI:** 10.1038/nsmb.3351

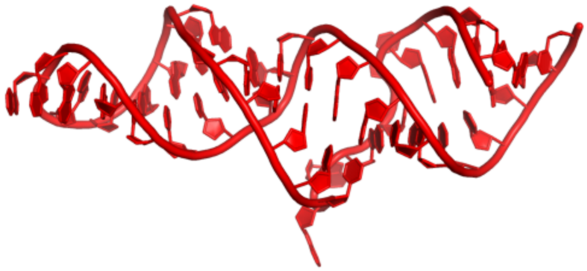

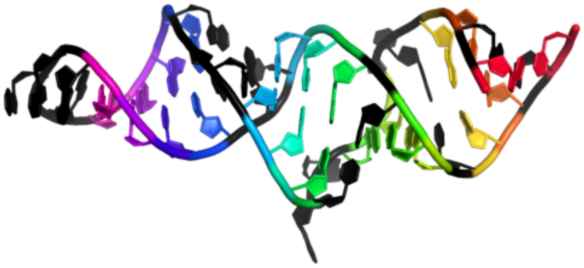

GAUAACUGAAUCGCUAAGGAUGAAAGUCUAUGCGACAUUAUC
(.((..((...(((...((((....))))..))).)).)).)

#### 7.7 SCNMV xrRNA

### 7.7.1 6d3p_A

Crystal structure of an exoribonuclease-resistant RNA from Sweet clover necrotic mosaic virus (SCNMV)

**Release date:** 2018-06-20
**Method:** x-ray diffraction
**Resolution:** 2.90 Å
**Chain length:** 45
**Extracted length:** 45

**Total clashes:** 18
**Clashes per residue:** 0.400
**Clashscore:** 6.560

**Description:** RNA (45-MER)
**Organism:** Sweet clover necrotic mosaic virus

**Steckelberg, A.L., Akiyama, B.M., Costantino, D.A., Sit, T.L., Nix, J.C., Kieft, J.S. (2018) A folded viral noncoding RNA blocks host cell exoribonucleases through a conformationally dynamic RNA structure. Proc. Natl. Acad. Sci. U.S.A.**
**DOI:** 10.1073/pnas.1802429115

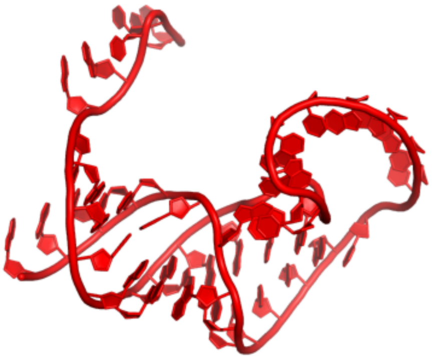

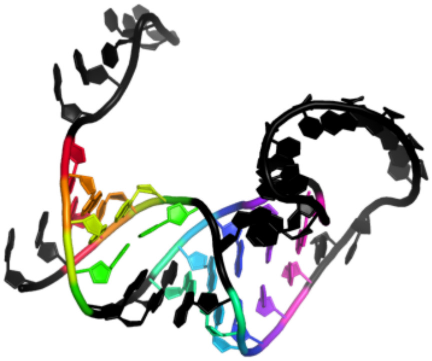

```
GGGCGUAACCUCCAUCCGAGUUGCAAGAGAGGGAAACGCAGUCUC
..((((..(((((.............).))))...))))......
```

#### 7.8 Influenza B vRNA promoter

### 7.8.1 6t2c_V

Bat Influenza A polymerase recycling complex

**Release date:** 2020-04-15
**Method:** electron microscopy
**Resolution:** 3.52 Å
**Chain length:** 34
**Extracted length:** 34

**Total clashes:** 0
**Clashes per residue:** 0.000
**Clashscore:** 4.190

**Description:** vRNA
**Organism:** Influenza B virus

**Wandzik, J.M., Kouba, T., Karuppasamy, M., Pflug, A., Drncova, P., Provaznik, J., Azevedo, N., Cusack, S. (2020) A Structure-Based Model for the Complete Transcription Cycle of Influenza Polymerase. Cell.**
**DOI:** 10.1016/j.cell.2020.03.061

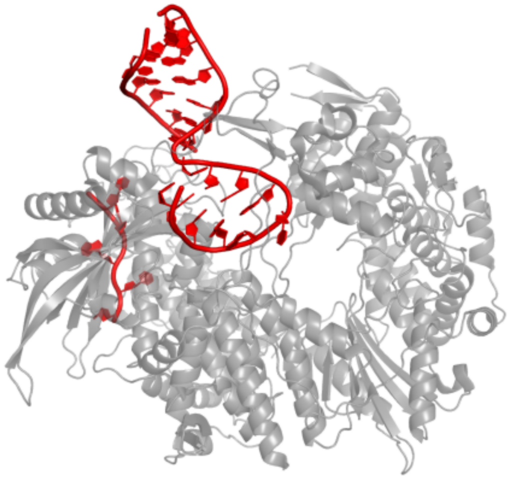

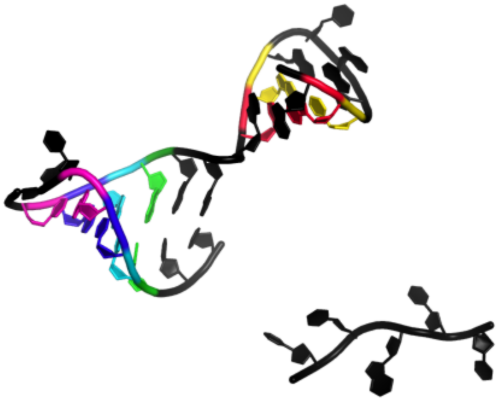

```
AGUAGUAACAAGAGGUAUUACCUCUGCUUCUGCU
.((....))...((((...))))...........
```

#### 7.9 NAD-II riboswitch

### 7.9.1 8hb8_A

Crystal structure of NAD-II riboswitch (single strand) with NMN

**Release date:** 2023-03-22
**Method:** x-ray diffraction
**Resolution:** 2.30 Å
**Chain length:** 55
**Extracted length:** 55

**Total clashes:** 8
**Clashes per residue:** 0.145
**Clashscore:** 2.180

**Description:** RNA (55-MER)
**Organism:** Streptococcus parasanguinis

---

**Peng, X., Liao, W., Lin, X., Lilley, D.M.J., Huang, L. (2023) Crystal structures of the NAD+-II riboswitch reveal two distinct ligand-binding pockets. Nucleic Acids Res.**
**DOI:** 10.1093/nar/gkad102

---

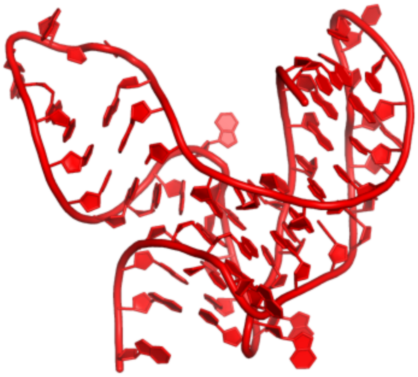

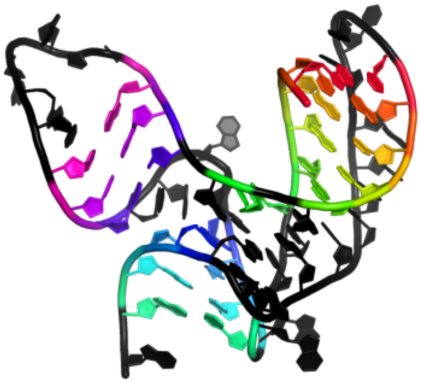

```
GCGGCGUUGCGUCCGAAAGUCUAAACAGACACGGCCGCUUAAAAACAAAAGGAGA
((((((...<.<<<....(((......))).).))))))............>>>>.
```

##### 7.9.2 8hba_A

Crystal structure of NAD-II riboswitch (single strand) with NAD

**Release date:** 2023-03-22
**Method:** x-ray diffraction
**Resolution:** 2.64 Å
**Chain length:** 56
**Extracted length:** 56

**Total clashes:** 53
**Clashes per residue:** 0.946
**Clashscore:** 15.750

**Description:** RNA (55-MER)
**Organism:** Streptococcus parasanguinis

---

**Peng, X., Liao, W., Lin, X., Lilley, D.M.J., Huang, L. (2023) Crystal structures of the NAD+-II riboswitch reveal two distinct ligand-binding pockets. Nucleic Acids Res.**
**DOI:** 10.1093/nar/gkad102

---

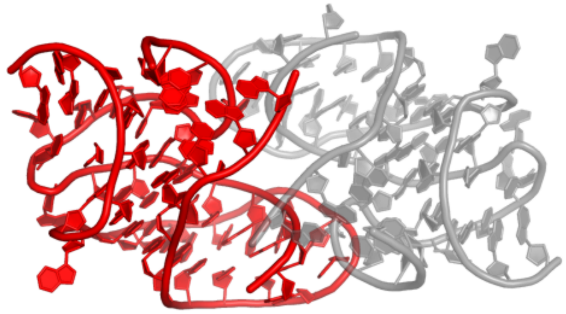

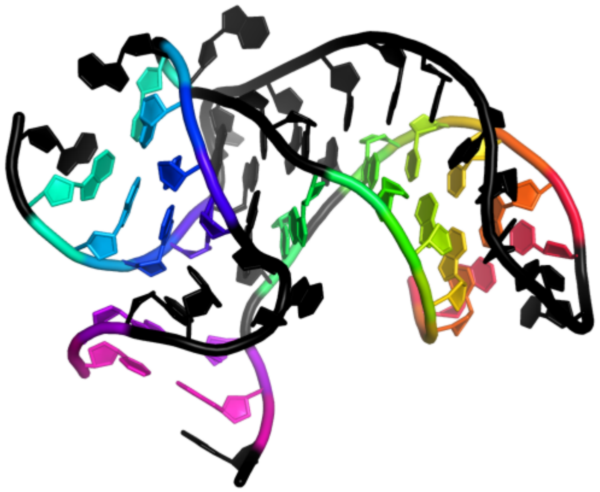

```
CGCGGCGUUGCGUCCGAAAGUCUAAACAGACACGGCCGCUUAAAAACAAAAGGAGA
.((((((...<.<<<....((........)).).)))))............>>>>.
```
